# Nuclear phylogenomics of grasses (Poaceae) supports current classification and reveals repeated reticulation

**DOI:** 10.1101/2024.05.28.596153

**Authors:** Grass Phylogeny Working Group III, Watchara Arthan, William J. Baker, Matthew D. Barrett, Russell L. Barrett, Jeffrey Bennetzen, Guillaume Besnard, Matheus E. Bianconi, Joanne L. Birch, Pilar Catalán, Wenli Chen, Maarten Christenhusz, Pascal-Antoine Christin, Lynn G. Clark, J. Travis Columbus, Charlotte Couch, Darren M. Crayn, Gerrit Davidse, Soejatmi Dransfield, Luke T. Dunning, Melvin R. Duvall, Sarah Z. Ficinski, Amanda E. Fisher, Siri Fjellheim, Felix Forest, Lynn J. Gillespie, Jan Hackel, Thomas Haevermans, Trevor R. Hodkinson, Chien-Hsun Huang, Weichen Huang, Aelys M. Humphreys, Richard W. Jobson, Canisius J. Kayombo, Elizabeth A. Kellogg, John M. Kimeu, Isabel Larridon, Rokiman Letsara, De-Zhu Li, Jing-Xia Liu, Ximena Londoño, Quentin W.R. Luke, Hong Ma, Terry D. Macfarlane, Olivier Maurin, Michael R. McKain, Todd G.B. McLay, Maria Fernanda Moreno-Aguilar, Daniel J. Murphy, Olinirina P. Nanjarisoa, Guy E. Onjalalaina, Paul M. Peterson, Rivontsoa A. Rakotonasolo, Jacqueline Razanatsoa, Jeffery M. Saarela, Lalita Simpson, Neil W. Snow, Robert J. Soreng, Marc Sosef, John J.E. Thompson, Paweena Traiperm, G. Anthony Verboom, Maria S. Vorontsova, Neville G. Walsh, Jacob D. Washburn, Teera Watcharamongkol, Michelle Waycott, Cassiano A.D. Welker, Martin D. Xanthos, Nianhe Xia, Lin Zhang, Alexander Zizka, Fernando O. Zuloaga, Alexandre R. Zuntini

## Abstract

- Grasses (Poaceae) comprise around 11,800 species and are central for human livelihoods and terrestrial ecosystems. Knowing their relationships and evolutionary history is key to comparative research and crop breeding. Advances in genome-scale sequencing allow for increased breadth and depth of phylogenomic analyses, making it possible to infer a new reference species tree of the family.
- We inferred a comprehensive species tree of grasses by combining new and published sequences for 331 nuclear genes from genome, transcriptome, target enrichment and shotgun data. Our 1,153-tip tree covers 79% of grass genera (including 21 genera sequenced for the first time) and all but two small tribes. We compared it to a 910-tip plastome tree.
- The nuclear phylogeny matches that of the plastome at most deep branches, with only a few instances of incongruence. Gene tree–species tree reconciliation suggests that reticulation events occurred repeatedly in the history of grasses.
- We provide a robust framework for the grass tree of life to support research on grass evolution, including modes of reticulation, and genetic diversity for sustainable agriculture.

## Introduction

With almost 11,800 species in 791 genera (Soreng *et al*., 2022), grasses (Poaceae) are among the largest plant families and one of the most important for humans. Grasses include the primary food crops rice, maize and wheat, sources of fibre and building materials such as reed and bamboo, and biofuel crops such as sugarcane and switchgrass. Much of the global land surface is covered by grass-dominated ecosystems, where grasses impact productivity, nutrient cycling and vegetation structure by mediating fire and herbivory (Edwards *et al*., 2010; Bond, 2016). Grasses are also overrepresented among the world’s most damaging agricultural weeds (Holm *et al*., 1977) and invasive plants (Linder *et al*., 2018). Understanding functional diversification, adaptation and novel crop breeding in this important plant group requires a solid understanding of its evolutionary relationships.

Efforts to uncover the phylogenetic history of grasses have tracked the development of new technology and analytical tools, beginning with cladistic analysis of morphology (e.g., Campbell & Kellogg, 1987). Almost as soon as nucleotide sequencing became possible, it was used to investigate grasses (rRNA sequencing, Hamby & Zimmer, 1988, and chloroplast DNA, Clark *et al*., 1995), and the results interpreted in the light of known morphology and classification. Hundreds of papers have been published since using nucleic acids, most recently DNA, to assess grass phylogeny at all taxonomic levels and assembling information from all three genomes in the cell (plastid, mitochondrial, and nuclear). These efforts have been punctuated by two major phylogenetic analyses, Grass Phylogeny Working Group I (GPWG, 2001) and GPWG II (2012), which were used to generate updated classifications of the family (Kellogg, 2015; Soreng *et al*., 2022).

The major outlines of grass phylogeny have now been known for several decades and corroborated by accumulating data, with major lineages recognised as subfamilies (Kellogg, 2015; Soreng *et al*., 2022). The earliest divergences in the grass family gave rise to three successive lineages, Anomochlooideae, Pharoideae, and Puelioideae, each comprising just a few species. After the divergence of those three, however, the remaining grasses gave rise to two sister lineages, known as BOP and PACMAD, each of which became a species-rich clade with several robust subclades. This sturdy phylogenetic framework is reflected in a strong subfamilial classification, with subfamilies divided into equally robust tribes. Attention in recent years has largely shifted to relationships of tribes, subtribes, and genera.

Reticulate evolution is common in the grasses. Allopolyploidy is widespread in the family, particularly among closely related species and genera, with as many as 80% of species estimated to be of polyploid origin (Stebbins, 1985). The textbook example is bread wheat (*Triticum aestivum*) and its ruderal annual ancestors, the history of which was determined in the first part of the 20th century using cytogenetic tools (Kihara, 1982; Tsunewaki, 2018). Nucleotide sequence data have verified the hybrid origin of wheat and gone on to show that reticulate evolution is the norm in the entire tribe Triticeae (Feldman & Levy, 2023, and references therein). We have also learned that three of the four major clades of Bambusoideae are of allopolyploid origin (Triplett *et al*., 2014; Guo *et al*., 2019; Chalopin *et al*., 2021; Ma *et al*., 2024), as are at least one third of the species in Andropogoneae (Estep *et al*., 2014). Large-scale lateral gene transfer has also been demonstrated in *Alloteropsis semialata* (Dunning *et al*., 2019) and for a number of genomes across the family (Hibdige *et al*., 2021), although it remains unclear how common such genetic exchanges are. Network-like reticulations are therefore expected throughout Poaceae, but there is no indication so far that they obscure a dominant tree-like phylogeny.

Data relevant to grass phylogeny continue to accumulate in the genomic era, but in an uneven pattern. Major recent studies have inferred family trees based on the plastid genome (Saarela *et al*., 2018; Gallaher *et al*., 2022; Hu *et al*., 2023) or large parts of the nuclear genome (Huang *et al*., 2022). In addition, a wealth of full-genome assemblies is now available for grasses, mainly for groups that have been studied intensively, such as major crops and their congeners including rice (Wang & Han, 2022), maize (Hufford *et al*., 2021), wheat (Walkowiak *et al*., 2020) and sugarcane (Healey *et al*., 2024), among many others. At the same time, some genera and many species remain virtually unknown beyond a scientific name and general morphology. While the poorly known taxa may be represented in major herbaria, fresh material can be hard to obtain, weakening attempts to fully sample the grass tree of life with phylogenomic technologies.

Fortunately, we are now experiencing the confluence of: 1) global sources of diversity data including plant specimens held in herbaria worldwide, 2) widespread use of short-read sequencing that can accommodate even fragmented DNA, 3) analytical tools for assembling and interpreting massive amounts of sequence data, and 4) technical tools for efficient sequencing, such as target capture. For example, the development of a universal probe set for flowering plants, Angiosperms353 (Johnson *et al*., 2019; Baker *et al*., 2021), has enabled initiatives to sequence all angiosperm plant genera (Baker *et al*., 2022; Zuntini *et al*., 2024) or entire continental floras such as that of Australia (https://www.genomicsforaustralianplants.com/). It became apparent that an updated synthesis of existing and new data for grasses, similar to the previous Grass Phylogeny Working Group efforts (GPWG, 2001; GPWG II 2012), would be timely and make possible a phylogeny that incorporates representatives of most of the 791 genera of the family using genome-scale data. In the process, we will gain a broader understanding of the frequency and impact of reticulation.

Accordingly, here we present the most comprehensive nuclear phylogenomic tree of the grass family to date. Via a large community effort, we maximised taxon sampling by combining whole-genome, transcriptome, target capture and shotgun datasets. Based on the Angiosperms353 gene set, we inferred a nuclear multigene species tree using a coalescent-based method that uses information from multicopy gene trees. We also inferred a plastome tree and tested for incongruence between plastome and nuclear trees. Finally, we used gene tree–species tree reconciliation analyses to explore the signal for reticulation in the nuclear data.

## Material and Methods

### Datasets and species sampling

Drawing from a combined effort of the Poaceae research community, we leveraged five diverse sets of genomic data (see full accession table in the data repository, https://doi.org/10.5281/zenodo.10996136). We deployed a set of automated filters and repeated expert input from the group to remove duplicates, samples with insufficient data, and potentially misidentified accessions. The final set of accessions included:

1. 450 Illumina target capture read accessions enriched with the Angiosperms353 probe set (Johnson *et al*., 2019), generated as part of the ‘Genomics for Australian Plants’ (GAP) and ‘Plant and Fungal Trees of Life’ (PAFTOL; Baker *et al*., 2022) initiatives as well as a project focused on Loliinae grasses (P. Catalán et al. in prep.). Sampling focused on genera without existing nuclear or plastome genomic data.
2. 295 Illumina shotgun, whole-genome sequencing accessions, of which 204 are ‘genome skims’ with a sequencing depth <5× estimated for our target gene set (see below). Of these shotgun accessions, many had been used in previous studies for the assembly of plastid genomes (see metadata table).
3. 17 Illumina target capture read accessions enriched in 122 nuclear loci (different from Angiosperms353) that were previously used in a phylogenetic study of the subfamily Chloridoideae (Fisher *et al*., 2016). These are treated here like the shotgun datasets.
4. 343 assembled transcriptomes from two recent Poaceae studies (331 samples; Huang *et al*., 2022; Zhang *et al*., 2022) and the 1KP initiative (12 samples, One Thousand Plant Transcriptomes Initiative, 2019).
5. 48 assembled and annotated genome sequences from Phytozome v.13, Ensembl Plants, or other sources.

Angiosperms353 target capture data were generated by the PAFTOL project following the protocols of Baker *et al*. (2022). Methods varied for the other contributed datasets (details in accession table and Supplementary Methods). Leaves were sampled mostly from herbarium specimens, although silica dried material was used in some cases. Sampling was iteratively refined using expert input from the working group to remove accessions with unclear identity and duplicates per species (retaining the highest-coverage accession, i.e. genome > transcriptome > target capture > shotgun). Species names were harmonised using the World Checklist of Vascular Plants (Govaerts *et al*., 2021) as well as expertise from our working group.

### Grass-specific Angiosperms353 reference dataset

Prior to sequence assembly from target capture and shotgun datasets, we produced a Poaceae-specific set of reference Angiosperms353 sequences to improve recovery and account for grass-wide gene duplications. This grass-specific reference dataset consists of coding sequences (CDS) extracted from published genomes and transcriptomes of 60 species, representing seven of the 12 grass subfamilies and including an available genome sequence from the sister group Ecdeiocoleaceae–Joinvilleaceae (*Joinvillea ascendens*). First, CDS of the Angiosperms353 homologs were extracted from the reference genomes and transcriptomes using the tblastn tool of BLAST+ v.2.2.29 (Camacho *et al*., 2009), with the original Angiosperms353 probe set used as protein queries (e-value ≤ 10^-3^). To reduce false positives, only hits with alignments >65% of the query length and sequence identity >60% were retained. This filtered homolog set was then sorted into orthogroups using Orthofinder v.2.5.2 (Emms & Kelly, 2019), with the MSA mode using MAFFT v.7.481 (Katoh & Standley, 2013) as the sequence aligner, and FastTree v.2.1.11 (Price *et al*., 2010) to generate gene trees, using default parameters in each case.

Using the phylogenetic hierarchical method of Orthofinder, we extracted orthogroups at the level of the most recent common ancestor of the BOP–PACMAD clade, the crown group which covers >99% of grass species and most available reference genomes. Two of the original Angiosperms353 markers (g5422 and g6924) were not detected in any of the reference genomes or transcriptomes and were therefore not used. Five other markers were duplicated before the BOP–PACMAD split (g4527, g5434, g5945, g5950 and g7024); these duplicates were therefore treated as separate markers in our analyses. For these five duplicated genes, homologs of the three reference samples representing subfamily Anomochlooideae, sister to all other Poaceae, and the outgroup Joinvilleaceae were subsequently added to each of the two corresponding orthogroups.

This initial reference dataset was then curated to remove non-homologous sequences and potential pseudogenes (see Supplementary Methods). The final reference dataset consisted of 356 orthogroups, and encompassed all homologous sequences of the 60 reference species, including paralogs from lineage-specific duplications within each orthogroup. Note that three of the markers (g5328, g5922 and g6128) were removed prior to phylogenetic analysis on the basis that they contained regions of low complexity in their sequences, which resulted in low quality assemblies (see below) as revealed by preliminary analyses.

### Angiosperms353 sequence assembly

The orthogroup dataset was used as a reference for sequence assembly using HybPiper v.1.3.1 (Johnson *et al*., 2016). Illumina reads were initially trimmed using Trimmomatic v.0.38 (Bolger *et al*., 2014) to remove adapters, low quality bases and short reads (SLIDINGWINDOW:4:20, MINLEN:40). Sequences were assembled using the Burrows-Wheeler Alignment tool (BWA, Li & Durbin, 2009) with default parameters, except the coverage cut-off level, which was reduced to 4× for the target capture datasets, and to 1× for shotgun accessions due to the low sequencing depth of a subset of samples. Given the low number of markers recovered for most shotgun accessions, we used a custom assembly strategy optimised for the assembly of sequences from low coverage datasets (explained below). When a sequence was assembled by both HybPiper and the custom method, only the longest assembly was retained.

The custom assembly strategy consisted of a mapping-consensus pipeline modified from Olofsson *et al*. (2019) and Bianconi *et al*. (2020) to support the assembly of paralogs (Fig. S1). First, filtered reads were mapped to the orthogroup reference dataset using Bowtie2 v.2.5.3 (Langmead & Salzberg, 2012) with the sensitive-local mode and reporting all alignments. Then, for each orthogroup, the reference sequence with the most bases covered was identified and included along with its paralogs (i.e. homeologs or paralogs from lineage-specific duplications) in a second, accession-specific reference dataset. This reduced the reference dataset to a single species per orthogroup, which allowed subsequent read mapping refinement, and simplified downstream processing. Read mapping was then repeated on this accession-specific reference using the parameters described above, and the resulting read alignments were converted into majority consensus sequences using Samtools v.1.19.2 (Li *et al*., 2009); *consensus* function, *--min-depth 1 --het-fract 1 --call-fract 0.5*). Only consensus sequences longer than 200 bp were retained for downstream analysis. Cases of multiple assemblies within a given orthogroup were treated as potential paralogs and subsequently inspected to remove spurious assemblies. First, identical assemblies (full length or partial) were removed using SeqKit v.2.7.0 (Shen *et al*., 2016) and CD-HIT v.4.8.1 (Fu *et al*., 2012). If multiple assemblies remained for a given orthogroup, these were aligned together with the reference sequences used for their assembly using MAFFT. A phylogenetic tree was then estimated using IQ-TREE v.2.1.3 (Minh *et al*., 2020; substitution model HKY) and rooted on the longest branch. Only assemblies that formed a monophyletic group with their corresponding references were validated as paralogs and retained for downstream analyses. In all other cases, only the longest assembly was retained. Steps that involved tree manipulation were implemented using Newick Utilities v1.6 (Junier & Zdobnov, 2010).

The performance of the custom assembly strategy was evaluated by reconstructing the Angiosperms353 sequences of two species from the reference dataset for which high-quality genomes are available (*Brachypodium distachyon* and *Oryza sativa*). For this, shotgun read datasets for these species were downloaded from the NCBI SRA database (accessions SRR891794 and SRR24031307) and subsampled to create four sets with varying sequencing depths (1, 5, 10 and 20×). Sequences were then assembled using our pipeline and compared to the sequences extracted from the reference genomes to assess the effect of sequencing depth on sequence completeness and identity, and on the recall of paralogs (Figs S2, S3).

### Extracting Angiosperms353 homologs from transcriptomes

To identify Angiosperms353 homologs in the transcriptome accessions, we performed a BLASTn search with the orthogroup reference dataset as query (e-value ≤ 10^-3^), and retained only hits with alignments covering > 50% of the query length and nucleotide identity > 70% for phylogenetic analysis. For the orthogroups corresponding to Angiosperms353 markers that were duplicated before the BOP–PACMAD split (see above), a BLASTn search was conducted and filtered as above, except that the query included the reference sequences of the two paralogous orthogroups. The putative homologous hits were then sorted into their corresponding orthogroups by aligning each hit with the query sequences using MAFFT, and estimating a tree using IQ-TREE. The hit was then assigned to one of the orthogroups based on the clade in which it was nested in the tree.

### Nuclear tree inference

We used the assembled sequences, including the validated paralogs, for inferring a species tree using a coalescent-based approach that accounts for paralogy, which has been shown to improve species tree estimation and vastly increase the data available for analysis (Smith & Hahn, 2021; Yan *et al*., 2021; Smith *et al*., 2022). Gene alignments were generated in a two-step approach. First, the reference sequences were aligned using MAFFT (*--maxiterate*=100) to generate a backbone alignment per gene. Then, gene assemblies of shotgun, target capture and transcriptome accessions were aligned one by one using the options *--addfragments* and *--keeplength* to improve the quality of the alignment of partially assembled sequences. Alignments were trimmed using trimAl v.1.4 (Capella-Gutiérrez *et al*., 2009) to remove columns with 90% or more missing data (*-gt* 0.1), and individual sequences shorter than 200 bp were removed from the trimmed alignments. To reduce uncertainty in tree estimation due to insufficient data, we discarded gene alignments with a total length of less than 500 bp after trimming. Finally, to further reduce the impact of missing data, only accessions with at least 50% of the total gene set were kept for analysis. The resulting dataset consisted of 1,153 accessions and 331 gene alignments. Gene trees were then inferred using RAxML v.8.2.12 (Stamatakis, 2014) with 100 rapid bootstrap pseudoreplicates. To strike a balance between computation time and modelling rate heterogeneity adequately, we used a GTR substitution model with a CAT rate heterogeneity approximation (25 rate categories; Stamatakis, 2006) across each alignment. Abnormally long branches detected using TreeShrink v.1.3.9 (Mai and Mirarab 2018; option *-q* 0.1) were then removed from the alignments, and the phylogenetic analysis was repeated. Finally, a multigene coalescent species tree was inferred using the resulting 331 gene trees with ASTRAL-Pro3 v1.17.3.5 (Zhang *et al*., 2020).

We evaluated tree stability across two additional data filtering strategies. In the first, the effect of missing data was assessed by increasing the alignment trimming threshold and removing columns with more than 50% missing data (all 1,153 samples retained). In the second filtered set, the same filtering strategy of the main dataset was used, but to be sure that our novel assembly methods were not biasing the results, we tested the impact of omitting the shotgun sequences altogether (841 tips retained; i.e. only accessions from target capture, transcriptome and complete genome sources). We compared support and conflict in the multigene coalescent tree across the three filtered sets using as metrics the local posterior support for the first and the second most preferred quartet configuration, respectively. We counted the number of matching branches (based on tip sets) of the additionally filtered sets compared to the main tree and summarised support and conflict at these branches.

### Gene tree–species tree reconciliation

We investigated the evidence for reticulations, whether from hybridisation, introgression, lateral transfers or incomplete lineage sorting. We used gene tree–species tree reconciliation under the maximum likelihood implementation of a duplication–transfer–loss model (UndatedDTL) in GeneRax v.2.0.1 (Morel *et al*., 2020). Running the analysis for the whole dataset was not feasible, so we performed a tribe-level reconciliation, where the species tree was collapsed to tribes, with gene trees matched to these tribes. We ran additional reconciliation analyses for three clades of economic importance and with well documented reticulation histories: subfamily Bambusoideae (bamboos), tribe Andropogoneae (maize, sorghum and relatives), and Triticeae (wheat and relatives).

From the gene trees, GeneRax infers, in addition to duplications and losses, gene transfers between two branches of the species tree. We summarised these transfers on the species tree using custom R scripts (see data repository, https://doi.org/10.5281/zenodo.10996136). Because an apparent transfer may also be an artefact of a poorly supported gene tree, we considered that a transfer between two branches had to be supported by at least five gene trees to indicate likely reticulation. Note that in the case of the tribe-level tree, this number of transfers combines gene tree tips from all species within a tribe. Transfers to/from the root were excluded (as they might involve any branch outside the ingroup that was not sampled). We highlighted the most frequent transfers as those with the top 10% quantile counts per species tree. We also evaluated, for each reticulate connection, if transfer counts were skewed in one direction by highlighting those with >50% proportional difference between counts in either direction.

### Plastome sequence assembly and tree inference

To compare the nuclear topology with the plastome topology, we inferred a 910-tip tree using the sequences of 70 coding plastome regions and the *trnL–trnF* intergenic region. We retrieved the 520 assembled plastome sequences that were already publicly available, representing in most cases shotgun accessions in the nuclear analysis, and sequences from the same species if the same accession was not available (see metadata table in data repository). New plastome CDS were assembled from shotgun and Angiosperms353 Illumina data using getOrganelle v1.7.5 (Jin *et al*., 2020) with default kmer settings for SPAdes (21,45,65,85,105) and 15 maximum extension rounds. We used a well-annotated plastome sequence (*Digitaria exilis*, INSDC accession KJ513091.1) as seed for assembly. The target sequences were then recovered from the full or partial assemblies via BLAST, with the *D. exilis* sequences as queries. Assemblies per sample were selected to cover at least 25% of the reference length for at least five genes or intergenic regions. Sequences were aligned per gene using MAFFT, gappy alignment columns were trimmed using the automated algorithm of trimAl v1.4.15 and all gene alignments concatenated using AMAS (Borowiec, 2016). After this step, accessions with 95% or more missing sites were removed, leaving the final 910 accessions. A maximum likelihood tree was then inferred using RAxML v.8.2.12 with a GTR-CAT model and 100 rapid bootstrap pseudoreplicates.

To measure to what degree nuclear relationships were supported by the plastome analysis, we mapped the transfer bootstrap expectation (TBE) metric (Lemoine *et al*., 2018) from 100 plastome bootstrap trees on the nuclear tree, after reducing both sets of trees to 751 tips we could match by accession, or, if the same accession was not available, by species. We also tested if nuclear–plastome conflict tends to affect branches where there is also conflicting signal within the nuclear genome. For this, we correlated the plastome TBE values with the local posterior support for the second-most supported quartet configuration as a measure of conflicting signal between nuclear gene trees.

## Results

### Nuclear reference dataset and genomic data

We compiled a grass-specific reference dataset for the assembly of 356 nuclear genes, available in the data repository (file “target_Ang353_sequences_grasses.zip”, https://doi.org/10.5281/zenodo.10996136). These genes were then extracted from genome and transcriptome sequences, and assembled from target-capture and shotgun data.

The final dataset used for phylogenetic analysis consisted of 1,153 accessions and 331 genes. Taxon occupancy was above 70% in 95% of all genes, and the number of genes recovered per accession ranged from 166 to 331 (median = 308). Median gene recovery was highest in shotgun accessions (98%), followed by transcriptomes (93%), target capture (92%) and genomes (91%) (Fig. S4a). Among shotgun accessions, gene recovery was correlated with sequencing depth, although sequencing depth as low as 1× was in most cases sufficient to recover sequences (> 200 bp) for more than 90% of all genes (Fig. S4b). Nonetheless, as expected, mean sequence completeness was higher among genome and transcriptome accessions (median = 85% and 83%) than in shotgun and target capture accessions (median = 63 and 60%; Fig. S4c). We were able to recover at least 80% of the Angiosperm353 genes (with sequences on average 49% complete) for the 17 target capture samples that had been originally enriched for 177 different nuclear loci (Fisher et al. 2016). Increasing filtering stringency overall reduced missing data, at the expense of reducing the number of tips and/or alignment length (Table S2). For example, mean alignment completeness was increased from 73 to 79% by increasing alignment trimming stringency, while reducing mean alignment length from 1160 to 864. Likewise, by removing the 312 shotgun accessions, mean alignment completeness was only slightly increased to 76% (Table S2).

### Nuclear genome phylogeny

Our 1,153-tip species tree was strongly supported (Fig. 1; see also full plot of the tree broken down into subclades in Suppl. Fig. S5). A single quartet configuration was preferred for most internal branches, with 80% having a local posterior support of 0.8 or higher, including all divergences at tribe level or above except Arundinelleae (0.7). For 65 internal branches (6%), all below tribe level, there was significant conflicting signal between gene trees, with two possible clade/quartet configurations each supported by one third or more of the gene trees (Fig. 1; hollow circles).

Conflict was distributed across the grass phylogeny and not concentrated in particular groups. Filtering the combined dataset more stringently had negligible effects on support or conflict (Fig. S6), both overall (Fig. S6a-b) and in direct comparison of matching branches (Fig. S6c-d), but the exclusion of assemblies from shotgun data led to slightly higher average bootstrap support in gene trees (Fig. S6e).

**Figure 1:**
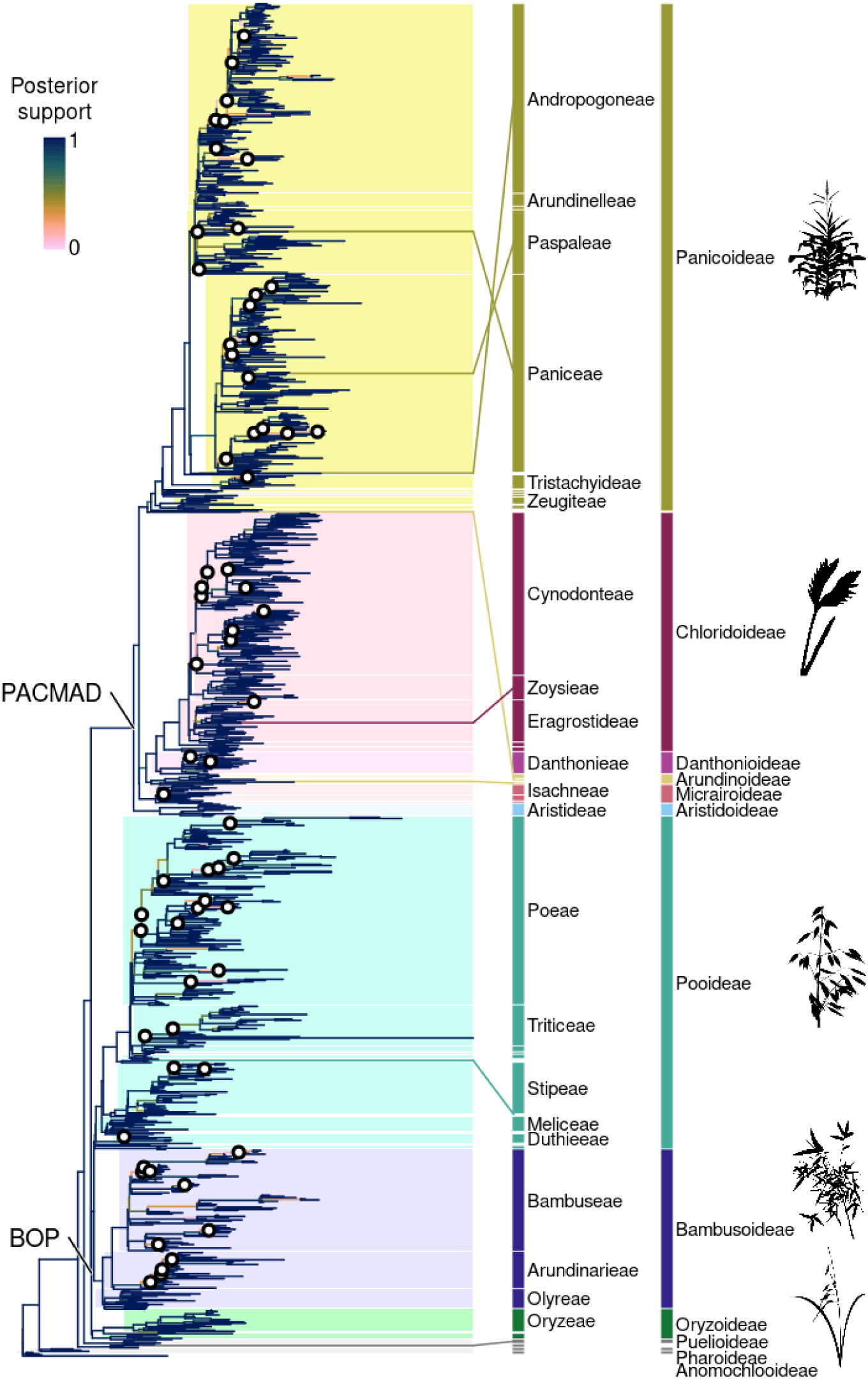
Phylogeny of 1,153 Poaceae accessions inferred from 331 nuclear genes, including paralogs, using a multi-species coalescent approach. Branch colours reflect local posterior support for the quartet configuration displayed. Hollow circles indicate supported conflict among nuclear gene trees at 48 internal branches, where two alternative quartet configurations each have >1/3 local posterior support. Subfamilies and larger tribes (abbreviated) are labelled according to the most recent Poaceae classification (Soreng et al., 2022). The coloured lines link taxonomic outliers at tribe to subfamily level to their nominal taxa. Silhouettes show representatives for large subfamilies (from top): Maize or corn, Zea mays (Panicoideae); Dactyloctenium radulans (Chloridoideae); oat, Avena sativa (Pooideae); Bambusa textilis (Bambudoideae); rice, Oryza sativa (Oryzoideae). See Fig. S5 for a detailed version of the tree.

We compared our tree to the most recent Poaceae classification (Soreng *et al*., 2022). The 1,153 accessions correspond to 1,133 accepted species, covering all but two (Anomochloeae and Streptogyneae) of the accepted tribes and 621 (79%) of the 791 genera. Twenty-one genera were sequenced for the first time: *Asthenochloa* Buse, *Bhidea* Stapf ex Bor, × *Cynochloris* Clifford & Everist, *Dilophotriche* (C.E.Hubb.) Jacq.-Fél., *Fimbribambusa* Widjaja, *Ekmanochloa* Hitchc., *Kaokochloa* De Winter, *Hydrothauma* C.E.Hubb, *Mniochloa* Chase, *Parabambusa* Widjaja, *Pinga* Widjaja, *Pogonachne* Bor, *Pommereulla* L.f., *Ratzeburgia* Kunth, *Ruhooglandia* S.Dransf. & K.M.Wong, *Spathia* Ewart, *Suddia* Renvoize, *Taeniorhachis* Cope, *Thedachloa* S.W.L.Jacobs, *Thyridachne* C.E.Hubb. and *Trilobachne* Schenck ex Henrard. All subfamilies were supported as monophyletic, except for the early-diverging Puelioideae which is paraphyletic, with its two genera *Guaduella* and *Puelia* forming separate lineages, as also noted by Huang *et al*. (2022). We found six further taxonomic discrepancies at tribe or subfamily level where tip positions did not match the taxonomy (Fig. 1, Table 1), and further cases of non-monophyly at the subtribe level (see detailed tree in Fig. S5).

Two accessions in surprising, isolated positions within Panicoideae (*Styppeiochloa hitchcockii* and *Ratzeburgia pulcherrima*, Table 1) passed all quality filtering steps. Manual investigation suggested unstable positions in gene trees, but no clear indication of a laboratory mix-up or contamination. Because no prior DNA data are available for these species, we retained them in the analysis but emphasise the need for further validation with independent samples.

**Table 1.** Taxonomic discrepancies in the nuclear tree at subfamily to tribe level. Taxa listed here will need follow-up studies to validate their placement. An asterisk (*) denotes genera whose type species was sampled.

| Genus/species | Nominal taxon | Nuclear tree position | Plastome tree position |
| --- | --- | --- | --- |
| *Amphipogon strictus* R.Br. | Arundinoideae: Arundineae | sister to Crinipedeae + Molinieae | in Arundinoideae: Arundineae |
| *Baptorachis foliacea* (Clayton) Clayton* | Paspaleae | Paniceae: Anthephorinae | Paniceae: Anthephorinae |
| *Chaetium festucoides* Nees* | Panicoideae: Paniceae | in Paspalinae, sister to *Streptostachys* | n/a |
| *Guaduella* Franch. | Puelioideae | sister to (*Puelia* + BOP + PACMAD) | sister to *Puelia* |
| *Neomolinia* Honda & Sakisaka* | Pooideae: Diarrheneae | sister to Brachypodieae + Triticodae + Poodae | n/a (sister to *Diarrhena* in Gallaher *et al.*, 2022) |
| *Ratzeburgia pulcherrima* Kunth* | Panicoideae: Andropogoneae: Ratzeburginae | sister to Paniceae | n/a |
| *Sporobolus subtilis* Kunth. | Chloridoideae: Zoysieae | in Eragrostideae, in *Eragrostis* | in Eragrostideae, in *Eragrostis* |
| *Styppeiochloa hitchcockii* (A.Camus) Cope | Arundinoideae: Crinipedeae | sister to Panicoideae | in Arundinoideae: Crinipedeae |

### Gene tree–species tree reconciliation

Reconciliation of gene trees with the species tree under a duplication–transfer–loss model suggests frequent reticulations in the grass family (Fig. 2, see also detailed plots in Fig. S6). The tribe-level reconciliation for the whole tree suggests reticulation early in the history of the grasses, involving the branch leading to the large crown group, the BOP–PACMAD clade (Fig. 2a). At this level of analysis, the most frequent reticulations primarily occurred in one direction (see arrows in Fig. 2a). Within Bambusoideae, the inferred transfers for both woody bamboo tribes, Arundinarieae and Bambuseae, reflect the allopolyploid origins of their subgenomes (Triplett *et al*., 2014; Guo *et al*., 2019; Chalopin *et al*., 2021; Ma *et al*., 2024). Note that in this tribe-level analysis, the number of transfers combine gene trees from all species within a tribe, i.e. high numbers could be driven either by a few genes or a few (or a single) species. We interpret transfers inferred from ancestors to descendants as transfers to a lineage that is now extinct (or not sampled in our tree) but descended from the same common ancestor.

Reconciliations at the species level (Fig. 2b–d) also support frequent reticulation. In the Andropogoneae and the Bambusoideae, the most frequent reticulations are not between deeper branches but within particular clades, such as within the Andropogoninae, the temperate woody bamboos (Arundinarieae), and, within the paleotropical woody bamboos (Bambuseae), the Malagasy Hickeliinae bamboos and the *Bambusa–Dendrocalamus–Gigantochloa* complex. In Triticeae, reticulation is frequent across the tribe. The assembled genome of the known allohexaploid *Thinopyrum intermedium* accounts for a large proportion of the highly supported transfers in Triticeae (species in bold in Fig. 2d). The origin of *Pascopyrum smithii* from past hybridisation between *Elymus* and *Leymus* (Dewey, 1975), and the origins of bread wheat, *Triticum aestivum*, from *Aegilops* ancestors are also evident.

**Figure 2:**
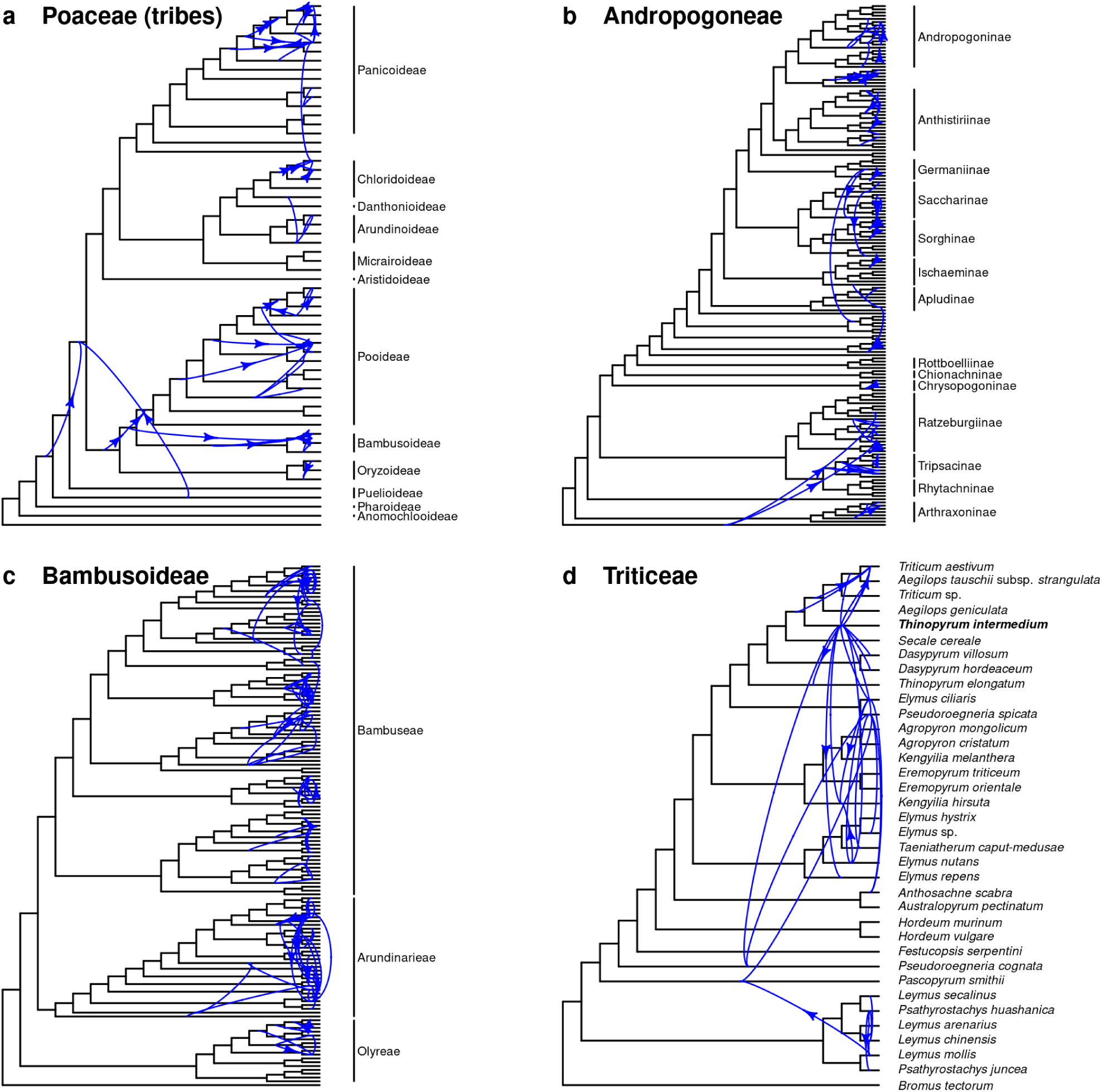
Nuclear gene reticulation in the grass family. For selected subgroups, the 331 gene trees were reconciled with the species tree under a duplication–transfer–loss model. Blue curves represent transfer events between two branches inferred at least five times (for different genes or within a gene family). Only the most frequent reticulations (top 10% quantile counts) are shown. Arrows indicate where transfers are highly skewed in one direction (>50% of proportional difference). Branch lengths are not proportional to time, and transfer lines start at the midpoint of a branch but the actual timing was not inferred. (a) Whole grass family, where tips were relabelled with tribes. Note that here, numbers of transfers combine gene trees from all species within a tribe. (b) Maize tribe, Andropogoneae. (c) Bamboos, Bambusoideae. (c) Wheat tribe, Triticeae. See also detailed plots in Fig. S7.

### Nuclear–plastome tree comparison

Nuclear–plastome conflict is uncommon across the grass phylogeny. We inferred a plastome tree for 910 accessions, representing 893 species, 508 genera and all tribes except Ampelodesmeae and Steyermarkochloeae (Fig. 3; see also detailed plot broken down into subclades in Fig. S5). Of these, 751 species, 53 tribes and 478 genera were also present in the nuclear tree and their relationships are compared between both trees (Fig. 3). Most branches above tribe level in the nuclear tree were also highly supported by plastome data. Only 54 branches showed strong signals of conflict, i.e. they were highly supported in the nuclear tree (local posterior probability > 0.8) but had poor support (TBE < 0.3) in the plastome tree, all of them at shallow levels (hollow circles in Fig. 3).

High support for a second local quartet configuration in the nuclear tree was negatively correlated with plastome TBE support (t = -5.25, p < 0.001; Pearson’s correlation test, two-sided), i.e. branches with higher intra-nuclear conflict also tend to show higher conflict with the plastome tree (Fig. 3, scatter plot inset).

When directly comparing the positions of clades at subfamily to tribe level, differences are evident in some cases. The Puelioideae genera *Guaduella* and *Puelia* are sister taxa in the plastome tree (Puelioideae) but not in the nuclear tree (paraphyletic Puelioideae). Arundinoideae and Micrairoideae are sisters in the plastome tree but paraphyletic in the nuclear tree. A striking difference was found in the position of *Styppeiochloa hitchcockii*, placed in Arundinoideae in classifications and in the plastome tree, but as sister to Panicoideae in the nuclear tree, based on the same target capture sample. In Pooideae, tribe Diarrheneae, although monophyletic in plastome trees (Gallaher *et al*., 2022), is polyphyletic in the nuclear tree, as its two genera *Diarrhena* and *Neomolinia* align in different clades. Triticeae plastomes appear to be paraphyletic with regard to Bromeae, as described previously (Bernhardt *et al*., 2017), while Triticeae is monophyletic in the nuclear tree. Finally, the nuclear tree grouped the two woody bamboo tribes, Arundinarieae and Bambuseae, which have distinct allopolyploid origins (Triplett *et al*., 2014; Guo *et al*., 2019; Chalopin *et al*., 2021; Ma *et al*., 2024), while in the plastid tree they are paraphyletic with regard to the herbaceous bamboos, Olyreae (Sungkaew *et al*., 2008). Below tribe level (see detailed tree in Fig. S5), the nuclear tree confirms previous studies in finding the C_4_-photosynthetic subtribe Anthephorinae (Paniceae) sister to the C_4_ MCP clade of Melinidinae, Cenchrinae and Panicinae (Washburn *et al*., 2015, 2017; Huang *et al*., 2022), while the chloroplast lineage of Anthephorinae is sister to the rest of Paniceae as in previous studies (GPWG II, 2012; Washburn *et al*., 2017; Saarela *et al*., 2018; Gallaher *et al*., 2022). Further differences in the branching order of subtribes are found in the tribes Arundinarieae (temperate woody bamboos), Bambuseae (tropical woody bamboos), Paspaleae, and Poeae.

**Figure 3:**
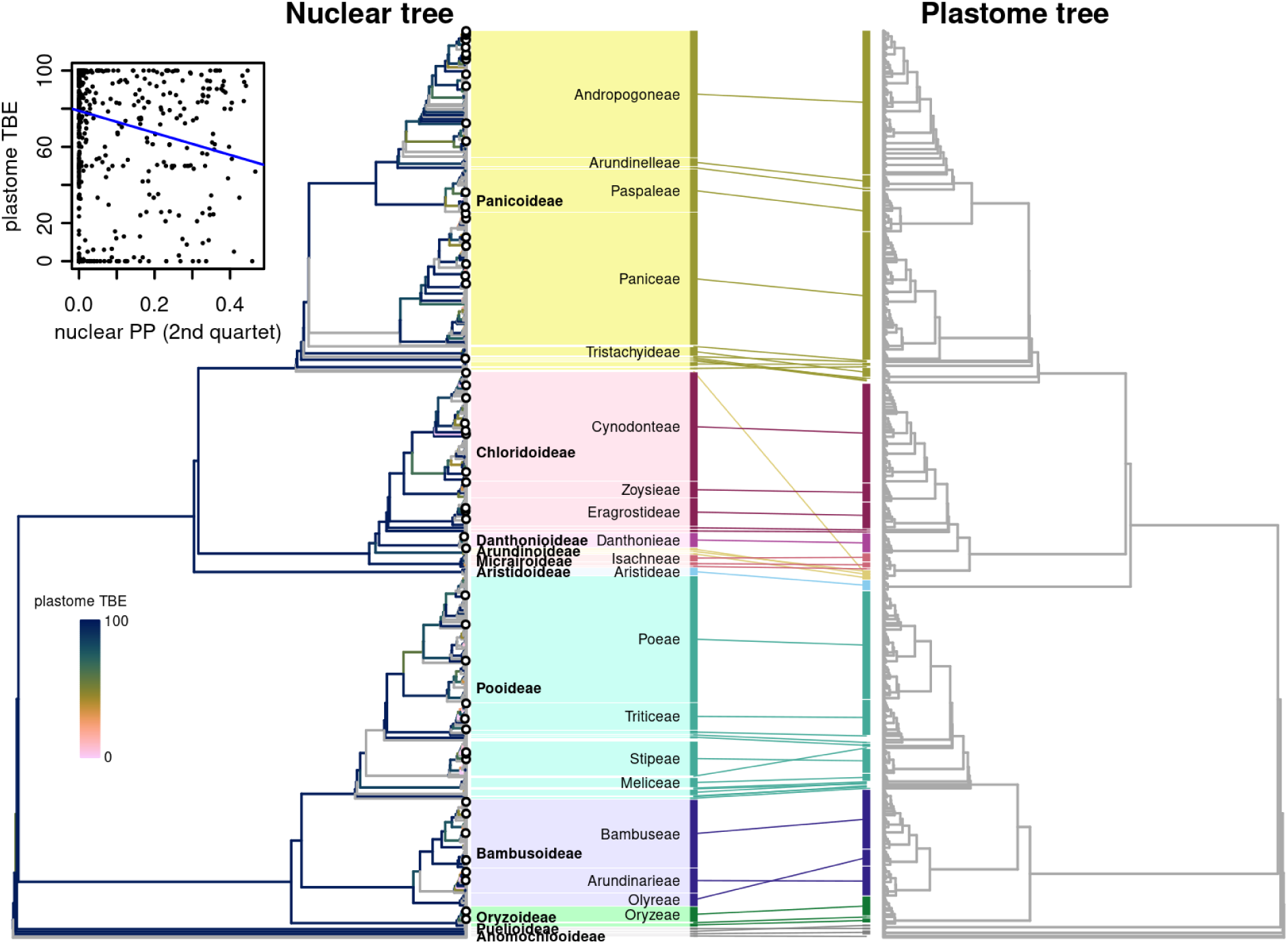
Comparison of nuclear and plastome topologies for the Poaceae. The 1,153-tip nuclear tree is shown on the left, the 910-tip plastome tree on the right. Plastome support (transfer bootstrap expectation, TBE) was summarised for branches present in both trees (814 shared species). Grey branches in the nuclear tree had no equivalent for comparison in the plastome tree. Hollow circles indicate strong signals of conflict, i.e. high support in the nuclear tree (local posterior probability > 0.8) but poor support (TBE < 0.3) in the plastome tree. Tribes are matched between the two in both trees, and larger tribes are labelled for orientation. The inset plots plastome support against a measure of conflict between nuclear gene trees (local posterior support for the second-most supported quartet per branch), which are negatively correlated. The blue line is a simple linear trend line.

## Discussion

### Nuclear phylogenomic data support relationships of current subfamilies and tribes

We show that the nuclear genome topology of the grass family overall supports existing subfamilies and tribes as monophyletic (Kellogg, 2015; Soreng *et al*., 2022). The subfamily-to tribe-level classification of the grasses has proven remarkably stable over previous community-wide phylogenetic efforts (GPWG, 2001; GPWG II, 2012). Our nuclear phylogenomic analysis further substantiates this framework, building on previous work to provide the largest nuclear phylogenomic sampling to date, with 79% of grass genera and all but two small tribes. Such sampling has previously been a considerable challenge in such a species-rich family. This phylogeny will help clarify generic limits and guide the search for useful genes and traits in wild relatives of cereal, forage, biofuel and turf crops.

Some taxonomic realignments will be necessary, despite the overall consistency with previous work. In addition, the placement of a few taxa will need to be validated by additional sequences or samples (Table 1). These may represent cases of biological interest (e.g., true reticulations) but our current data cannot entirely rule out possible technical artefacts. More taxonomic mismatches will require attention at the subtribe level. Using the sequence data of Huang *et al*. (2022) we were able to reproduce their results suggesting paraphyly of subfamily Puelioideae (*Guaduella* and *Puelia*) in the nuclear tree, although this result was not tested with independent plant samples. If the cyto-nuclear conflict continues to be supported, the well-supported monophyly of the group in the plastid phylogeny would suggest an introgression event in the early history of the grasses. Future morphological studies are needed to determine whether there are characters that support either monophyly or paraphyly of Puelioideae, one of the least well known of the grass subfamilies. More generally, as nuclear genome-scale data continue to accumulate, the grass taxonomic community will have to decide whether the nuclear genome, ultimately underlying most phenotypic characters, should dictate taxonomy in case of conflicting signals. In the bamboos, the nuclear topology better reflected morphological differences than the plastome phylogeny in previous work (Wang *et al*., 2017). In the Paniceae, the position of Anthephorinae sister to Melinidinae–Cenchrinae–Panicinae is in line with a common origin of C_4_ photosynthesis in the combined clade (Washburn *et al*., 2015), with two separate plastome sources. We refer taxonomic and nomenclatural changes to further studies by specialists of the relevant grass subgroups.

Nuclear–plastome discordance is rare between higher taxonomic levels in the grasses. In the large PACMAD clade, we confirm previous nuclear (Bianconi *et al*., 2020; Huang *et al*., 2022) and plastome studies (GPWG II, 2012) in finding Aristidoideae sister to the other five subfamilies, whereas more recent plastome studies (Saarela *et al*., 2018; Duvall *et al*., 2020; Gallaher *et al*., 2022) suggested this position might be artifactual and favoured a ‘panicoid sister’ hypothesis.

Arundinoideae and Micrairoideae subfamilies do not form a clade as in the plastome tree, a result also found in the nuclear analysis of Huang *et al*. (2022), but we note that there appears to be some gene tree conflict around the divergence of these subfamilies. There appears to be no concentration of gene tree conflict at the base of PACMAD, a period of rapid grass diversification (Christin *et al*., 2014). Across angiosperms, episodes of rapid diversification are correlated with higher conflict among gene trees (Guo *et al*., 2023; Zuntini *et al*., 2024). Further investigation of this relationship for the grasses could build on the dataset we compiled here but will require tackling the complex issue of time calibration in grasses (see below).

### Reticulation has been frequent in the grass family

Reticulation within the nuclear genome occurred repeatedly across the grass phylogeny, although this is not sufficient to mask the underlying species tree structure. Approximately 45-80% of grasses are polyploid (Stebbins, 1985; DeWet, 1986), with an appreciable proportion of those being allopolyploid, i.e., hybrids. The methods used here are not designed to detect allopolyploidy but are still able to identify frequent reticulation events. The actual modes of reticulation in grasses certainly need more study, as we cannot distinguish here between introgression and hybridisation. Recent work also demonstrated the frequency of lateral gene transfers of large blocks in the genomes of *Alloteropsis semialata* (Dunning *et al*., 2019; Raimondeau *et al*., 2023) and other grass species (Hibdige *et al*., 2021). Contamination and gene-tree errors can potentially obscure patterns in short-read data as we included in our analysis, but encouragingly, we retrieved known patterns such as the mosaic origins of the *Thinopyrum intermedium* genome (Mahelka *et al*., 2011). This suggests that reduced-representation nuclear datasets do retain signals of reticulation.

Our analysis offers a glimpse of how the accumulation of assembled genome data for grasses beyond model and crop species could foster research on reticulation, particularly in three areas. First, clarifying where apparent reticulate relationships may actually stem from differential retention of homologs after whole-genome duplication. For example, the reticulations we inferred at the base of the BOP-PACMAD clade, the large crown radiation of grasses, could potentially be remnants from the *rho* whole-genome duplication event at the stem of Poaceae (McKain *et al*., 2016; Zhang *et al*., 2024). Second, correlating reticulation frequency with ecological and morphological predictors to identify the physical mechanisms of lateral transfers, which remain speculative (Pereira *et al*., 2023). Third, using synteny information to identify the precise locations and origins of functional variation, potentially using new deep learning approaches for identifying introgression (Zhang *et al*., 2023). New crops, such as *Thinopyrum intermedium* (intermediate wheatgrass or kernza) with its mosaic genome and its potential as perennial cereal or genetic resource (Baker *et al*., 2020), are certainly prime candidates for such research. However, given the frequency of allopolyploidisation, and if lateral transfers are as frequent as recent work suggests (Hibdige *et al*., 2021), the grass family as a whole may well constitute a ‘single genetic system’ (Freeling, 2001; Mascher *et al*., 2024) or higher-level ‘pangenome’ (Dunning *et al*., 2019), so species-level sampling including the more distant relatives of crops is needed to access the entire genetic diversity potentially available for future sustainable agriculture.

### Towards a complete grass tree of life

Our community effort resulted in the most comprehensive nuclear phylogenomic tree for the grass family to date, including 1,133 species, with five genera sequenced for the first time. This tree, and the dataset associated with it, paves the way towards placing all approximately 11,800 species in the grass tree of life. Already, the International Nucleotide Sequence Database Collaboration hosts sequence data for more than 6,200 grass species, as of April 2024. The comprehensive phylogenomic backbone we provide here could provide a basis for assembling these shorter sequences into a grass supertree for analyses of trait evolution and biogeography, as attempted previously with smaller Poaceae backbones (Spriggs *et al*., 2014; Elliott *et al*., 2024).

We show that the Angiosperms353 gene set can be successfully used to anchor different types of genomic datasets, including unenriched Illumina sequence data. Sequencing depth and paralog recovery obviously vary across such different datasets, which needs to be taken into account, e.g. in future studies of whole-genome duplications and events of auto- and allopolyploidy (Thomas *et al*., 2017; Morales-Briones *et al*., 2021; Rothfels, 2021) or large-scale gene duplications that preceded major innovations like cold tolerance (Schubert *et al*., 2020; Zhang *et al*., 2022). The improved Angiosperms353 reference set constructed here for the grasses will facilitate the inclusion of previously unsequenced grass species. This target capture approach allows in particular sequencing degraded DNA from herbarium specimens in a cost-efficient manner and thus filling the remaining gaps of the grass tree of life, even where there are logistical barriers to obtaining high molecular weight DNA for full genome sequencing.

The timeline of grass evolution continues to be a matter of debate, with recent studies suggesting a mid-Cretaceous origin for the grasses (Huang *et al*., 2022; Gallaher *et al*., 2022) and thus supporting earlier suggestions based on phytolith fossils (Prasad *et al*., 2005, 2011). However, such age estimates hinge on several factors (Christin *et al*., 2014), such as the placement of phytolith fossils, appropriate modelling of rate correlation, and the upper bound set by the age of flowering plants, which itself remains unclear and fraught with methodological challenges (Brown & Smith, 2018; Silvestro *et al*., 2021; Sauquet *et al*., 2022; Carruthers & Scotland, 2023). The nuclear dataset we provide here, along with recent advances in grass phytolith classification (Gallaher *et al*., 2020) as well as a better understanding of rate variation across branches (Carruthers *et al*., 2020; Carruthers & Scotland, 2020) and gene tree conflict (Carruthers *et al*., 2022) on divergence time estimation suggest a new comprehensive analysis of grass divergence times as a promising avenue forward.

## Acknowledgements

We thank the herbarium curators at BR, CAN, EA, FTOH, KIB, MEL, NSW, P, PE, SI, TAN and US for supporting this project and the following people for providing technical support or advice at various stages: Paul Bailey (RBG Kew), Kevin Lempoel (RBG Kew), Sophie Manzi (CNRS Toulouse) and Martin R. Smith (University of Durham). We thank M.C. Romay and E.S. Buckler for pre-publication access to sequences from the Panandropogoneae project.

We dedicate this paper to the memory of W. Derek Clayton who passed away in September 2023. His contribution to grass systematics can hardly be overestimated and laid the foundations for many of the advances in this paper.

## Data Accessibility

Data used and produced in this study, including metadata for all accessions, are available in an open Zenodo repository (https://doi.org/10.5281/zenodo.10996136). Accession numbers for new sequence data will be made available at publication.

## Author contributions

W.J.B., M.E.B., P.-A.C., L.T.D., J.H., E.A.K., R.J.S., M.S.V. conceptualised the study. M.D.B., R.L.B., G.B., M.E.B., P.-A.C, P.C., D.M.C., G.D., L.T.D., M.R.D., S.Z.F., S.F., J.H., T.R.H., W.H., R.W.J., E.A.K., J.M.K., X.L., O.M., T.G.M., M.M., D.J.M., J.R., L.S., R.J.S., M.S.V., M.W., C.A.W., M.D.X., L.Z., F.O.Z. curated the data. M.E.B., J.H. performed formal analysis. W.J.B., G.B., P.-A.C., J.T.C., D.M.C., F.F., E.A.K., L.Z., A.Z. acquired funding. M.E.B., J.T.C., J.H., I.L. performed investigation. W.J.B., M.E.B., P.-A.C., L.T.D., J.H., A.M.H., E.A.K., R.J.S., M.S.V., A.R.Z. developed methodology. W.J.B., M.E.B., P.-A.C., J.H., M.S.V. had administrative responsibility in the project. W.A., M.D.B., R.L.B., J.B., G.B., J.L.B., P.C., W.C., M.C., L.G.C., C.A.C., D.M.C., G.D., M.R.D., .D., A.E.F., S.F., F.F., L.J.G., T.H., T.R.H., C.H., R.W.J., E.A.K., C.J.K., J.M.K., I.L., R.L., D.L., J.L., X.L., Q.W.L., H.M., T.D.M., O.M., M.R.M., T.G.M., D.J.M., O.P.N., G.E.O., P.M.P., R.A.R., J.R., J.M.S., L.S., N.W.S., R.J.S., M.S., E.J.T., P.T., G.A.V., M.S.V., N.G.W., J.D.W., T.W., M.W., C.A.W., M.D.X., N.X., L.Z., F.O.Z. provided resources. M.E.B., J.H. developed software code. W.J.B., P.-A.C., E.A.K., R.J.S., M.S.V. supervised the project. G.B., M.E.B., L.T.D., J.H., E.A.K., J.M.K., D.L., J.L., H.M., R.J.S., M.S., M.S.V., A.R.Z. validated the results. R.L.B., M.E.B., J.H. visualised the data. M.E.B., J.H. wrote the original manuscript. W.A., W.J.B., M.D.B., R.L.B., J.B., G.B., M.E.B., P.-A.C., W.C., P.C., L.G.C., D.M.C., L.T.D., A.E.F., F.F., J.H., T.R.H., W.H., A.M.H., R.W.J., E.A.K., J.M.K., I.L., D.L., H.M., M.R.M., T.G.M., D.J.M., J.M.S., N.W.S., R.J.S., M.S., M.S.V., N.G.W., J.D.W., M.W., C.A.W., L.Z., A.Z. reviewed and edited the manuscript.

