## Supplementary Tables for "Nuclear phylogenomics of grasses (Poaceae) supports current classification and reveals repeated reticulation"

**Table S1.** Nuclear data statistics.

| | Main dataset | Stringent trimming | Main w/o shotgun data |
|---|---|---|---|
| Total accessions | 1153 | 1153 | 841 |
| Total genes | 331 | 315 | 331 |
| Alignment length (bp)<br>mean; median (range) | 1160; 1002 (501–4411) | 864; 1011 (502–3564) | 1160; 1002 (501–4411) |
| Taxon occupancy (%)<br>mean; median (range) | 89; 93 (15–99) | 90; 92 (14–99) | 89; 93 (14–99) |
| Alignment completeness (%)<br>mean; median (range) | 73; 74 (52–91) | 79; 79 (66–92) | 76; 77 (47–95) |
| Genes recovered per accession<br>mean; median (range) | | | |
| Overall | 296; 308 (166–331) | 282; 294 (156–315) | 294; 306 (166–328) |
| Reference genomes | 286; 300 (167–328) | 273; 285 (163–312) | 286; 300 (167–328) |
| Transcriptomes | 300; 308 (170–327) | 287; 294 (165–311) | 300; 308 (170–327) |
| Target capture | 290; 304 (166–327) | 278; 290 (161–312) | 290; 304 (166–327) |
| Shotgun | 301; 324 (168–331) | 285; 308 (156–315) | - |
| Mean sequence completeness per accession (%)<br>mean; median (range) | | | |
| Overall | 65; 67 (14–98) | 70; 75 (15–99) | 67; 68 (20–98) |
| Reference genomes | 82; 85 (43–98) | 83; 86 (45–99) | 82; 85 (43–98) |
| Transcriptomes | 80; 83 (32–95) | 83; 86 (34–96) | 80; 83 (32–95) |
| Target capture | 56; 60 (20–83) | 64; 69 (23–89) | 56; 60 (20–83) |
| Shotgun | 61; 63 (14–97) | 64; 66 (15–97) | - |
