## Supplementary Methods for "Nuclear phylogenomics of grasses (Poaceae) supports current classification and reveals repeated reticulation"

### DNA isolation, library preparation and sequencing

*Angiosperms353 accessions, Genomics of Australian Plants (GAP)*
All laboratory work was performed at the Australian Genome Research Facility, Melbourne, with the exception of some in-house DNA extractions performed using a Qiagen DNeasy Plant Mini Kit (Qiagen), preceded by a D-sorbitol pre-wash. Genomic DNA from herbarium specimens was extracted using the NucleoSpin Plant II Kit (Macherey-Nagel, Düren, Germany). When not already sheared, extracted DNA was fragmented enzymatically to ~350 bps. DNA libraries were prepared with NEBNextUltra II FS Library Prep Kit (New England Biolabs, Ipswich, MA, USA). Hybridisation capture was conducted on pools of 12–16 libraries, enriched using the Angiosperms353 v1 probe kit (Johnson *et al.*, 2019) with V5 chemistry produced by MYbaits kit (Daicel Arbor Biosciences, Ann Arbor, MI, USA;Cat. #308108.v5). Enriched libraries were sequenced on Illumina NovaSeq 6000 SP (Illumina Inc., San Diego, USA) with v1.5 chemistry and 150bp paired-end reads.

*Angiosperms353 accessions, Catalán (Loliinae)*
Loliinae leaf samples were analysed for capturing single-copy nuclear gene targets with the Angiosperm353 kit at Arbor Biosciences (Michigan, USA). Capture reactions were pooled in equimolar ratios and sequenced on a partial lane of the Illumina NovaSeq 6000 platform in paired-end (PE) mode (2 x 150 bp).

*Illumina shotgun accessions, Besnard/Christin/Watcharamongkol*
Low-coverage sequencing was performed using Illumina technology. Genomic DNA was isolated from ca. 5–10 mg of leaf material using the BioSprint 15 DNA Plant Kit (Qiagen). Between 100 and 500 ng of double-stranded DNA were used to construct sequencing libraries with the Illumina TruSeq Nano DNA LT Sample Prep kit (Illumina, San Diego, CA, USA), following the manufacturer's instructions (for more details, see Besnard *et al.*, 2018). Each sample was multiplexed with samples from the same or different projects and paired-end sequenced on 1/24th of an Illumina HiSeq3000 lane.

*Illumina shotgun accessions, Duvall*

Of the genome skims contributed to this project, 80 had been used in previously published papers of complete plastomes where detailed sequencing methods can be found (Jones *et al.*, 2014; Cotton *et al.*, 2015; Saarela *et al.*, 2015, 2018; Wysocki *et al.*, 2015; Burke *et al.*, 2016b,a; Duvall *et al.*, 2016, 2017; Attigala *et al.*, 2016; Orton *et al.*, 2017, 2021; Burke, 2018). In brief, total DNA was extracted from silica dried, or in some cases fresh leaf tissues, by manual homogenization of tissue in liquid nitrogen followed by use of the DNeasy Plant Mini Kit protocol (Qiagen, Valencia, CA, USA). DNA extracts were quantified with the Qubit assay (Invitrogen, ThermoFisher Scientific, Wilmington, DE, USA), and diluted to 2.5 ng/µl in 20 µl sterile water. DNA libraries were prepared from the purified extracts in the laboratory of M. R. Duvall (Northern Illinois University, DeKalb, IL, USA) using Illumina Prep Kits (Illumina Inc., San Diego, CA, USA). DNA libraries were purified with Clean and Concentrator kits (Zymo Research, Irvine, CA, USA). Standard manufacturer protocols for each respective sample preparation kit were used. Sequencing by synthesis was performed at the Core DNA Facility at Iowa State University (Ames, IA, USA) on Illumina HiSeq platforms.

*Illumina shotgun accessions, Fjellheim*

Leaves were flash frozen in liquid nitrogen before they were homogenised using a TissueLyser (QIAGEN®,Valencia, CA, USA) at 30 Hz for 2x1 minute. DNA was extracted using the DNeasy Qiagen kit (QIAGEN®) following the manufacturer's protocol. Strand specific libraries were prepared and paired-end sequenced using Illumina HighSeq 2000 or Illumina HighSeq 4000 and HighSeq X platforms at Novogene Co. Ltd (Cambridge, UK) or Norwegian Sequencing Center (Oslo, Norway), respectively.

*PANAND samples, Kellogg*

Leaves were lyophilised and ground for DNA extraction with a Qiagen Kit (Qiagen Inc., Germantown, MD). DNA concentration was estimated. Depending on the concentration, libraries were constructed using Illumina Tru-Seq or nano Tru-Seq. 24 samples were pooled and sequenced in 1 lane of an S4 flowcell on an Illumina Novaseq 6000 System. Reads were 150 bp, paired-end.

**Curation of the grass-specific Angiosperms353 reference dataset**

The initial reference dataset was inspected to identify cases of inaccurate split of orthogroups, and to remove non-homologous sequences and pseudogenes.

First, orthogroups containing homologs of more than one Angiosperms353 gene were identified and subsequently aligned using MAFFT v.7.481 (Katoh & Standley, 2013); individual sequences were removed if belonging to an Angiosperms353 gene that was different from the majority, following confirmation by visual inspection of the alignment.

Second, Angiosperms353 genes that were split into two or more orthogroups were analysed to verify if the orthogroups a) indeed corresponded to paralogs at the level of the most recent common ancestor of the BOP-PACMAD clade (see Methods), b) were incorrectly split by Orthofinder and therefore should be merged into a single orthogroup, or c) included non-homologous sequences. For that, orthogroups from the same Angiosperms353 gene were grouped and aligned using MAFFT, and a phylogenetic tree was inferred using IQ-TREE v.1.6.12 (Nguyen *et al.*, 2015). We inspected the trees to decide whether a) to keep the orthogroups as separate Angiosperms353 paralogs (orthogroups forming a sister group in the tree), b) merge them into a single orthogroup (sequences from distinct orthogroups placed in their expected lineages within BOP-PACMAD), or c) remove non-homologous sequences (sequences placed outside BOP-PACMAD).

Finally, we inferred gene trees for all orthogroups to identify any remaining non-homologous sequences and pseudogenes. For this, sequences were aligned using MAFFT, and alignments were trimmed to remove columns with >25% missing data using trimAl v.1.4 (Capella-Gutiérrez et al., 2009), with sequences <200 bp being subsequently removed. Gene trees were then inferred using RAxML v.8.2.12 (Stamatakis, 2014) with the GTR-CAT model. Sequences leading to abnormally long branches were flagged using TreeShrink v.1.3.9 (Mai and Mirarab, 2018) and subsequently removed from the dataset. The final reference dataset was then built with the trimmed sequences extracted from the curated orthogroup alignments.
