## Supplementary Figures for "Nuclear phylogenomics of grasses (Poaceae) supports current classification and reveals repeated reticulation"

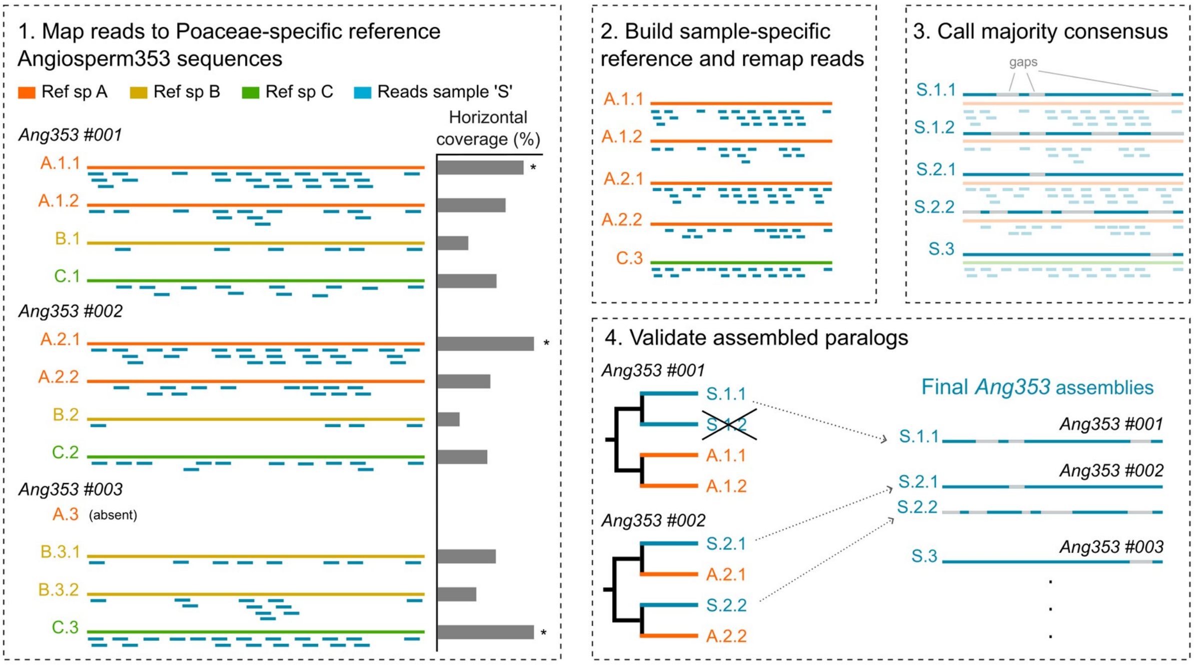


**Figure S1.** Schematic overview of custom workflow for sequence assembly from Illumina shotgun accessions. Long bars represent sequences from three different reference species (A, B and C), and short blue bars represent shotgun reads from a sample (S). (1) Reads were mapped to Angiosperm353 reference sequences, which included, for each orthogroup (e.g. #001, #002, #003, etc), all sequences belonging to each reference species, including paralogs (e.g. ‘A.1.1’ and ‘A.1.2’ are paralogs of gene ‘#001’ from reference species ‘A’). In this step, the sequence with the highest horizontal coverage (i.e. percentage of the reference sequence that is covered by mapped reads; represented by the bar plot on the right) was recorded for each orthogroup (indicated here with an asterisk). (2) A new, sample-specific orthogroup reference dataset was then built with the reference sequences recorded in the previous step, including their paralogs, if any (e.g. for gene #001, sequence A.1.1 had the highest coverage, therefore both sequences of reference species ‘A’ – A.1.1 and A.1.2 – were included in the new reference dataset). Reads were then mapped to this new reference dataset. (3) A majority consensus was called from the read alignments; bases with no coverage were called as gaps. (4) To distinguish potential paralogs from spurious assemblies in cases of multiple assemblies within a given orthogroup, sequences were inspected using a phylogenetic approach. For each orthogroup, a phylogenetic tree was inferred using the assemblies and their respective references. Here, an assembly was retained only if it formed a clade with the reference used for its assembly.

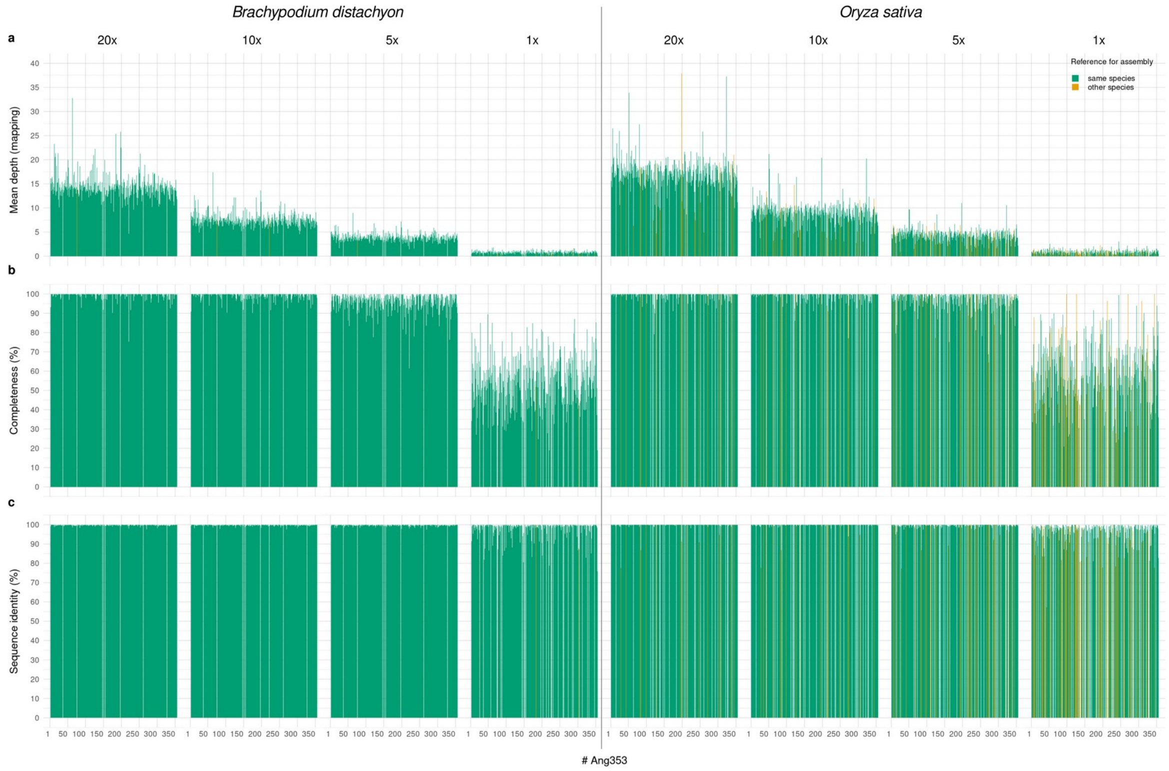


**Figure S2.** Test of custom assembly workflow on full genome sequences. For each of the two species tested (*Brachypodium distachyon*, left and *Oryza sativa*, right), four short read datasets with varying sequencing depths (20x, 10x, 5x and 1x) were generated by subsampling the original short read dataset downloaded from a public database. Sequences for each of the Angiosperm353 genes (*x* axis) were then assembled using the custom pipeline (see Methods and Fig. S1 for details). (a) Mean depth computed from read mapping for each gene. (b) Sequence completeness. (c) Sequence identity. Bars with different colours correspond to sequences that were assembled using as reference a sequence from the same species (green) or from a different one (orange).

a Overall copy number recall

b Multi-copy gene recall

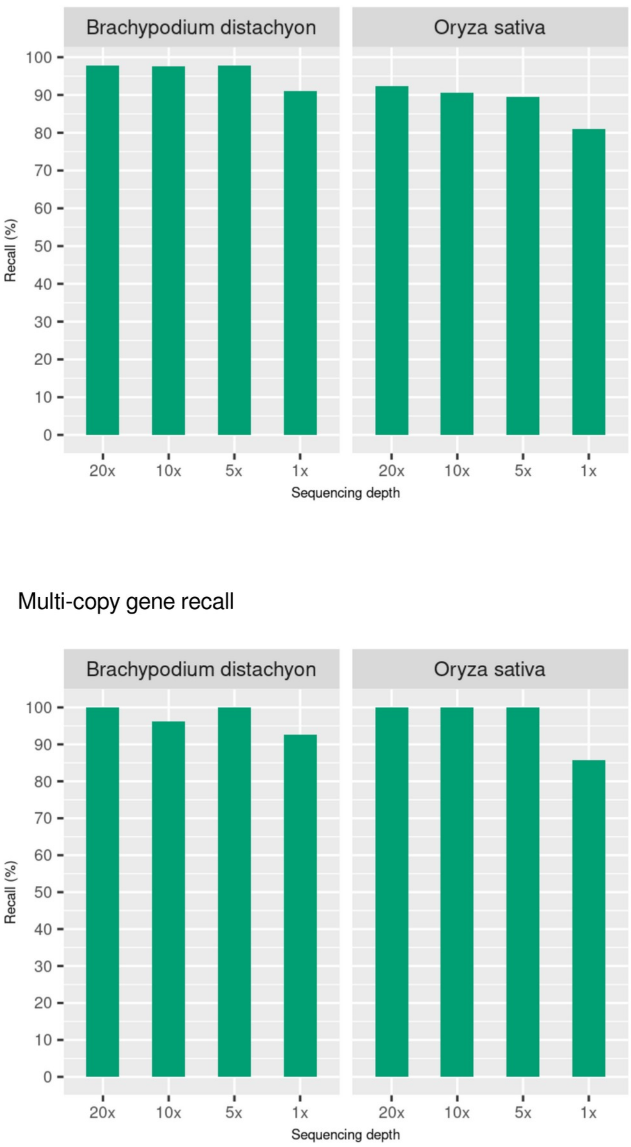


**Figure S3.** Test of custom assembly workflow on full genome sequences – copy number recall. (a) Overall copy number recall, i.e. of all Angiosperms353 genes, those for which the number of assembled sequences in the short read dataset is equal to the number of sequences in the reference genome. (b) Multi-copy gene recall, i.e. of all Angiosperms353 genes represented by multiple copies in the reference genome (n = 27 in *B. distachyon*, and n = 14 in *O. sativa*), those for which more than one sequence was assembled in the short read dataset (i.e. n > 1 in the short read dataset when n > 1 in the reference genome).

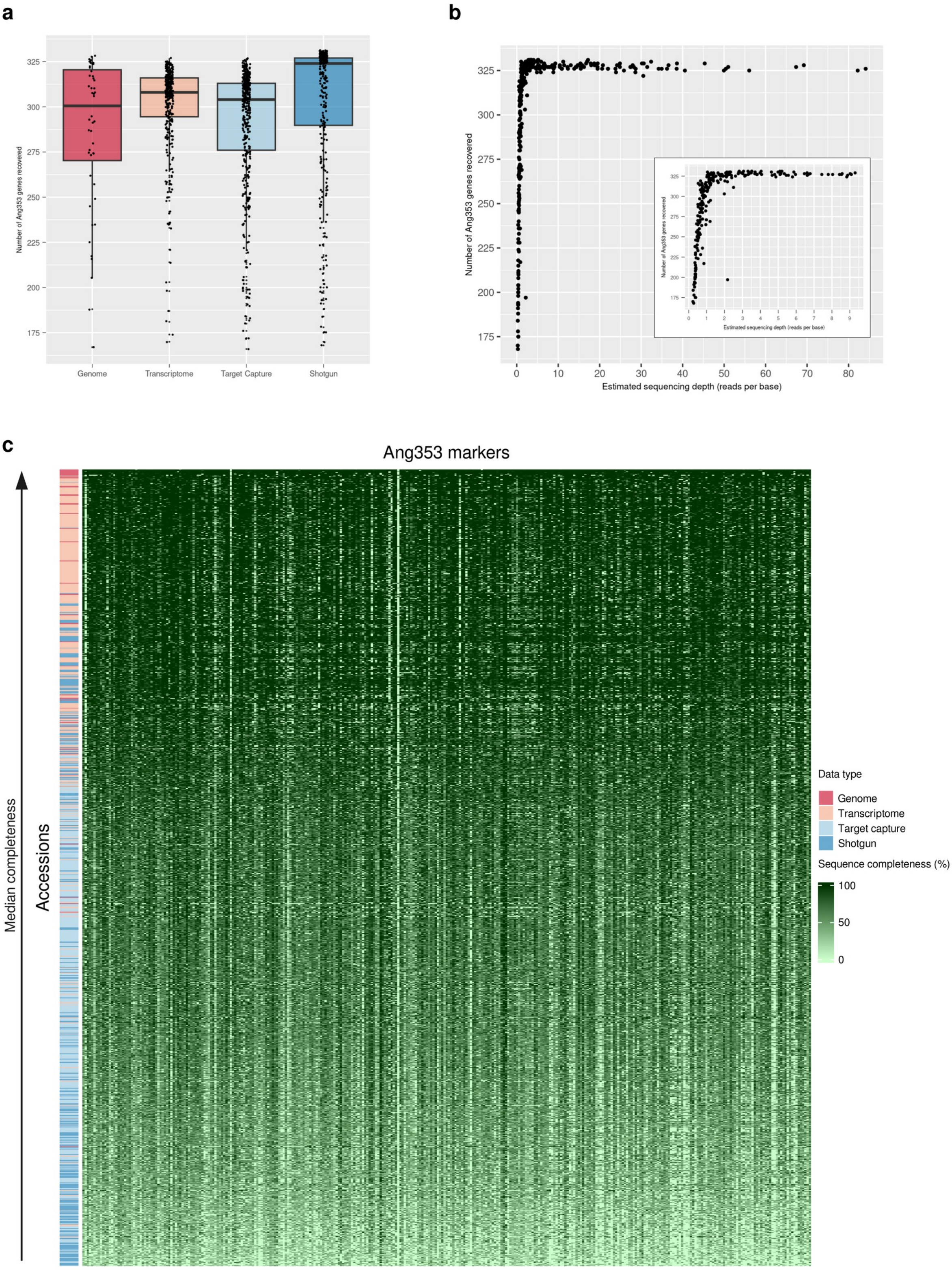
a
Number of Ang353 genes recovered
325
300
275
250
225
200
175
Genome
Transcriptome
Target Capture
Shotgun
b
Number of Ang353 genes recovered
Estimated sequencing depth (reads per base)
0
10
20
30
40
50
60
70
80
c
Ang353 markers
Median completeness
Accessions
Data type
Genome
Transcriptome
Target capture
Shotgun
Sequence completeness (%)
100
50
0

**Figure S4 (previous page).** Nuclear gene recovery and sequence completeness. (a) Boxplot of the number of Angiosperm353 genes recovered per accession, according to the type of data. (b) Effect of sequencing depth on the number of genes recovered per shotgun accession. The embedded plot is a zoomed view of accessions with estimated sequencing depth < 10x. (c) Heatmap of sequence completeness across all Angiosperm353 genes (*x* axis) and accessions (*y* axis). Accessions are sorted by median completeness (highest on top).

**Figure S5 (following pages).** Detailed version of the multispecies coalescent nuclear species tree (Fig. 1 in the main text) inferred for 311 genes and 1,153 accessions, broken down into subclades. Pies at nodes indicate the frequency of gene trees supporting each quartet and text at nodes gives the local posterior probability of the quartet shown. Tip labels show data type, species and voucher, isolate or germplasm information, where available, for each accession. Taxa from subtribe to subfamily level are labelled with coloured polygons. Taxonomic outliers falling outside the clade corresponding to their nominal taxon are labelled in brackets after the accession information.

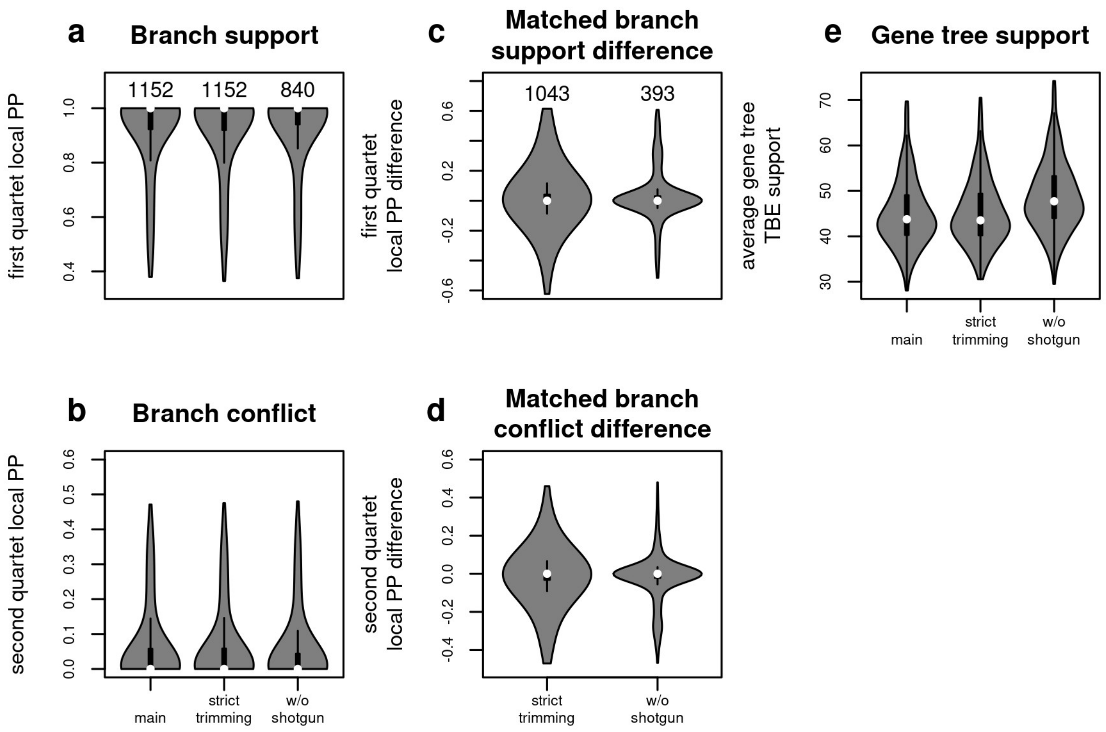


**Figure S6.** Nuclear species tree stability under different data filtering strategies. The main tree was inferred from gene alignments where columns with > 90% missing data were removed (331 genes > 500 bp retained), and included only accessions with at least 50% of gene recovery (i.e. > 166 genes). This tree was compared to trees obtained from an alignment set with more stringent trimming threshold (removal of columns > 50% missing data, 315 genes > 500 bp retained; ‘stringent trimming’), and a set that excluded shotgun accessions from the main tree dataset (‘w/o shotgun’). (a) Internal branch support (local posterior support for the preferred quartet configuration) across the three species trees. Numbers above violin plots give the number of internal branches per tree. (b) Internal branch conflict (local posterior support for the second-most supported quartet configuration) across the three species trees. (c) Difference in support for branches recovered with the two reduced alignment sets compared to the main tree. Numbers above violin plots give the number of matching branches. (d) Difference in conflict for branches recovered with the two reduced alignment sets compared to the main tree. (e) Mean gene tree support (transfer bootstrap expectation) across the three species trees.

**Figure S7 (following pages).** Detailed plots of the reticulations inferred with gene tree–species tree reconciliation (Fig. 2 in the main text). The black phylogeny represents the species tree. Blue lines correspond to inferred reticulate connections (transfers). Very frequent transfers (upper 10% quantile of the number of genes involved, see inset histogram) are coloured in darker blue and less frequent transfers in lighter blue. Arrowheads indicate where transfer counts are skewed by more than 50% in one direction. (a) Full Poaceae tree at tribe level, where accessions were mapped to their respective tribes. (b) Andropogoneae tribe (maize, sorghum, sugarcane and relatives). (c) Bambusoideae subfamily (bamboos). (d) Triticeae tribe (wheat, barley and relatives).

**Figure S8 (following pages).** Detailed version of the plastome tree inferred from a concatenated alignment of 70 coding regions and *trnL-trnF* (tree on the right in Fig. 3 in the main text) for 910 accessions, broken down into subclades. Text labels at nodes give the branch support as transfer bootstrap expectation. Tip labels show data type, species and voucher, isolate or germplasm information, where available, for each accession. Taxa from subtribe to subfamily level are labelled with coloured polygons. Taxonomic outliers falling outside the clade corresponding to their nominal taxon are labelled in brackets after the accession information.

### Figure S5 − Nuclear tree (Astral−PRO based on 331 gene trees)

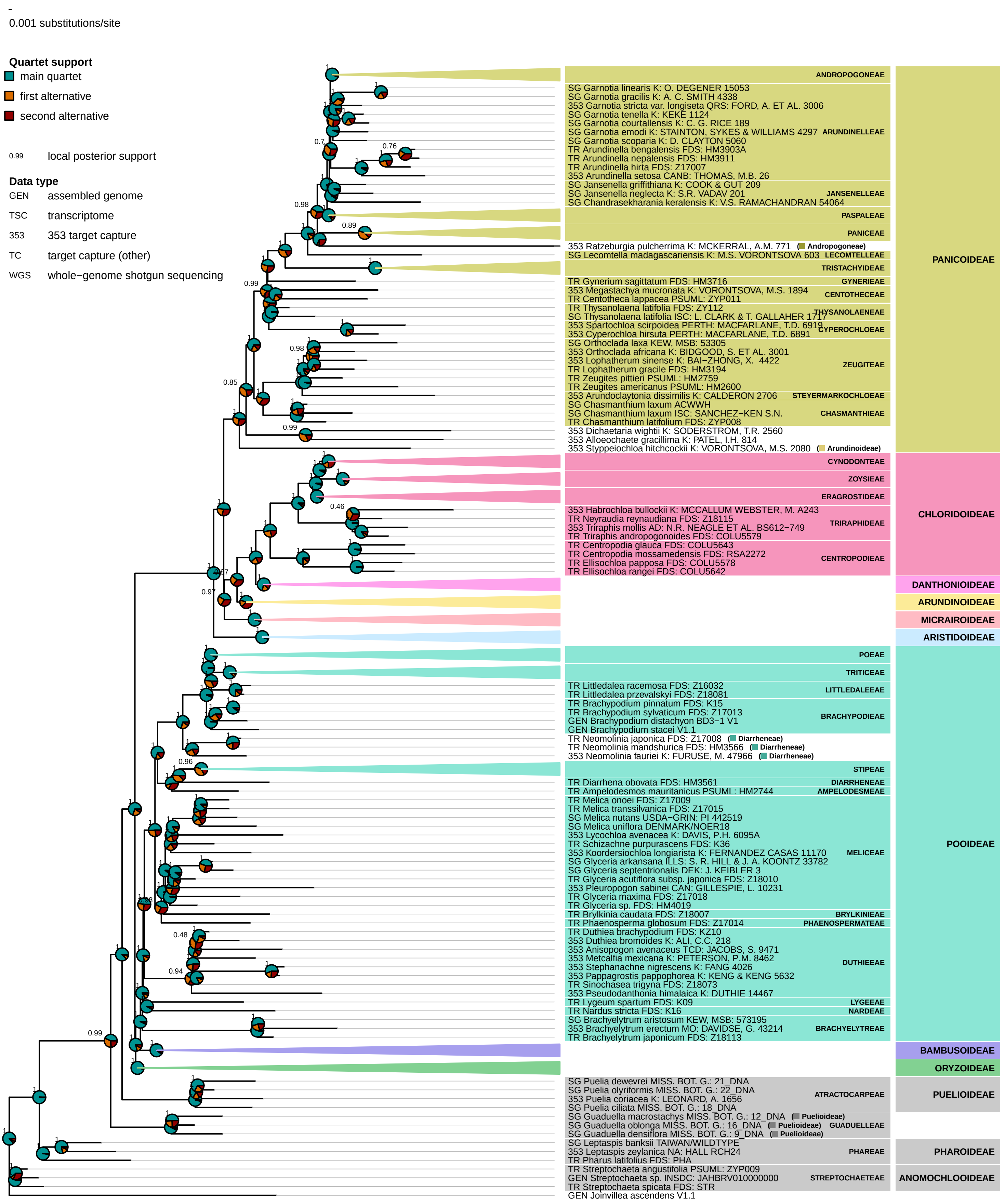

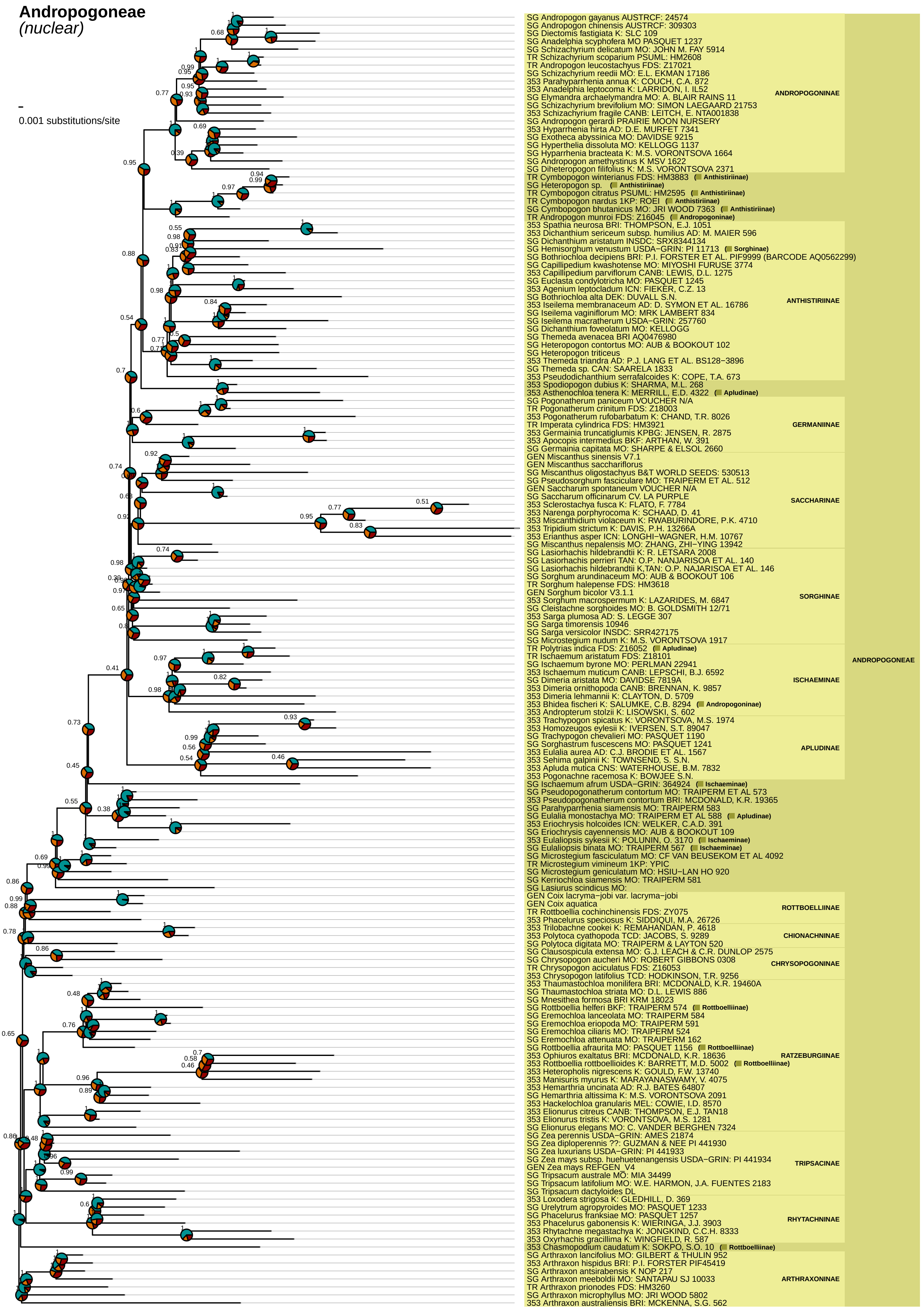
Andropogoneae
(nuclear)
0.001 substitutions/site
SG Andropogon gayanus AUSTRCF: 24574
SG Andropogon chinensis AUSTRCF: 309303
SG Diectomis fastigiata K: SLC 109
SG Anadelphia scyphofera MO PASQUET 1237
SG Schizachyrium delicatum MO: JOHN M. FAY 5914
TR Schizachyrium scoparium PSUML: HM2608
TR Andropogon leucostachyus FDS: Z17021
SG Schizachyrium reedii MO: E.L. EKMAN 17186
353 Parahyparrhenia annua K: COUCH, C.A. 872
353 Anadelphia leptocoma K: LARRIDON, I. IL52
SG Elymandra archaelymandra MO: A. BLAIR RAINS 11
SG Schizachyrium brevifolium MO: SIMON LAEGAARD 21753
353 Schizachyrium fragile CANB: LEITCH, E. NTA001838
SG Andropogon gerardi PRAIRIE MOON NURSERY
353 Hyparrhenia hirta AD: D.E. MURFET 7341
SG Exotheca abyssinica MO: DAVIDSE 9215
SG Hyperthelia dissoluta MO: KELLOGG 1137
SG Hyparrhenia bracteata K: M.S. VORONTSOVA 1664
SG Andropogon amethystinus K MSV 1622
SG Diheteropogon filifolius K: M.S. VORONTSOVA 2371
ANDROPOGONINAE
TR Cymbopogon winterianus FDS: HM3883 (Anthistiriinae)
SG Heteropogon sp. (Anthistiriinae)
TR Cymbopogon citratus PSUML: HM2595 (Anthistiriinae)
TR Cymbopogon nardus 1KP: ROEI (Anthistiriinae)
SG Cymbopogon bhutanicus MO: JRI WOOD 7363 (Anthistiriinae)
TR Andropogon munroi FDS: Z16045 (Andropogoninae)
353 Spathia neurosa BRI: THOMPSON, E.J. 1051
353 Dichanthium sericeum subsp. humilius AD: M. MAIER 596
SG Dichanthium aristatum INSDC: SRX8344134
SG Hemisorghum venustum USDA−GRIN: PI 11713 (Sorghinae)
SG Bothriochloa decipiens BRI: P.I. FORSTER ET AL. PIF9999 (BARCODE AQ0562299)
SG Capillipedium kwashotense MO: MIYOSHI FURUSE 3774
353 Capillipedium parviflorum CANB: LEWIS, D.L. 1275
SG Euclasta condylotricha MO: PASQUET 1245
353 Agenium leptocladum ICN: FIEKER, C.Z. 13
SG Bothriochloa alta DEK: DUVALL S.N.
353 Iseilema membranaceum AD: D. SYMON ET AL. 16786
SG Iseilema vaginiflorum MO: MRK LAMBERT 834
SG Iseilema macratherum USDA−GRIN: 257760
SG Dichanthium foveolatum MO: KELLOGG
SG Themeda avenacea BRI AQ0476980
SG Heteropogon contortus MO: AUB & BOOKOUT 102
SG Heteropogon triticeus
353 Themeda triandra AD: P.J. LANG ET AL. BS128−3896
SG Themeda sp. CAN: SAARELA 1833
353 Pseudodichanthium serrafalcoides K: COPE, T.A. 673
ANTHISTIRIINAE
353 Spodiopogon dubius K: SHARMA, M.L. 268
353 Asthenochloa tenera K: MERRILL, E.D. 4322 (Apludinae)
SG Pogonatherum paniceum VOUCHER N/A
TR Pogonatherum crinitum FDS: Z18003
353 Pogonatherum rufobarbatum K: CHAND, T.R. 8026
TR Imperata cylindrica FDS: HM3921
353 Germainia truncatiglumis KPBG: JENSEN, R. 2875
353 Apocopis intermedius BKF: ARTHAN, W. 391
SG Germainia capitata MO: SHARPE & ELSOL 2660
GERMAINIINAE
GEN Miscanthus sinensis V7.1
GEN Miscanthus sacchariflorus
SG Miscanthus oligostachyus B&T WORLD SEEDS: 530513
SG Pseudosorghum fasciculare MO: TRAIPERM ET AL. 512
GEN Saccharum spontaneum VOUCHER N/A
SG Saccharum officinarum CV. LA PURPLE
353 Sclerostachya fusca K: FLATO, F. 7784
353 Narenga porphyrocoma K: SCHAAD, D. 41
353 Miscanthidium violaceum K: RWABURINDORE, P.K. 4710
353 Tripidium strictum K: DAVIS, P.H. 13266A
353 Erianthus asper ICN: LONGHI−WAGNER, H.M. 10767
SG Miscanthus nepalensis MO: ZHANG, ZHI−YING 13942
SACCHARINAE
SG Lasiorhachis hildebrandtii K: R. LASTRA 2008
SG Lasiorhachis perrieri TAN: O.P. NANJARISOA ET AL. 140
SG Lasiorhachis hildebrandtii K,TAN: O.P. NAJARISOA ET AL. 146
SG Sorghum arundinaceum MO: AUB & BOOKOUT 106
TR Sorghum halepense FDS: HM3618
GEN Sorghum bicolor V3.1.1
353 Sorghum macrospermum K: LAZARIDES, M. 6847
SG Cleistachne sorghoides MO: B. GOLDSMITH 12/71
353 Sarga plumosa AD: S. LEGGE 307
SG Sarga timorensis 10946
SG Sarga versicolor INSDC: SRR427175
SG Microstegium nudum K: M.S. VORONTSOVA 1917
SORGHINAE
TR Polytrias indica FDS: Z16052 (Apludinae)
TR Ischaemum aristatum FDS: Z18101
SG Ischaemum byrone MO: PERLMAN 22941
353 Ischaemum muticum CANB: LEPSCHI, B.J. 6592
SG Dimeria aristata MO: DAVIDSE 7819A
353 Dimeria ornithopoda CANB: BRENNAN, K. 9857
353 Dimeria lehmannii K: CLAYTON, D. 5709
353 Bhidea fischeri K: SALUMKE, C.B. 8294 (Andropogoninae)
353 Andropterum stolzii K: LISOWSKI, S. 602
ISCHAEMINAE
353 Trachypogon spicatus K: VORONTSOVA, M.S. 1974
353 Homozeugos eylesii K: IVERSEN, S.T. 89047
SG Trachypogon chevalieri MO: PASQUET 1190
SG Sorghastrum fuscescens MO: PASQUET 1241
353 Eulalia aurea AD: C.J. BRODIE ET AL. 1567
353 Sehima galpinii K: TOWNSEND, S. S.N.
353 Apluda mutica CNS: WATERHOUSE, B.M. 7832
353 Pogonachne racemosa K: BOWJEE S.N.
APLUDINAE
SG Ischaemum afrum USDA−GRIN: 364924 (Ischaeminae)
SG Pseudopogonatherum contortum MO: TRAIPERM ET AL 573
353 Pseudopogonatherum contortum BRI: MCDONALD, K.R. 19365
SG Parahyparrhenia siamensis MO: TRAIPERM 583
SG Eulalia monostachya MO: TRAIPERM ET AL 588 (Apludinae)
353 Eriochrysis holcoides ICN: WELKER, C.A.D. 391
SG Eriochrysis cayennensis MO: AUB & BOOKOUT 109
353 Eulaliopsis sykesii K: POLUNIN, O. 3170 (Ischaeminae)
SG Eulaliopsis binata MO: TRAIPERM 567 (Ischaeminae)
SG Microstegium fasciculatum MO: CF VAN BEUSEKOM ET AL 4092
TR Microstegium vimineum 1KP: YPIC
SG Microstegium geniculatum MO: HSIU−LAN HO 920
SG Kerriochloa siamensis MO: TRAIPERM 581
SG Lasiurus scindicus MO:
GEN Coix lacryma−jobi var. lacryma−jobi
GEN Coix aquatica
TR Rottboellia cochinchinensis FDS: ZY075
353 Phacelurus speciosus K: SIDDIQUI, M.A. 26726
ROTTBOELLIINAE
353 Trilobachne cookei K: REMAHANDAN, P. 4618
353 Polytoca cyathopoda TCD: JACOBS, S. 9289
SG Polytoca digitata MO: TRAIPERM & LAYTON 520
CHIONACHNINAE
SG Clausospicula extensa MO: G.J. LEACH & C.R. DUNLOP 2575
SG Chrysopogon aucheri MO: ROBERT GIBBONS 0308
TR Chrysopogon aciculatus FDS: Z16053
353 Chrysopogon latifolius TCD: HODKINSON, T.R. 9256
CHRYSOPOGONINAE
353 Thaumastochloa monilifera BRI: MCDONALD, K.R. 19460A
SG Thaumastochloa striata MO: D.L. LEWIS 886
SG Mnesithea formosa BRI KRM 18023
SG Rottboellia helferi BKF: TRAIPERM 574 (Rottboelliinae)
SG Eremochloa lanceolata MO: TRAIPERM 584
SG Eremochloa eriopoda MO: TRAIPERM 591
SG Eremochloa ciliaris MO: TRAIPERM 524
SG Eremochloa attenuata MO: TRAIPERM 162
SG Rottboellia afraurita MO: PASQUET 1156 (Rottboelliinae)
353 Ophiuros exaltatus BRI: MCDONALD, K.R. 18636
353 Rottboellia rottboellioides K: BARRETT, M.D. 5002 (Rottboelliinae)
353 Heteropholis nigrescens K: GOULD, F.W. 13740
353 Manisuris myurus K: MARAYANASWAMY, V. 4075
353 Hemarthria uncinata AD: R.J. BATES 64807
SG Hemarthria altissima K: M.S. VORONTSOVA 2091
353 Hackelochloa granularis MEL: COWIE, I.D. 8570
353 Elionurus citreus CANB: THOMPSON, E.J. TAN18
353 Elionurus tristis K: VORONTSOVA, M.S. 1281
SG Elionurus elegans MO: C. VANDER BERGHEN 7324
RATZEBURGIINAE
SG Zea perennis USDA−GRIN: AMES 21874
SG Zea diploperennis ??: GUZMAN & NEE PI 441930
SG Zea luxurians USDA−GRIN: PI 441933
SG Zea mays subsp. huehuetenangensis USDA−GRIN: PI 441934
GEN Zea mays REFGEN_V4
SG Tripsacum australe MO: MIA 34499
SG Tripsacum latifolium MO: W.E. HARMON, J.A. FRENCH 2183
SG Tripsacum dactyloides DL
TRIPSACINAE
353 Loxodera strigosa K: GLEDHILL, D. 369
SG Urelytrum agropyroides MO: PASQUET 1233
SG Phacelurus franksiae MO: PASQUET 1257
353 Phacelurus gabonensis K: WIERINGA, J.J. 3903
353 Rhytachne megastachya K: JONGKIND, C.C.H. 8333
353 Oxyrhachis gracillima K: WINGFIELD, R. 587
RHYTACHNINAE
353 Chasmopodium caudatum K: SOKPO, S.O. 10 (Rottboelliinae)
SG Arthraxon lancifolius MO: GILBERT & THULIN 952
353 Arthraxon hispidus BRI: P.I. FORSTER PIF45419
SG Arthraxon antsirabensis K NOP 217
SG Arthraxon meeboldii MO: SANTAPAU SJ 10033
TR Arthraxon prionodes FDS: HM3260
SG Arthraxon microphyllus MO: JRI WOOD 5802
353 Arthraxon australiensis BRI: MCKENNA, S.G. 562
ARTHRAXONINAE
ANDROPOGONEAE

**Paspaleae**
*(nuclear)*

0.001 substitutions/site

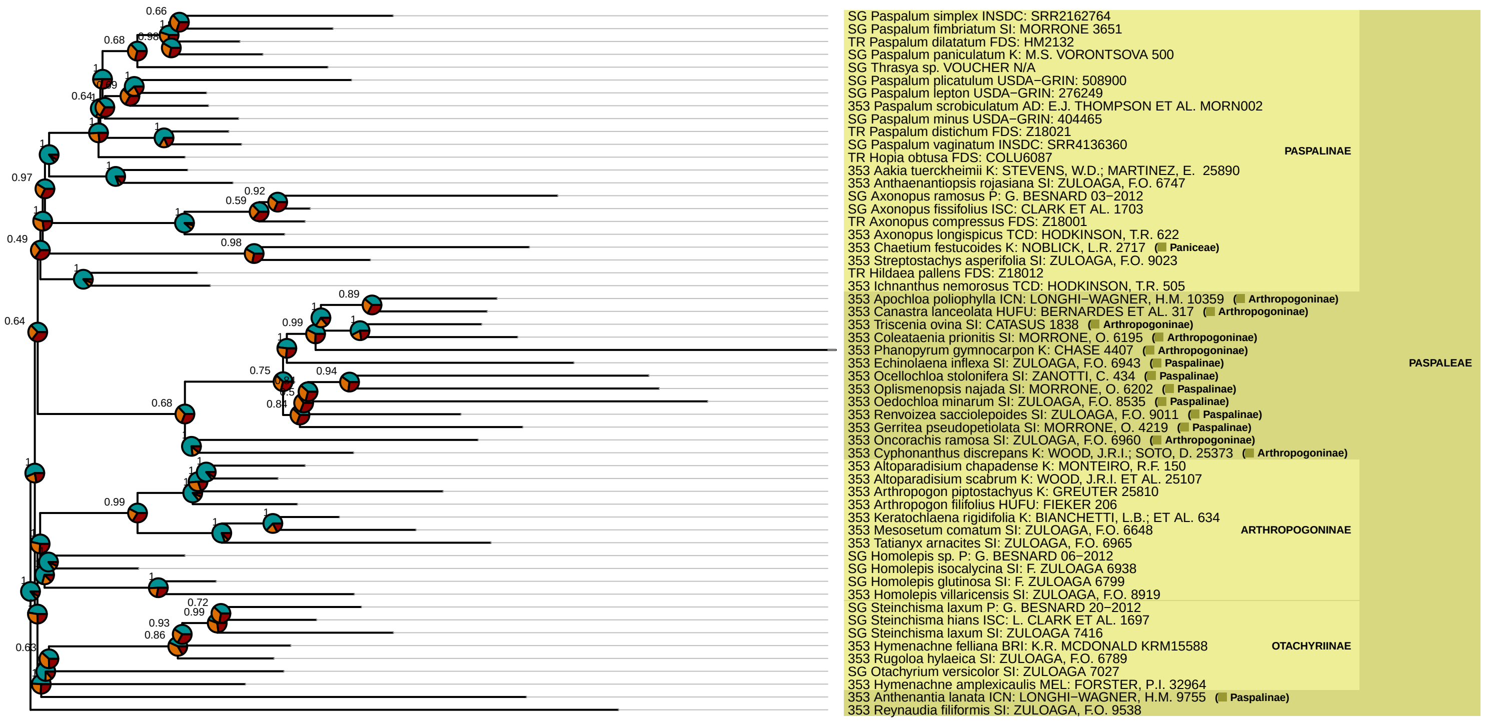

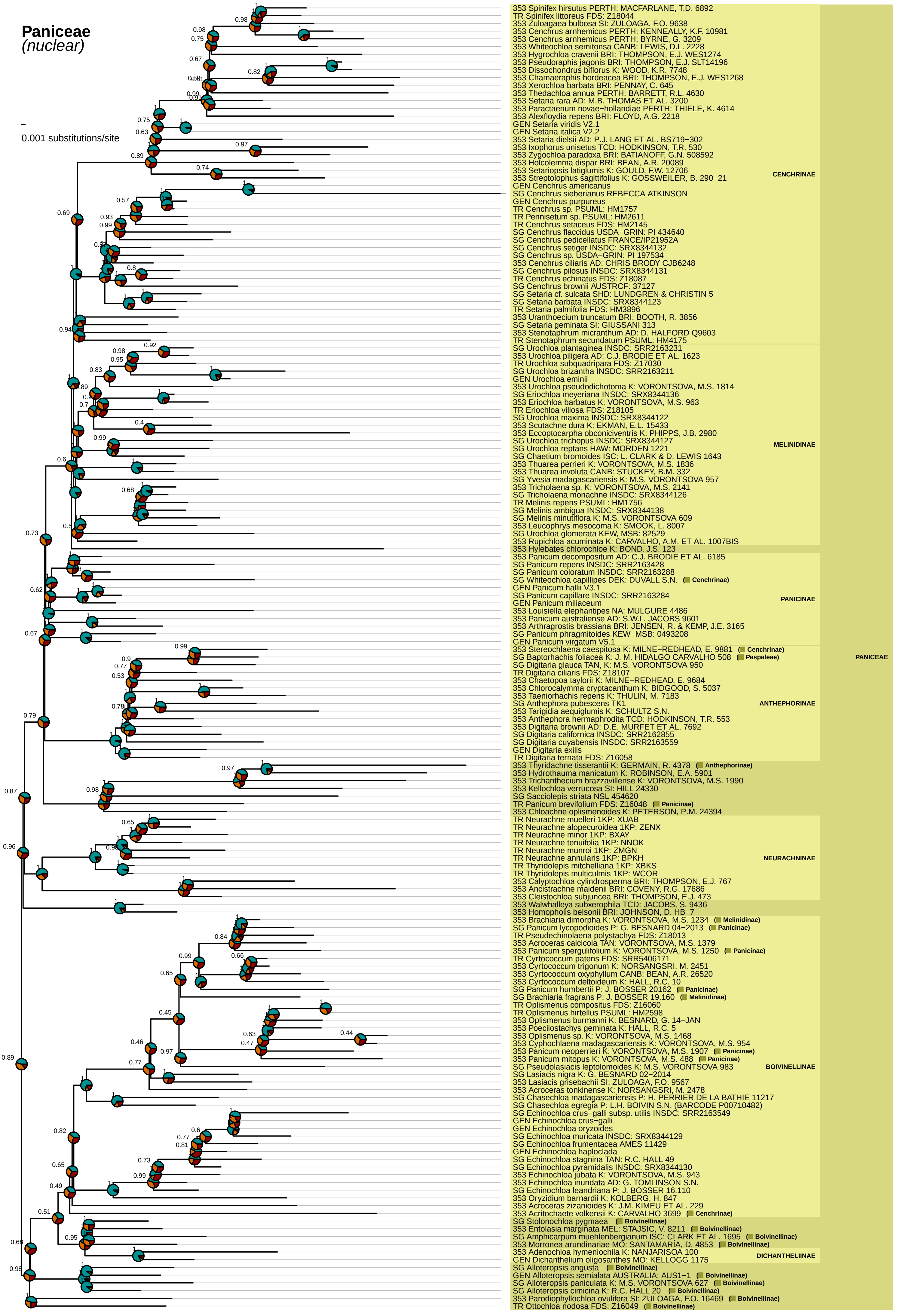
Paniceae
(nuclear)
0.001 substitutions/site
353 Spinifex hirsutus PERTH: MACFARLANE, T.D. 6892
TR Spinifex littoreus FDS: Z18044
353 Zuloagaea bulbosa SI: ZULOAGA, F.O. 9638
353 Cenchrus arnhemicus PERTH: KENNEALLY, K.F. 10981
353 Cenchrus arnhemicus PERTH: BYRNE, G. 3209
353 Whiteochloa semitonsa CANB: LEWIS, D.L. 2228
353 Hygrochloa cravenii BRI: THOMPSON, E.J. WES1274
353 Pseudoraphis jagonis BRI: THOMPSON, E.J. SLT14196
353 Dissochondrus biflorus K: WOOD, K.R. 7748
353 Chamaeraphis hordeacea BRI: THOMPSON, E.J. WES1268
353 Xerochloa barbata BRI: PENNAY, C. 645
353 Thedachloa annua PERTH: BARRETT, R.L. 4630
353 Setaria rara AD: M.B. THOMAS ET AL. 3200
353 Paractaenum novae−hollandiae PERTH: THIELE, K. 4614
353 Alexfloydia repens BRI: FLOYD, A.G. 2218
GEN Setaria viridis V2.1
GEN Setaria italica V2.2
353 Setaria dielsii AD: P.J. LANG ET AL. BS719−302
353 Ixophorus unisetus TCD: HODKINSON, T.R. 530
353 Zygochloa paradoxa BRI: BATIANOFF, G.N. 508592
353 Holcolemma dispar BRI: BEAN, A.R. 20089
353 Setariopsis latiglumis K: GOULD, F.W. 12706
353 Streptolophus sagittifolius K: GOSSWEILER, B. 290−21
GEN Cenchrus americanus
SG Cenchrus sieberianus REBECCA ATKINSON
GEN Cenchrus purpureus
TR Cenchrus sp. PSUML: HM1757
TR Pennisetum sp. PSUML: HM2611
TR Cenchrus setaceus FDS: HM2145
SG Cenchrus flaccidus USDA−GRIN: PI 434640
SG Cenchrus pedicellatus FRANCE/IP21952A
SG Cenchrus setiger INSDC: SRX8344132
SG Cenchrus sp. USDA−GRIN: PI 197534
353 Cenchrus ciliaris AD: CHRIS BRODY CJB6248
SG Cenchrus pilosus INSDC: SRX8344131
TR Cenchrus echinatus FDS: Z18087
SG Cenchrus brownii AUSTRCF: 37127
SG Setaria cf. sulcata SHD: LUNDGREN & CHRISTIN 5
SG Setaria barbata INSDC: SRX8344123
TR Setaria palmifolia FDS: HM3896
353 Uranthoecium truncatum BRI: BOOTH, R. 3856
SG Setaria geminata SI: GIUSSANI 313
353 Stenotaphrum micranthum AD: D. HALFORD Q9603
TR Stenotaphrum secundatum PSUML: HM4175
CENCHRINAE
SG Urochloa plantaginea INSDC: SRR2163231
353 Urochloa piligera AD: C.J. BRODIE ET AL. 1623
TR Urochloa subquadripara FDS: Z17030
SG Urochloa brizantha INSDC: SRR2163211
GEN Urochloa eminii
353 Urochloa pseudodichotoma K: VORONTSOVA, M.S. 1814
SG Eriochloa meyeriana INSDC: SRX8344136
353 Eriochloa barbatus K: VORONTSOVA, M.S. 963
TR Eriochloa villosa FDS: Z18105
SG Urochloa maxima INSDC: SRX8344122
353 Scutachne dura K: EKMAN, E.L. 15433
353 Eccoptocarpha obconiciventris K: PHIPPS, J.B. 2980
SG Urochloa trichopus INSDC: SRX8344127
SG Urochloa reptans HAW: MORDEN 1221
SG Chaetium bromoides ISC: L. CLARK & D. LEWIS 1643
353 Thuarea perrieri K: VORONTSOVA, M.S. 1836
353 Thuarea involuta CANB: STUCKEY, B.M. 332
SG Yvesia madagascariensis K: M.S. VORONTSOVA 957
353 Tricholaena sp. K: VORONTSOVA, M.S. 2141
SG Tricholaena monachne INSDC: SRX8344126
TR Melinis repens PSUML: HM1756
SG Melinis ambigua INSDC: SRX8344138
SG Melinis minutiflora K: M.S. VORONTSOVA 609
353 Leucophrys mesocoma K: SMOOK, L. 8007
SG Urochloa glomerata KEW, MSB: 82529
353 Rupichloa acuminata K: CARVALHO, A.M. ET AL. 1007BIS
MELINIDINAE
353 Hylebates chlorochloe K: BOND, J.S. 123
353 Panicum decompositum AD: C.J. BRODIE ET AL. 6185
SG Panicum repens INSDC: SRR2163428
SG Panicum coloratum INSDC: SRR2163288
SG Whiteochloa capillipes DEK: DUVALL S.N. (Cenchrinae)
GEN Panicum hallii V3.1
SG Panicum capillare INSDC: SRR2163284
GEN Panicum miliaceum
353 Louisiella elephantipes NA: MULGURE 4486
353 Panicum australiense AD: S.W.L. JACOBS 9601
353 Arthragrostis brassiana BRI: JENSEN, R. & KEMP, J.E. 3165
SG Panicum phragmitoides KEW−MSB: 0493208
GEN Panicum virgatum V5.1
PANICINAE
353 Stereochlaena caespitosa K: MILNE−REDHEAD, E. 9881 (Cenchrinae)
SG Baptorhachis foliacea K: J. M. HIDALGO CARVALHO 508 (Paspaleae)
SG Digitaria glauca TAN, K: M.S. VORONTSOVA 950
TR Digitaria ciliaris FDS: Z18107
353 Chaetopoa taylorii K: MILNE−REDHEAD, E. 9684
353 Chlorocalymma cryptacanthum K: BIDGOOD, S. 5037
353 Taeniorhachis repens K: THULIN, M. 7183
SG Anthephora pubescens TK1
353 Tarigidia aequiglumis K: CARVALHO S.N.
353 Anthephora hermaphrodita TCD: HODKINSON, T.R. 553
353 Digitaria brownii AD: D.E. MURFET ET AL. 7692
SG Digitaria californica INSDC: SRR2162855
SG Digitaria cuyabensis INSDC: SRR2163559
GEN Digitaria exilis
TR Digitaria ternata FDS: Z16058
ANTHEPHORINAE
353 Thyridachne tisserantii K: GERMAIN, R. 4378 (Anthephorinae)
353 Hydrothauma manicatum K: ROBINSON, E.A. 5901
353 Trichanthecium brazzavillense K: VORONTSOVA, M.S. 1990
353 Kellochloa verrucosa SI: HILL 24330
SG Sacciolepis striata NSL 454620
TR Panicum brevifolium FDS: Z16048 (Panicinae)
353 Chloachne oplismenoides K: PETERSON, P.M. 24394
TR Neurachne muelleri 1KP: XUAB
TR Neurachne alopecuroidea 1KP: ZENX
TR Neurachne minor 1KP: BXAY
TR Neurachne tenuifolia 1KP: NNOK
TR Neurachne munroi 1KP: ZMGN
TR Neurachne annularis 1KP: BPKH
TR Thyridolepis mitchelliana 1KP: XBKS
TR Thyridolepis multiculmis 1KP: WCOR
353 Calyptochloa cylindrosperma BRI: THOMPSON, E.J. 767
353 Ancistrachne maidenii BRI: COVENY, R.G. 17686
353 Cleistochloa subjuncea BRI: THOMPSON, E.J. 473
NEURACHNINAE
353 Walwhalleya subxerophila TCD: JACOBS, S. 9436
353 Homopholis belsonii BRI: JOHNSON, D. HB−7
353 Brachiaria dimorpha K: VORONTSOVA, M.S. 1234 (Melinidinae)
SG Panicum lycopodioides P: G. BESNARD 04−2013 (Panicinae)
TR Pseudechinolaena polystachya FDS: Z18013
353 Acroceras calcicola TAN: VORONTSOVA, M.S. 1379
353 Panicum spergulifolium K: VORONTSOVA, M.S. 1250 (Panicinae)
TR Cyrtococcum patens FDS: SRR5406171
353 Cyrtococcum trigonum K: NORSANGSRI, M. 2451
353 Cyrtococcum oxyphyllum CANB: BEAN, A.R. 26520
353 Cyrtococcum deltoideum K: HALL, R.C. 10
SG Panicum humbertii P: J. BOSSER 20162 (Panicinae)
SG Brachiaria fragrans P: J. BOSSER 19.160 (Melinidinae)
TR Oplismenus compositus FDS: Z16060
TR Oplismenus hirtellus PSUML: HM2598
353 Oplismenus burmanni K: BESNARD, G. 14−JAN
353 Poecilostachys geminata K: HALL, R.C. 5
353 Oplismenus sp. K: VORONTSOVA, M.S. 1468
353 Cyphochlaena madagascariensis K: VORONTSOVA, M.S. 954
353 Panicum neoperrieri K: VORONTSOVA, M.S. 1907 (Panicinae)
353 Panicum mitopus K: VORONTSOVA, M.S. 488 (Panicinae)
SG Pseudolasiacis leptolomoides K: M.S. VORONTSOVA 983
SG Lasiacis nigra K: G. BESNARD 02−2014
353 Lasiacis grisebachii SI: ZULOAGA, F.O. 9567
353 Acroceras tonkinense K: NORSANGSRI, M. 2478
SG Chasechloa madagascariensis P: H. PERRIER DE LA BATHIE 11217
SG Chasechloa egregia P: L.H. BOIVIN S.N. (BARCODE P00710482)
SG Echinochloa crus−galli subsp. utilis INSDC: SRR2163549
GEN Echinochloa crus−galli
GEN Echinochloa oryzoides
SG Echinochloa muricata INSDC: SRX8344129
SG Echinochloa frumentacea AMES 11429
GEN Echinochloa haploclada
SG Echinochloa stagnina TAN: R.C. HALL 49
SG Echinochloa pyramidalis INSDC: SRX8344130
353 Echinochloa jubata K: VORONTSOVA, M.S. 943
353 Echinochloa inundata AD: G. TOMLINSON S.N.
SG Echinochloa leandriana P: J. BOSSER 16.110
353 Oryzidium barnardii K: KOLBERG, H. 847
353 Acroceras zizanioides K: J.M. KIMEU ET AL. 229
353 Acritochaete volkensii K: CARVALHO 3699 (Cenchrinae)
BOIVINELLINAE
SG Stolonochloa pygmaea (Boivinellinae)
353 Entolasia marginata MEL: STAJSIC, V. 8211 (Boivinellinae)
SG Amphicarpum muehlenbergianum ISC: CLARK ET AL. 1695 (Boivinellinae)
353 Morronea arundinariae MO: SANTAMARIA, D. 4853 (Boivinellinae)
353 Adenochloa hymeniochila K: NANJARISOA 100
GEN Dichanthelium oligosanthes MO: KELLOGG 1175
DICHANTHELIINAE
SG Alloteropsis angusta (Boivinellinae)
GEN Alloteropsis semialata AUSTRALIA: AUS1−1 (Boivinellinae)
SG Alloteropsis paniculata K: M.S. VORONTSOVA 627 (Boivinellinae)
SG Alloteropsis cimicina K: R.C. HALL 20 (Boivinellinae)
353 Parodiophyllochloa ovulifera SI: ZULOAGA, F.O. 16469 (Boivinellinae)
TR Ottochloa nodosa FDS: Z16049 (Boivinellinae)
PANICEAE

### **Tristachyideae**
#### *(nuclear)*

0.001 substitutions/site

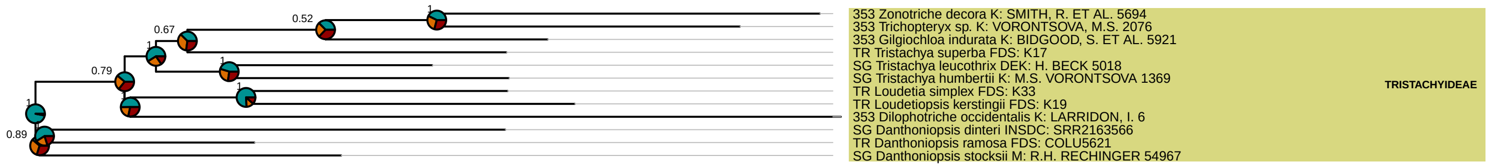

**Cynodonteae**
*(nuclear)*

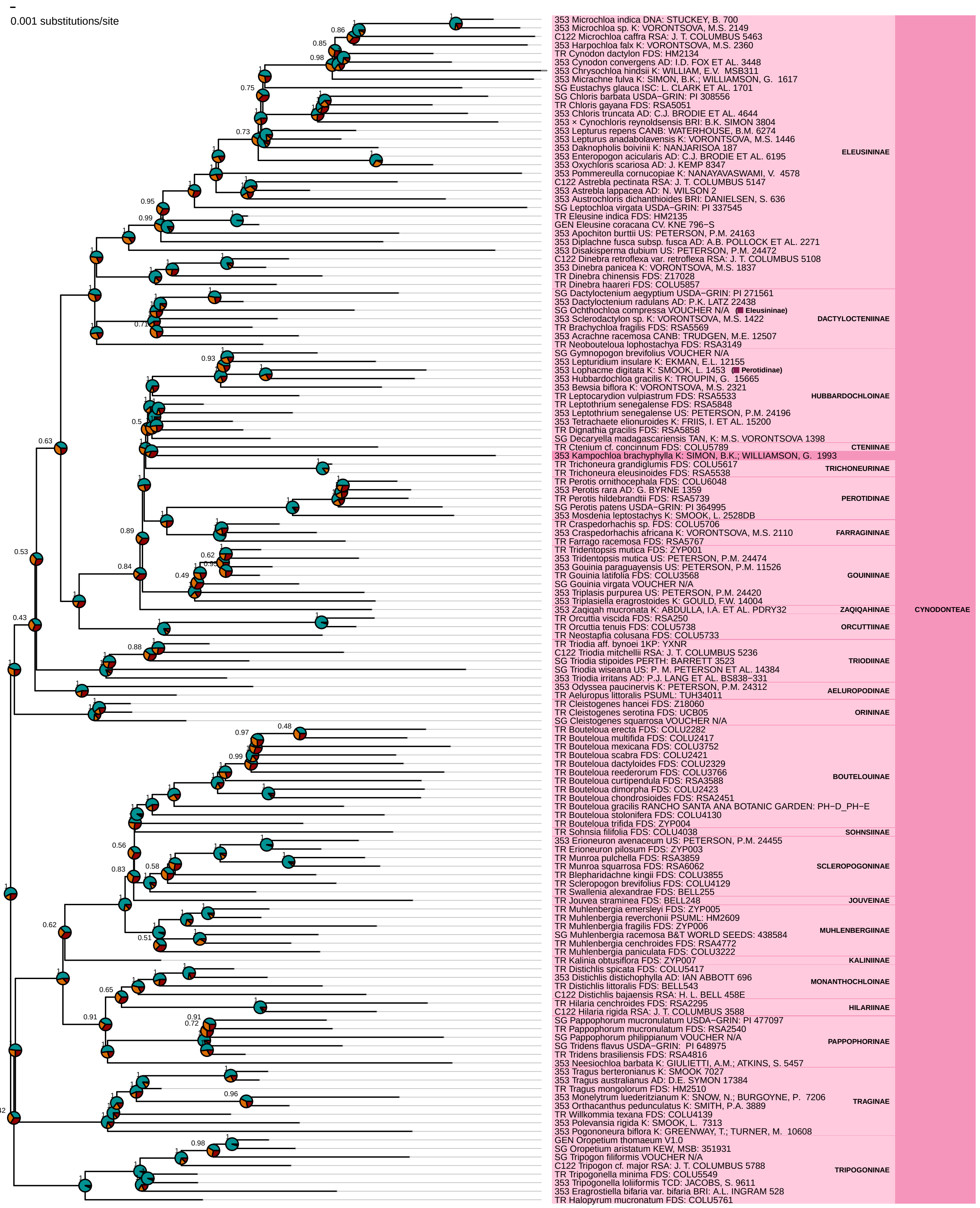

**Zoysieae**
*(nuclear)*

0.001 substitutions/site

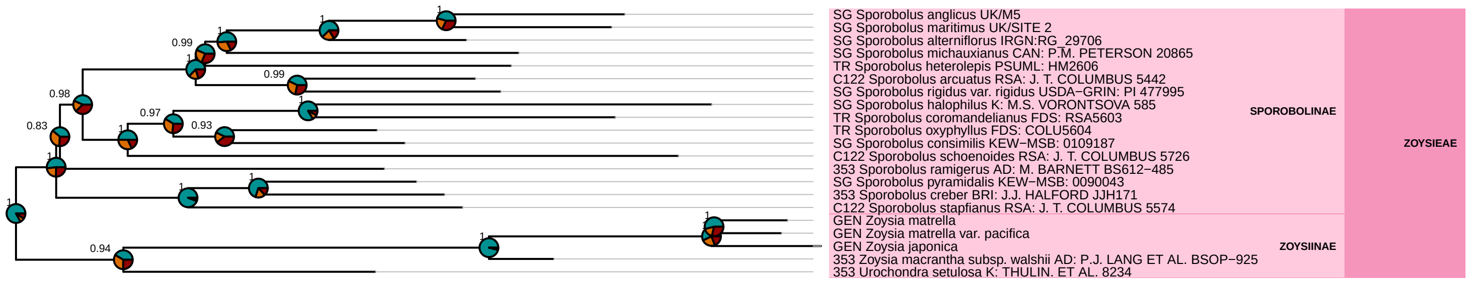

**Eragrostideae**
*(nuclear)*

0.001 substitutions/site

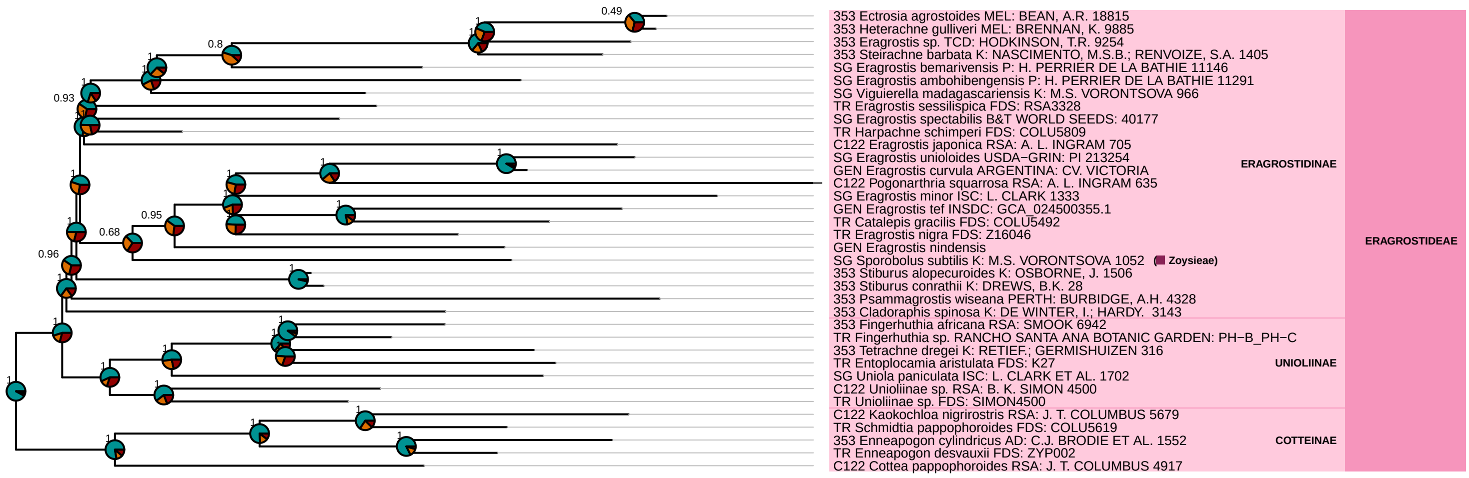

**Danthonioideae**
*(nuclear)*

0.001 substitutions/site

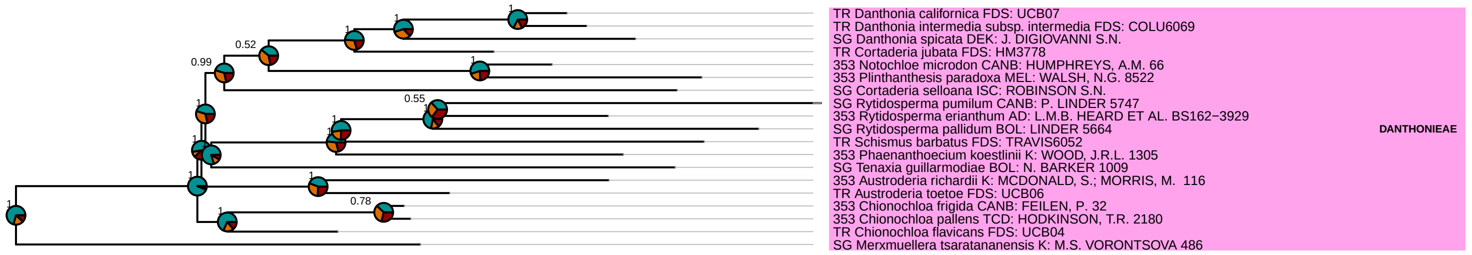

**Arundinoideae**
*(nuclear)*

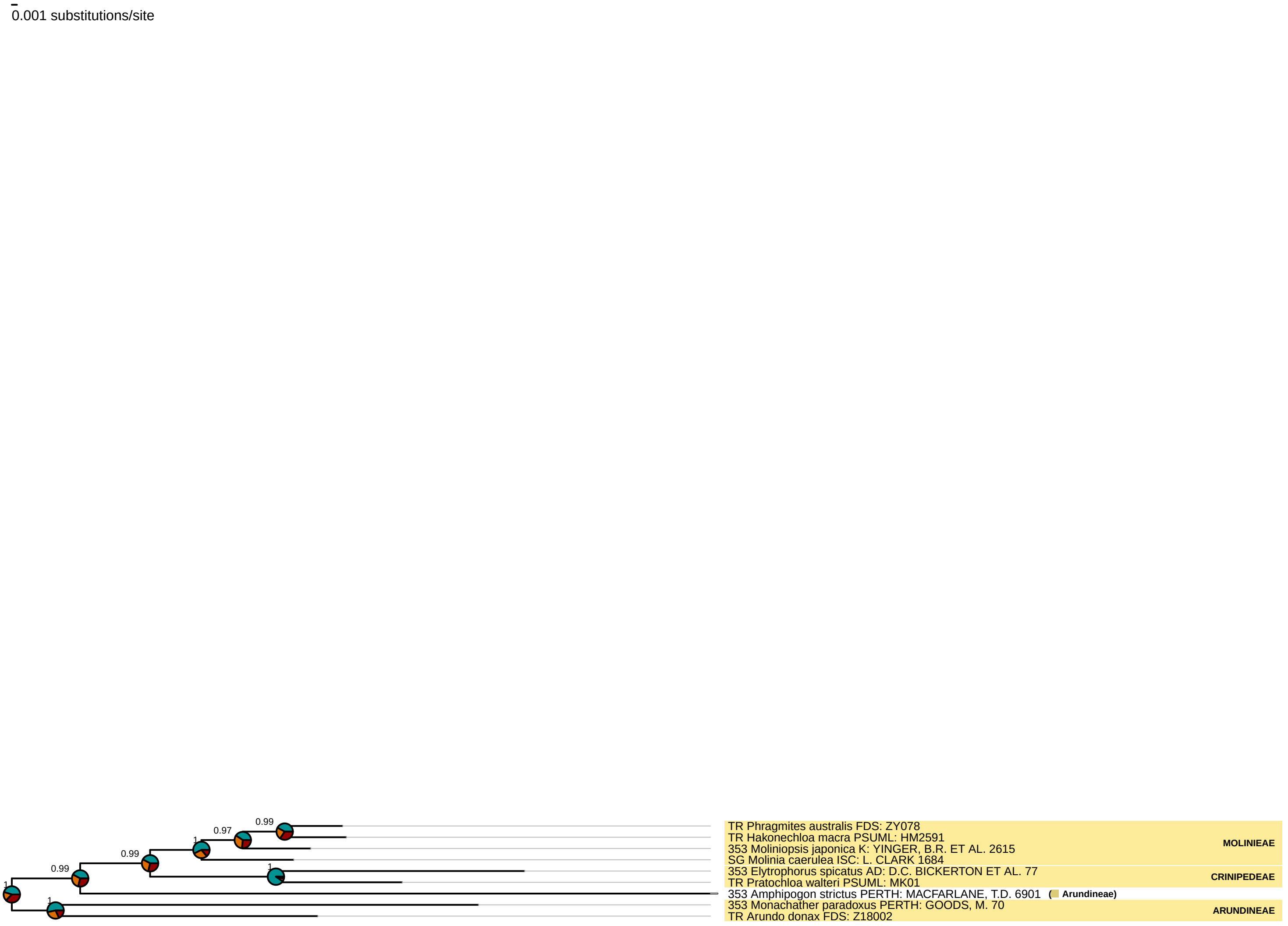

**Micrairoideae**
*(nuclear)*

0.001 substitutions/site

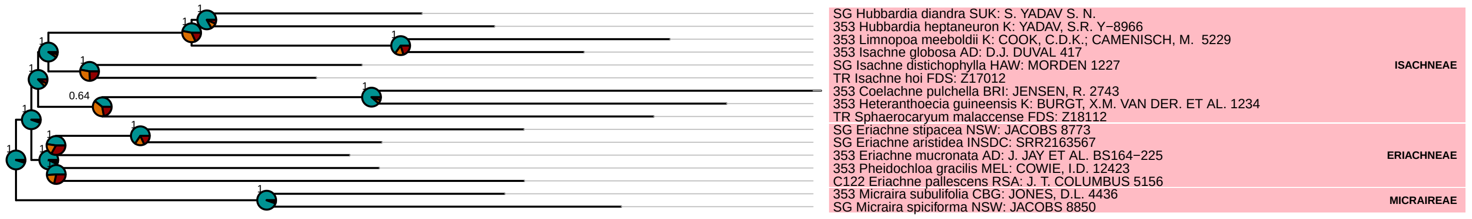

### Aristidoideae
*(nuclear)*

0.001 substitutions/site

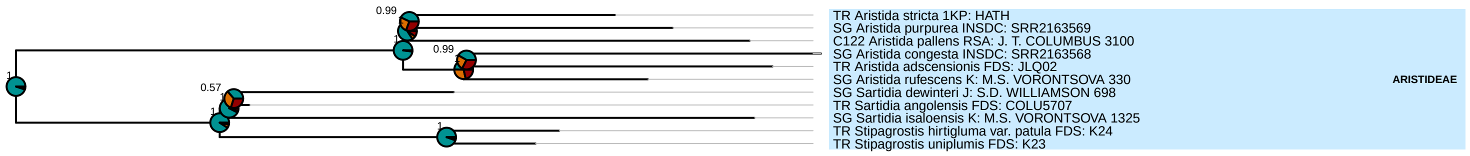

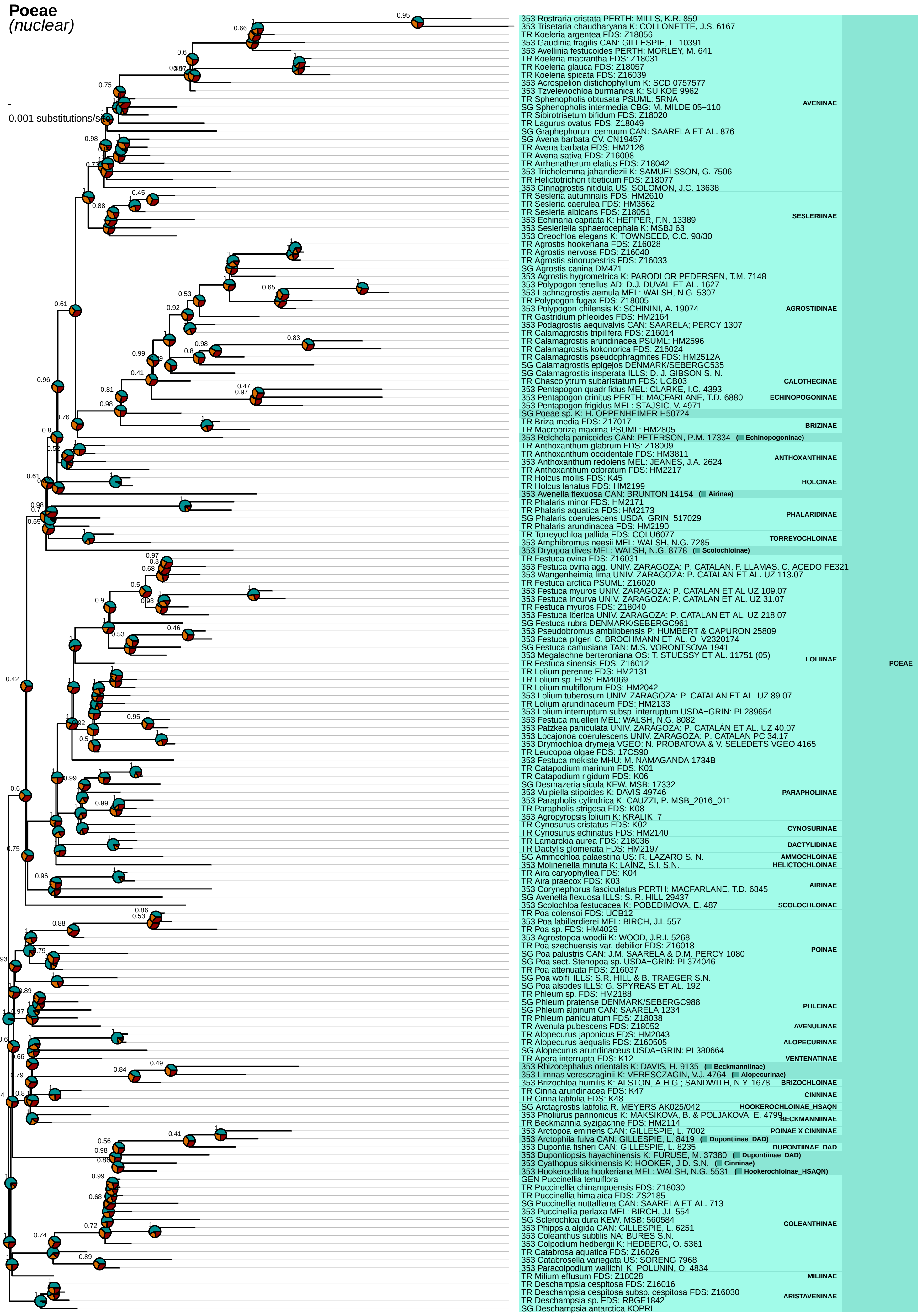

Poeae
(nuclear)
0.001 substitutions/site
353 Rostraria cristata PERTH: MILLS, K.R. 859
353 Trisetaria chaudharyana K: COLLONETTE, J.S. 6167
TR Koeleria argentea FDS: Z18056
353 Gaudinia fragilis CAN: GILLESPIE, L. 10391
353 Avellinia festucoides PERTH: MORLEY, M. 641
TR Koeleria macrantha FDS: Z18031
TR Koeleria glauca FDS: Z18057
TR Koeleria spicata FDS: Z16039
353 Acrospelion distichophyllum K: SCD 0757577
353 Tzveleviochloa burmanica K: SU KOE 9962
TR Sphenopholis obtusata PSUML: 5RNA
SG Sphenopholis intermedia CBG: M. MILDE 05−110
TR Sibirotrisetum bifidum FDS: Z18020
TR Lagurus ovatus FDS: Z18049
SG Graphephorum cernuum CAN: SAARELA ET AL. 876
SG Avena barbata CV. CN19457
TR Avena barbata FDS: HM2126
TR Avena sativa FDS: Z16008
TR Arrhenatherum elatius FDS: Z18042
353 Tricholemma jahandiezii K: SAMUELSSON, G. 7506
TR Helictotrichon tibeticum FDS: Z18077
353 Cinnagrostis nitidula US: SOLOMON, J.C. 13638
AVENINAE
TR Sesleria autumnalis FDS: HM2610
TR Sesleria caerulea FDS: HM3562
TR Sesleria albicans FDS: Z18051
353 Echinaria capitata K: HEPPER, F.N. 13389
353 Sesleriella sphaerocephala K: MSBJ 63
353 Oreochloa elegans K: TOWNSEED, C.C. 98/30
SESLERIINAE
TR Agrostis hookeriana FDS: Z16028
TR Agrostis nervosa FDS: Z16040
TR Agrostis sinorupestris FDS: Z16033
SG Agrostis canina DM471
353 Agrostis hygrometrica K: PARODI OR PEDERSEN, T.M. 7148
353 Polypogon tenellus AD: D.J. DUVAL ET AL. 1627
353 Lachnagrostis aemula MEL: WALSH, N.G. 5307
TR Polypogon fugax FDS: Z18005
353 Polypogon chilensis K: SCHININI, A. 19074
TR Gastridium phleoides FDS: HM2164
353 Podagrostis aequivalvis CAN: SAARELA; PERCY 1307
TR Calamagrostis tripilifera FDS: Z16014
TR Calamagrostis arundinacea PSUML: HM2596
TR Calamagrostis kokonorica FDS: Z16024
TR Calamagrostis pseudophragmites FDS: HM2512A
SG Calamagrostis epigejos DENMARK/SEBERGC535
SG Calamagrostis insperata ILLS: D. J. GIBSON S. N.
AGROSTIDINAE
TR Chascolytrum subaristatum FDS: UCB03
CALOTHECINAE
353 Pentapogon quadrifidus MEL: CLARKE, I.C. 4393
353 Pentapogon crinitus PERTH: MACFARLANE, T.D. 6880
353 Pentapogon frigidus MEL: STAJSIC, V. 4971
ECHINOPOGONINAE
SG Poeae sp. K: H. OPPENHEIMER H50724
TR Briza media FDS: Z17017
TR Macrobriza maxima PSUML: HM2805
BRIZINAE
353 Relchela panicoides CAN: PETERSON, P.M. 17334 (Echinopogoninae)
TR Anthoxanthum glabrum FDS: Z18009
TR Anthoxanthum occidentale FDS: HM3811
353 Anthoxanthum redolens MEL: JEANES, J.A. 2624
TR Anthoxanthum odoratum FDS: HM2217
ANTHOXANTHINAE
TR Holcus mollis FDS: K45
TR Holcus lanatus FDS: HM2199
HOLCINAE
353 Avenella flexuosa CAN: BRUNTON 14154 (Airinae)
TR Phalaris minor FDS: HM2171
TR Phalaris aquatica FDS: HM2173
SG Phalaris coerulescens USDA−GRIN: 517029
TR Phalaris arundinacea FDS: HM2190
PHALARIDINAE
TR Torreyochloa pallida FDS: COLU6077
353 Amphibromus neesii MEL: WALSH, N.G. 7285
TORREYOCHLOINAE
353 Dryopoa dives MEL: WALSH, N.G. 8778 (Scolochloinae)
TR Festuca ovina FDS: Z16031
353 Festuca ovina agg. UNIV. ZARAGOZA: P. CATALAN, F. LLAMAS, C. ACEDO FE321
353 Wangenheimia lima UNIV. ZARAGOZA: P. CATALAN ET AL. UZ 113.07
TR Festuca arctica PSUML: Z16020
353 Festuca myuros UNIV. ZARAGOZA: P. CATALAN ET AL UZ 109.07
353 Festuca incurva UNIV. ZARAGOZA: P. CATALAN ET AL. UZ 31.07
TR Festuca myuros FDS: Z18040
353 Festuca iberica UNIV. ZARAGOZA: P. CATALAN ET AL. UZ 218.07
SG Festuca rubra DENMARK/SEBERGC961
353 Pseudobromus ambilobensis P: HUMBERT & CAPURON 25809
353 Festuca pilgeri C. BROCHMANN ET AL. O−V2320174
SG Festuca camusiana TAN: M.S. VORONTSOVA 1941
353 Megalachne berteroniana OS: T. STUESSY ET AL. 11751 (05)
TR Festuca sinensis FDS: Z16012
TR Lolium perenne FDS: HM2131
TR Lolium sp. FDS: HM4069
TR Lolium multiflorum FDS: HM2042
353 Lolium tuberosum UNIV. ZARAGOZA: P. CATALAN ET AL. UZ 89.07
TR Lolium arundinaceum FDS: HM2133
353 Lolium interruptum subsp. interruptum USDA−GRIN: PI 289654
353 Festuca muelleri MEL: WALSH, N.G. 8082
353 Patzkea paniculata UNIV. ZARAGOZA: P. CATALÁN ET AL. UZ 40.07
353 Locajonoa coerulescens UNIV. ZARAGOZA: P. CATALAN PC 34.17
353 Drymochloa drymeja VGEO: N. PROBATOVA & V. SELEDETS VGEO 4165
TR Leucopoa olgae FDS: 17CS90
353 Festuca mekiste MHU: M. NAMAGANDA 1734B
LOLIINAE
TR Catapodium marinum FDS: K01
TR Catapodium rigidum FDS: K06
SG Desmazeria sicula KEW, MSB: 17332
353 Vulpiella stipoides K: DAVIS 49746
353 Parapholis cylindrica K: CAUZZI, P. MSB_2016_011
TR Parapholis strigosa FDS: K08
353 Agropyropsis lolium K: KRALIK 7
PARAPHOLIINAE
TR Cynosurus cristatus FDS: K02
TR Cynosurus echinatus FDS: HM2140
CYNOSURINAE
TR Lamarckia aurea FDS: Z18036
TR Dactylis glomerata FDS: HM2197
DACTYLIDINAE
SG Ammochloa palaestina US: R. LAZARO S. N.
AMMOCHLOINAE
353 Molineriella minuta K: LAINZ, S.I. S.N.
HELICTOCHLOINAE
TR Aira caryophyllea FDS: K04
TR Aira praecox FDS: K03
353 Corynephorus fasciculatus PERTH: MACFARLANE, T.D. 6845
SG Avenella flexuosa ILLS: S. R. HILL 29437
AIRINAE
353 Scolochloa festucacea K: POBEDIMOVA, E. 487
SCOLOCHLOINAE
TR Poa colensoi FDS: UCB12
353 Poa labillardierei MEL: BIRCH, J.L 557
TR Poa sp. FDS: HM4029
353 Agrostopoa woodii K: WOOD, J.R.I. 5268
TR Poa szechuensis var. debilior FDS: Z16018
SG Poa palustris CAN: J.M. SAARELA & D.M. PERCY 1080
SG Poa sect. Stenopoa sp. USDA−GRIN: PI 374066
TR Poa attenuata FDS: Z16037
SG Poa wolfii ILLS: S.R. HILL & B. TRAEGER S.N.
SG Poa alsodes ILLS: G. SPYREAS ET AL. 192
POINAE
TR Phleum sp. FDS: HM2188
SG Phleum pratense DENMARK/SEBERGC988
SG Phleum alpinum CAN: SAARELA 1234
TR Phleum paniculatum FDS: Z18038
PHLEINAE
TR Avenula pubescens FDS: Z18052
AVENULINAE
TR Alopecurus japonicus FDS: HM2043
TR Alopecurus aequalis FDS: Z160505
SG Alopecurus arundinaceus USDA−GRIN: PI 380664
ALOPECURINAE
TR Apera interrupta FDS: K12
VENTENATINAE
353 Rhizocephalus orientalis K: DAVIS, H. 9135 (Beckmanniinae)
353 Limnas veresczaginii K: VERESCZAGIN, V.J. 4764 (Alopecurinae)
353 Brizochloa humilis K: ALSTON, A.H.G.; SANDWITH, N.Y. 1678
BRIZOCHLOINAE
TR Cinna arundinacea FDS: K47
TR Cinna latifolia FDS: K48
CINNINAE
SG Arctagrostis latifolia R. MEYERS AK025/042
HOOKEROCHLOINAE_HSAQN
353 Pholiurus pannonicus K: MAKSIKOVA, B. & POLJAKOVA, E. 4799
TR Beckmannia syzigachne FDS: HM2114
BECKMANNIINAE
353 Arctopoa eminens CAN: GILLESPIE, L. 7002
POINAE X CINNINAE
353 Arctophila fulva CAN: GILLESPIE, L. 8419 (Dupontiinae_DAD)
353 Dupontia fisheri CAN: GILLESPIE, L. 8235
DUPONTIINAE_DAD
353 Dupontiopsis hayachinensis K: FURUSE, M. 37380 (Dupontiinae_DAD)
353 Cyathopus sikkimensis K: HOOKER, J.D. S.N. (Cinninae)
353 Hookerochloa hookeriana MEL: WALSH, N.G. 5531 (Hookerochloinae_HSAQN)
GEN Puccinellia tenuiflora
TR Puccinellia chinampoensis FDS: Z18030
TR Puccinellia himalaica FDS: ZS2185
SG Puccinellia nuttalliana CAN: SAARELA ET AL. 713
353 Puccinellia perlaxa MEL: BIRCH, J.L 554
SG Sclerochloa dura KEW, MSB: 560584
353 Phippsia algida CAN: GILLESPIE, L. 6251
353 Coleanthus subtilis NA: BURES S.N.
353 Colpodium hedbergii K: HEDBERG, O. 5361
TR Catabrosa aquatica FDS: Z16026
353 Catabrosella variegata US: SORENG 7968
353 Paracolpodium wallichii K: POLUNIN, O. 4834
COLEANTHINAE
TR Milium effusum FDS: Z18028
MILIINAE
TR Deschampsia cespitosa FDS: Z16016
TR Deschampsia cespitosa subsp. cespitosa FDS: Z16030
TR Deschampsia sp. FDS: RBGE1842
SG Deschampsia antarctica KOPRI
ARISTAVENINAE
POEAE

**Triticeae**
*(nuclear)*

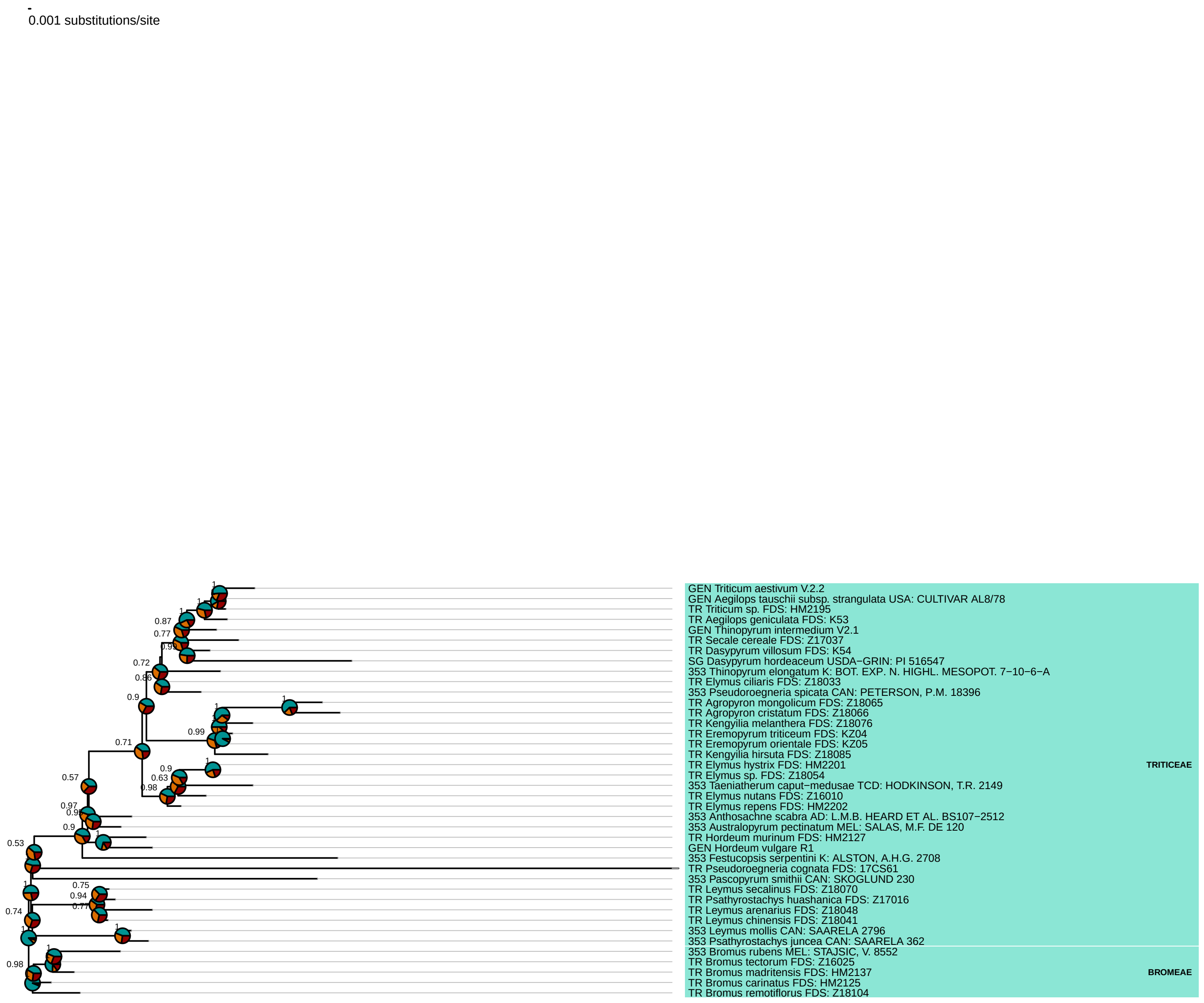

**Stipeae**
*(nuclear)*

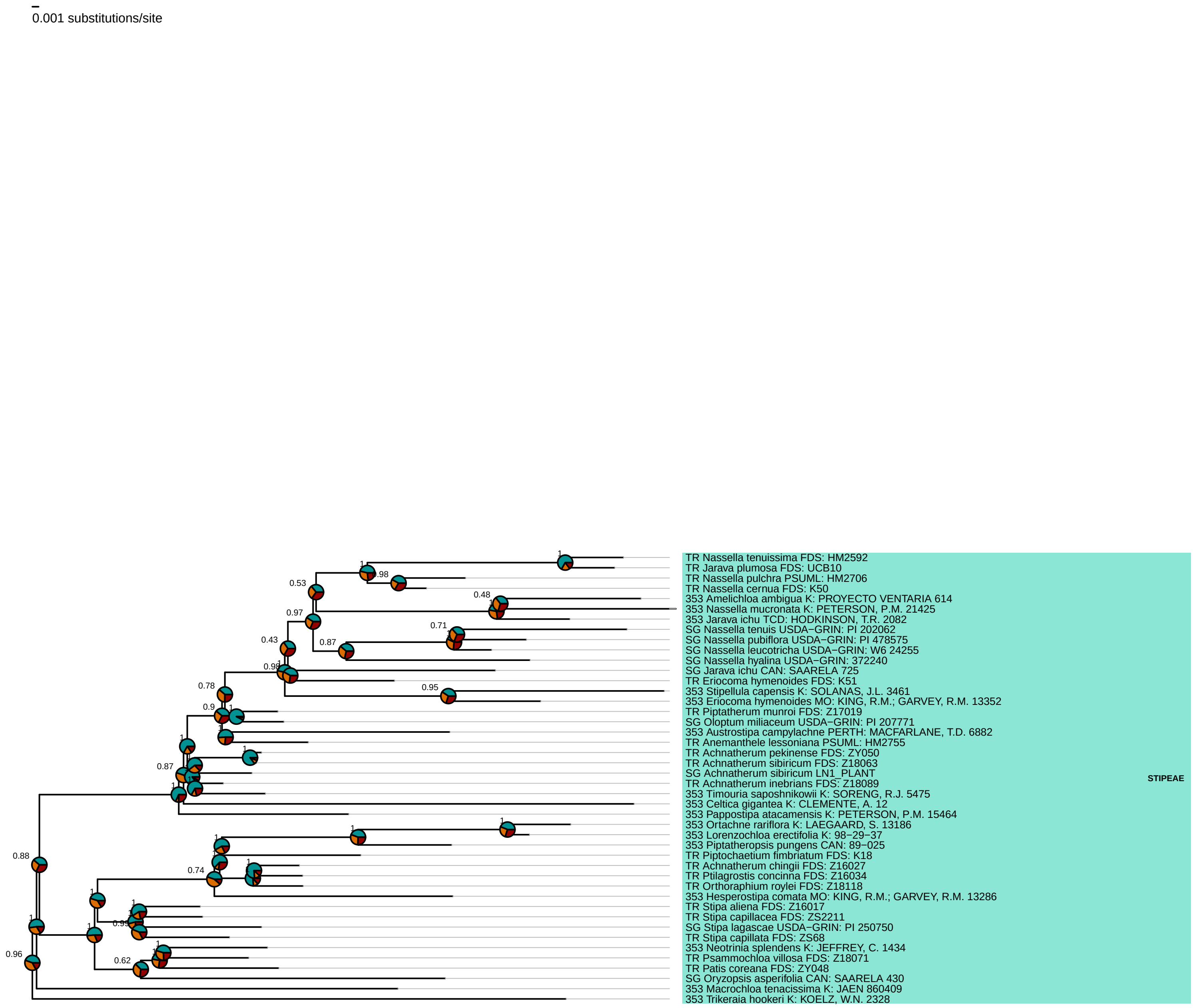

**Bambusoideae**
*(nuclear)*

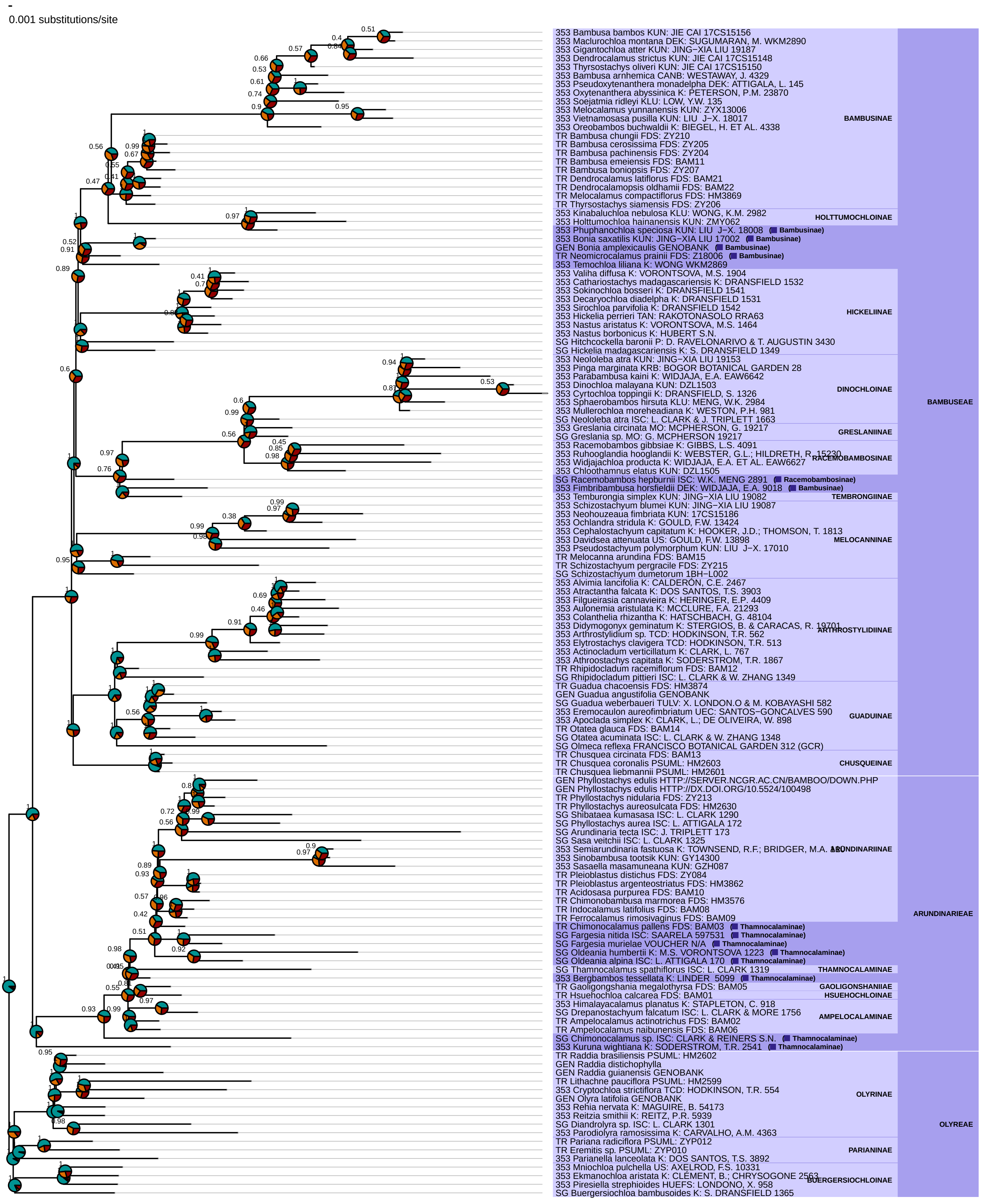

**Oryzoideae**
*(nuclear)*

0.001 substitutions/site

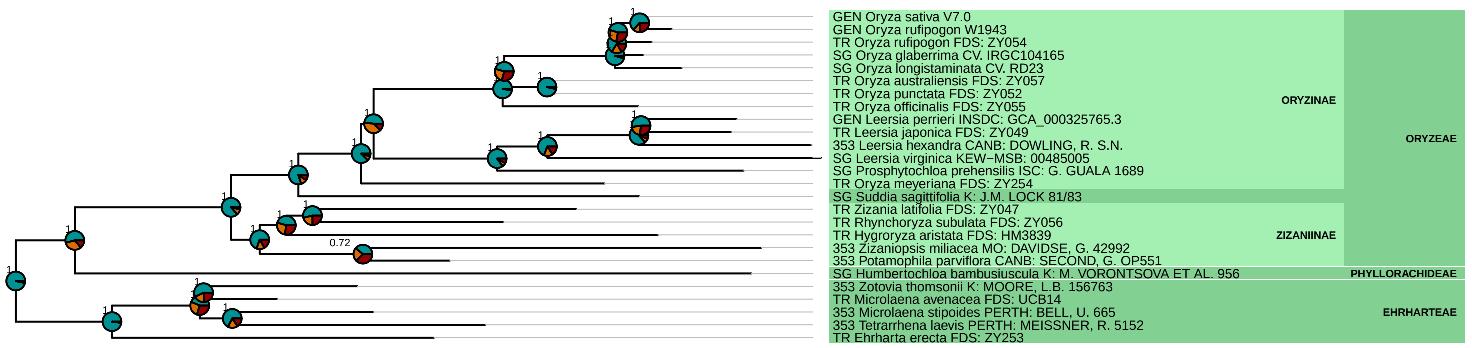

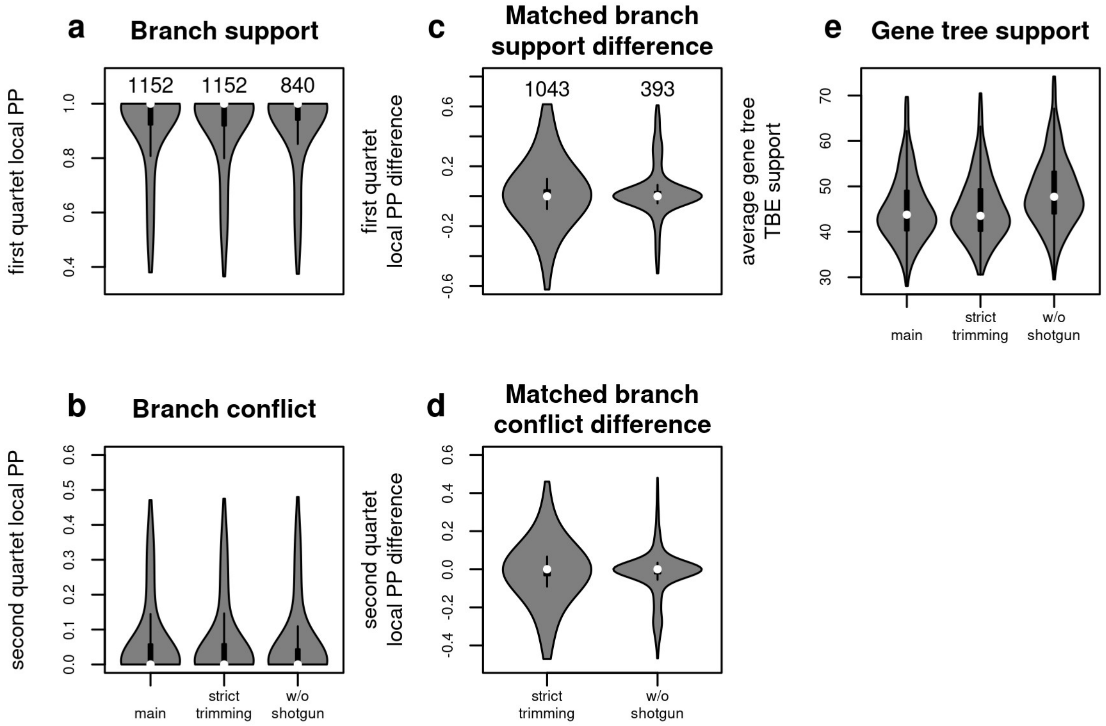


**Figure S6.** Nuclear species tree stability under different data filtering strategies. The main tree was inferred from gene alignments where columns with > 90% missing data were removed (331 genes > 500 bp retained), and included only accessions with at least 50% of gene recovery (i.e. > 166 genes). This tree was compared to trees obtained from an alignment set with more stringent trimming threshold (removal of columns > 50% missing data, 315 genes > 500 bp retained; 'stringent trimming'), and a set that excluded shotgun accessions from the main tree dataset ('w/o shotgun'). (a) Internal branch support (local posterior support for the preferred quartet configuration) across the three species trees. Numbers above violin plots give the number of internal branches per tree. (b) Internal branch conflict (local posterior support for the second-most supported quartet configuration) across the three species trees. (c) Difference in support for branches recovered with the two reduced alignment sets compared to the main tree. Numbers above violin plots give the number of matching branches. (d) Difference in conflict for branches recovered with the two reduced alignment sets compared to the main tree. (e) Mean gene tree support (transfer bootstrap expectation) across the three species trees.

**Figure S7 (following pages).** Detailed plots of the reticulations inferred with gene tree–species tree reconciliation (Fig. 2 in the main text). The black phylogeny represents the species tree. Blue lines correspond to inferred reticulate connections (transfers). Very frequent transfers (upper 10% quantile of the number of genes involved, see inset histogram) are coloured in darker blue and less frequent transfers in lighter blue. Arrowheads indicate where transfer counts are skewed by more than 50% in one direction. (a) Full Poaceae tree at tribe level, where accessions were mapped to their respective tribes. (b) Andropogoneae tribe (maize, sorghum, sugarcane and relatives). (c) Bambusoideae subfamily (bamboos). (d) Triticeae tribe (wheat, barley and relatives).

### Figure S7 − reticulations inferred with gene tree−species tree reconciliation

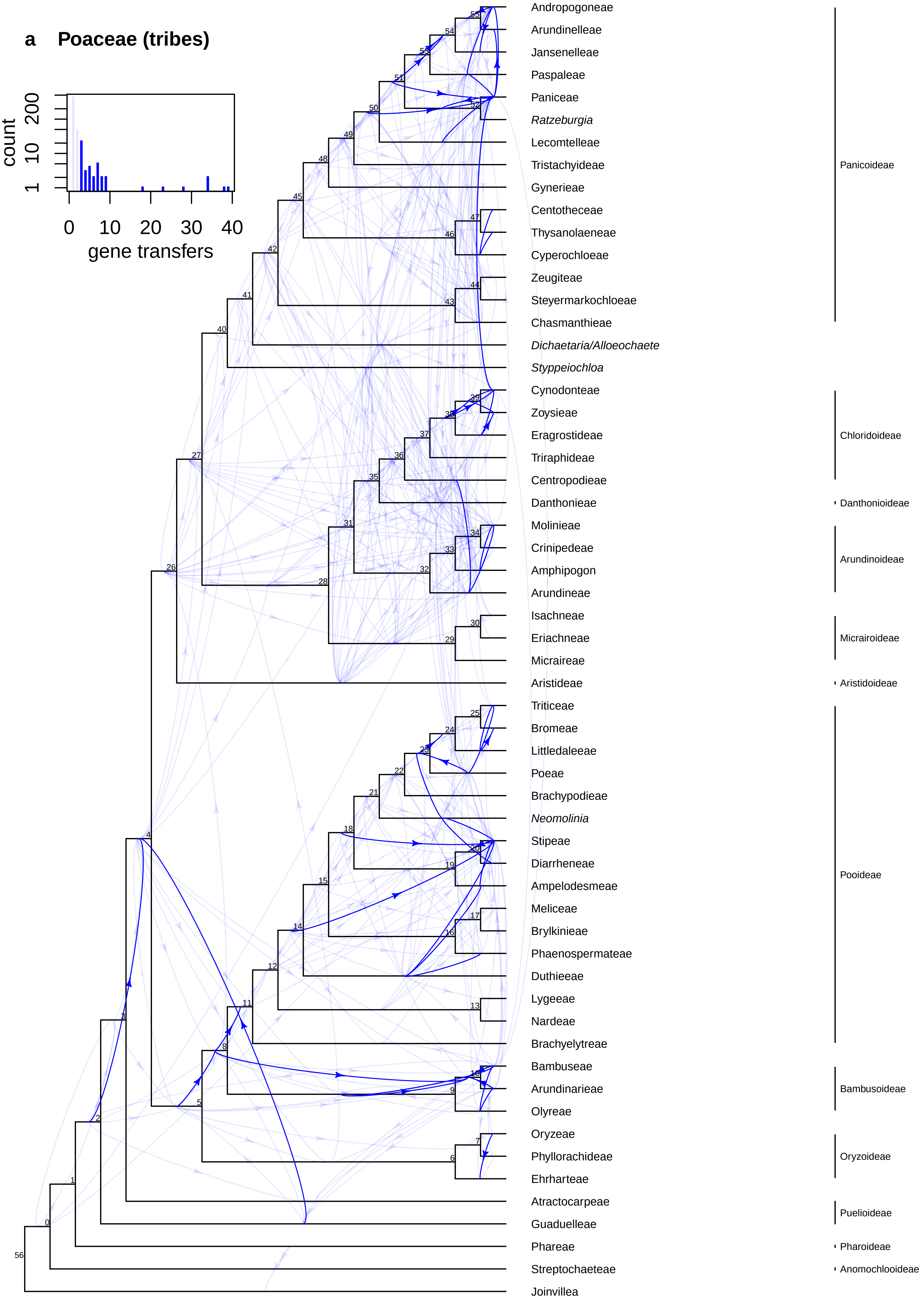

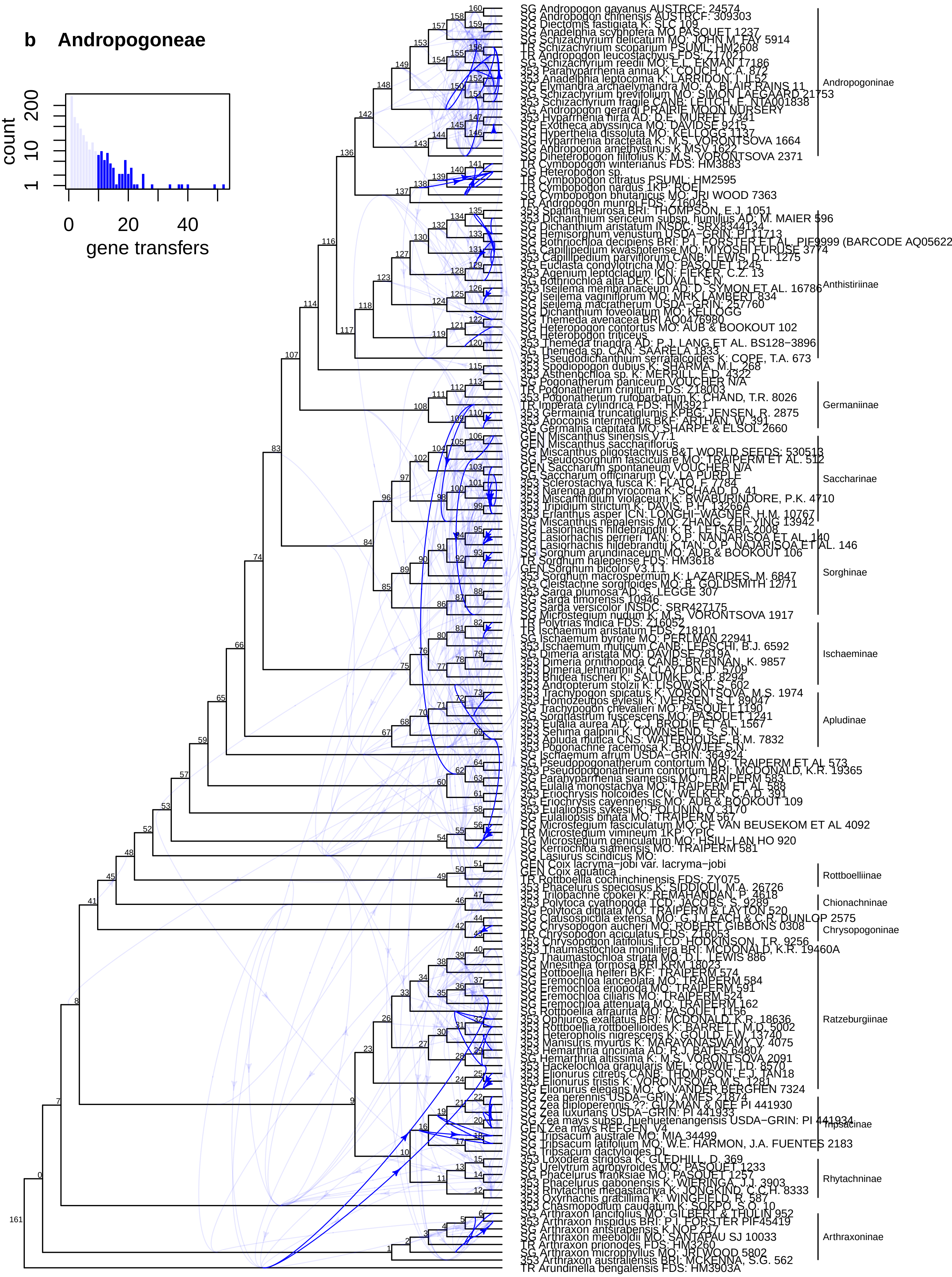
b Andropogoneae
count
1
10
200
0
20
40
gene transfers
SG Andropogon gayanus AUSTRCF: 24574
SG Andropogon chinensis AUSTRCF: 309303
SG Diectomis fastigiata K: SLC 109
SG Anadelphia scyphofera MO: PASQUET 1237
SG Schizachyrium delicatum MO: JOHN M. FAY 5914
TR Schizachyrium scoparium PSUML: HM2608
TR Andropogon leucostachyus FDS: Z17021
SG Schizachyrium reedii MO: E.L. EKMAN 17186
353 Parahyparrhenia annua K: COUCH, C.A. 872
353 Anadelphia leptocoma K: LARRIDON, I. IL52
SG Elymandra archaelymandra MO: A. BLAIR RAINS 11
SG Schizachyrium brevifolium MO: SIMON LAEGAARD 21753
353 Schizachyrium fragile CANB: LEITCH, E. NTA001838
SG Andropogon gerardi PRAIRIE MOON NURSERY
353 Hyparrhenia hirta AD: D.E. MURFET 7341
SG Exotheca abyssinica MO: DAVIDSE 9215
SG Hyperthelia dissoluta MO: KELLOGG 1137
SG Hyparrhenia bracteata K: M.S. VORONTSOVA 1664
SG Andropogon amethystinus K: MSV 1622
SG Diheteropogon filifolius K: M.S. VORONTSOVA 2371
TR Cymbopogon winterianus FDS: HM3883
SG Heteropogon sp.
TR Cymbopogon citratus PSUML: HM2595
TR Cymbopogon nardus 1KP: ROEL
SG Cymbopogon bhutanicus MO: JRI WOOD 7363
TR Andropogon munroi FDS: Z16045
353 Spathia neurosa BRI: THOMPSON, E.J. 1051
353 Dichanthium sericeum subsp. humilius AD: M. MAIER 596
SG Dichanthium aristatum INSDC: SRX8344134
SG Hemisorghum venustum USDA−GRIN: PI 11713
SG Bothriochloa decipiens BRI: P.I. FORSTER ET AL. PIF9999 (BARCODE AQ05622
SG Capillipedium kwashotense MO: MIYOSHI FURUSE 3774
353 Capillipedium parviflorum CANB: LEWIS, D.L. 1275
SG Euclasta condylotricha MO: PASQUET 1245
353 Agenium leptocladum ICN: IEKER, C.Z. 13
SG Bothriochloa alta DEK: DUVALL S.N.
353 Iseilema membranaceum AD: D. SYMON ET AL. 16786
SG Iseilema vaginiflorum MO: MRK LAMBERT 834
SG Iseilema macratherum USDA−GRIN: 257760
SG Dichanthium foveolatum MO: KELLOGG
SG Themeda avenacea BRI AQ0476980
SG Heteropogon contortus MO: AUB & BOOKOUT 102
SG Heteropogon triticeus
353 Themeda triandra AD: P.J. LANG ET AL. BS128−3896
SG Themeda sp. CAN: SAARELA 1833
353 Pseudodichanthium serrafalcoides K: COPE, T.A. 673
353 Spodiopogon dubius K: SHARMA, M. 268
353 Asthenochloa sp. K: MERRILL, E.D. 4322
SG Pogonatherum paniceum VOUCHER N/A
TR Pogonatherum crinitum FDS: Z18003
353 Pogonatherum rufobarbatum K: CHAND, T.R. 8026
TR Imperata cylindrica FDS: HM3921
353 Germainia truncatiglumis KPBG: JENSEN, R. 2875
353 Apocopis intermedius BKF: ARTHAN, W. 391
SG Germainia capitata MO: SHARPE & ELSOL 2660
GEN Miscanthus sinensis V7.1
GEN Miscanthus sacchariflorus
SG Miscanthus oligostachyus B&T WORLD SEEDS: 530518
SG Pseudosorghum fasciculare MO: TRAIPERM ET AL. 512
GEN Saccharum spontaneum VOUCHER N/A
SG Saccharum officinarum CV. LA PURPLE
353 Sclerostachya fusca K: FLATO, F 7784
353 Narenga porphyrocoma K: SCHAAD, D. 41
353 Miscanthidium violaceum K: BWABURINDORE, P.K. 4710
353 Tripidium strictum K: DAVIS, P.H. 13266A
353 Erianthus asper ICN: LONGHI−WAGNER, H.M. 10767
SG Miscanthus nepalensis MO: ZHANG, ZHI−YING 13942
SG Lasiorhachis hildebrandtii K: R. LETSARA 2008
SG Lasiorhachis perrieri TAN: O.P. NANJARISOA ET AL. 140
SG Lasiorhachis hildebrandtii K,TAN: O.P. NAJARISOA ET AL. 146
SG Sorghum arundinaceum MO: AUB & BOOKOUT 106
TR Sorghum halepense FDS: HM3618
GEN Sorghum bicolor V3.1.1
353 Sorghum macrospermum K: LAZARIDES, M. 6847
SG Cleistachne sorghoides MO: B. GOLDSMITH 12/71
353 Sarga plumosa AD: S. LEGGE 307
SG Sarga timorensis 10946
SG Sarga versicolor INSDC: SRR427175
SG Microstegium nudum K: M.S. VORONTSOVA 1917
TR Polytrias indica FDS: Z16052
TR Ischaemum aristatum FDS: Z18101
SG Ischaemum byrone MO: PERLMAN 22941
353 Ischaemum muticum CANB: LEPSCHI, B.J. 6592
SG Dimeria aristata MO: DAVIDSE 7819A
353 Dimeria ornithopoda CANB: BRENNAN, K. 9857
353 Dimeria lehmannii K: CLAYTON, D. 5709
353 Bhidea fischeri K: SALUNKE, C.B. 8294
353 Andropterum stolzii K: LISOWSKI, S. 602
353 Trachypogon spicatus K: VORONTSOVA, M.S. 1974
353 Homozeugos eylesii K: IVERSEN, S.T. 89047
SG Trachypogon chevalieri MO: PASQUET 1190
SG Sorghastrum fuscescens MO: PASQUET 1241
353 Eulalia aurea AD: C.J. BRODIE ET AL. 1567
353 Sehima galpinii K: TOWNSEND, S. S.N.
353 Apluda mutica CNS: WATERHOUSE, B.M. 7832
353 Pogonachne racemosa K: BOWDEN S.N.
SG Ischaemum afrum USDA−GRIN: 364924
SG Pseudopogonatherum contortum MO: TRAIPERM ET AL 573
353 Pseudopogonatherum contortum BRI: MCDONALD, K.R. 19365
SG Parahyparrhenia siamensis MO: TRAIPERM 583
SG Eulalia monostachya MO: TRAIPERM ET AL 588
353 Eriochrysis holcoides ICN: WELKER, C.A.D. 391
SG Eriochrysis cayennensis MO: AUB & BOOKOUT 109
353 Eulaliopsis sykesii K: POLUNIN, O. 3170
SG Eulaliopsis binata MO: TRAIPERM 567
SG Microstegium fasciculatum MO: CF VAN BEUSEKOM ET AL 4092
TR Microstegium vimineum 1KP: YPIC
SG Microstegium geniculatum MO: HSIU−LAN HO 920
SG Kerriochloa siamensis MO: TRAIPERM 581
SG Lasiurus scindicus MO:
GEN Coix lacryma−jobi var. lacryma−jobi
GEN Coix aquatica
TR Rottboellia cochinchinensis FDS: ZY075
353 Phacelurus speciosus K: SIDDIQUI, M.A. 26726
353 Trilobachne cookei K: REMAHANDAN, P. 4618
353 Polytoca cyathopoda TCD: JACOBS, S. 9289
SG Polytoca digitata MO: TRAIPERM & LAYTON 520
SG Clausospicula extensa MO: G.J. LEACH & C.R. DUNLOP 2575
SG Chrysopogon aucheri MO: ROBERT GIBBONS 0308
TR Chrysopogon aciculatus FDS: Z16053
353 Chrysopogon latifolius TCD: HODKINSON, T.R. 9256
353 Thaumastochloa monilifera BRI: MCDONALD, K.R. 19460A
SG Thaumastochloa striata MO: D.L. LEWIS 886
SG Mnesithea formosa BRI KRM 18023
SG Rottboellia helferi BKF: TRAIPERM 574
SG Eremochloa lanceolata MO: TRAIPERM 584
SG Eremochloa eriopoda MO: TRAIPERM 591
SG Eremochloa ciliaris MO: TRAIPERM 524
SG Eremochloa attenuata MO: TRAIPERM 162
SG Rottboellia afraurita MO: PASQUET 1156
353 Ophiuros exaltatus BRI: MCDONALD, K.R. 18636
353 Rottboellia rottboellioides K: BARRETT, M.D. 5002
353 Heteropholis nigrescens K: GOULD, F.W. 13740
353 Manisuris myurus K: MARAYANASWAMY, V. 4075
353 Hemarthria uncinata AD: R.J. BATES 64807
SG Hemarthria altissima K: M.S. VORONTSOVA 2091
353 Hackelochloa granularis MEL: COWIE, I.D. 8570
353 Elionurus citreus CANB: THOMPSON, E.J. TAN18
353 Elionurus tristis K: VORONTSOVA, M.S. 1281
SG Elionurus elegans MO: C. VANDER BERGHEN 7324
SG Zea perennis USDA−GRIN: AMES 21874
SG Zea diploperennis ??: GUZMAN & NEE PI 441930
SG Zea luxurians USDA−GRIN: PI 441933
SG Zea mays subsp. huehuetenangensis USDA−GRIN: PI 441934
GEN Zea mays REFGEN_V4
SG Tripsacum australe MO: MIA 34499
SG Tripsacum latifolium MO: W.E. HARMON, J.A. FUENTES 2183
SG Tripsacum dactyloides DL
353 Loxodera strigosa K: GLEDHILL, D. 369
SG Urelytrum agropyroides MO: PASQUET 1233
SG Phacelurus franksiae MO: PASQUET 1257
353 Phacelurus gabonensis K: WIERINGA, J.J. 3903
353 Rhytachne megastachya K: JONGKIND, C.C.H. 8333
353 Oxyrhachis gracillima K: WINGFIELD, R. 587
353 Chasmopodium caudatum K: SOKPO, S.O. 10
SG Arthraxon lancifolius MO: GILBERT & THULIN 952
353 Arthraxon hispidus BRI: P.I. FORSTER PIF45419
SG Arthraxon antsirabensis K: KNOP 27
SG Arthraxon meeboldii MO: SANTAPAU SJ 10033
TR Arthraxon prionodes FDS: HM3260
SG Arthraxon microphyllus MO: JRI WOOD 5802
353 Arthraxon australiensis BRI: MCKENNA, S.G. 562
TR Arundinella bengalensis FDS: HM3903A
Andropogoninae
Anthistiriinae
Germaniinae
Saccharinae
Sorghinae
Ischaeminae
Apludinae
Rottboelliinae
Chionachninae
Chrysopogoninae
Ratzeburgiinae
Tripsacinae
Rhytachninae
Arthraxoninae

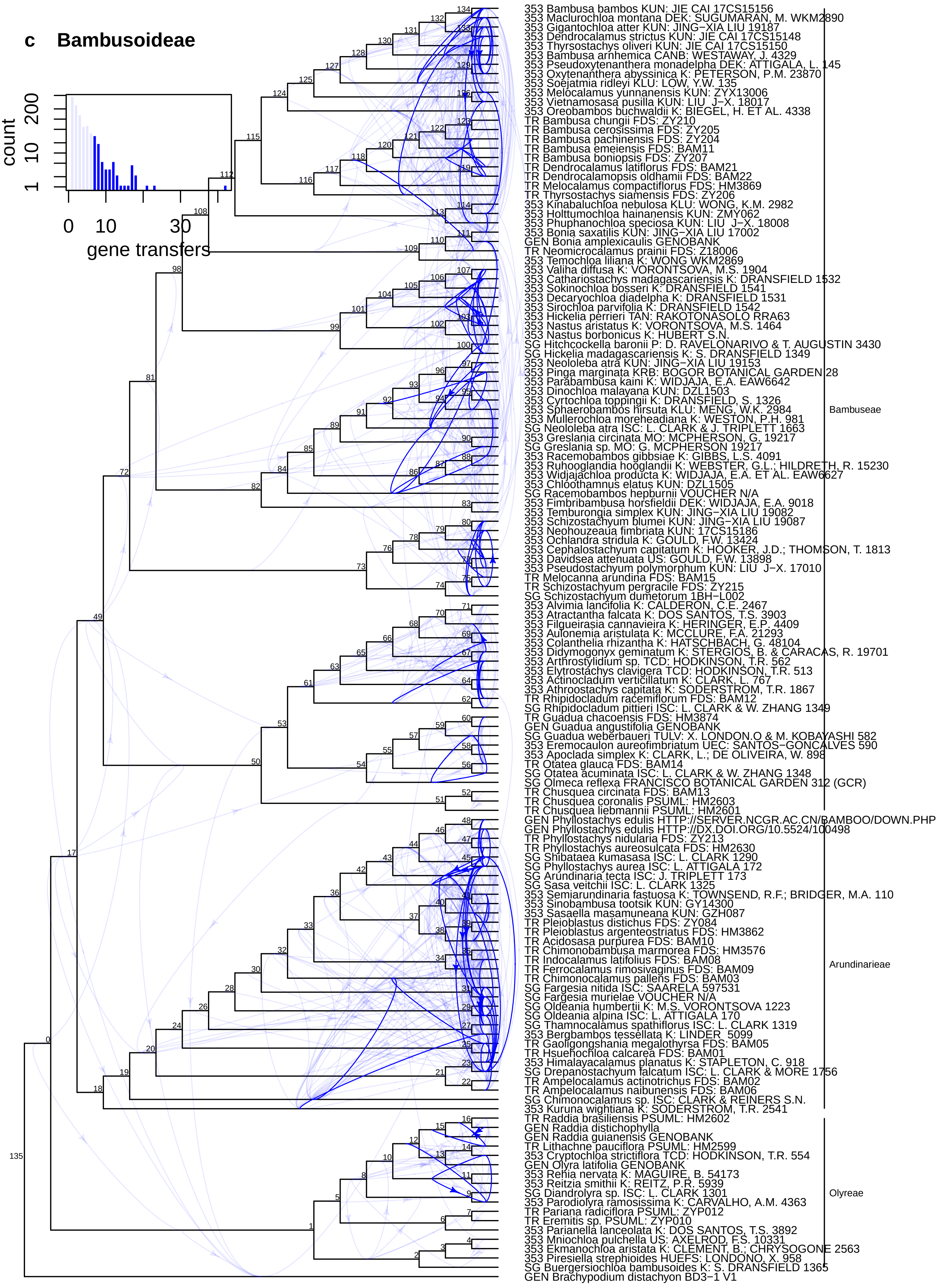
c Bambusoideae
count
1
10
200
0
10
30
gene transfers
353 Bambusa bambos KUN: JIE CAI 17CS15156
353 Maclurochloa montana DEK: SUGUMARAN, M. WKM2890
353 Gigantochloa atter KUN: JING−XIA LIU 19187
353 Dendrocalamus strictus KUN: JIE CAI 17CS15148
353 Thyrsostachys oliveri KUN: JIE CAI 17CS15150
353 Bambusa arnhemica CANB: WESTAWAY, J. 4329
353 Pseudoxytenanthera monadelpha DEK: ATTIGALA, L. 145
353 Oxytenanthera abyssinica K: PETERSON, P.M. 23870
353 Soejatmia ridleyi KLU: LOW, Y.W. 135
353 Melocalamus yunnanensis KUN: ZYX13006
353 Vietnamosasa pusilla KUN: LIU J−X. 18017
353 Oreobambos buchwaldii K: BIEGEL, H. ET AL. 4338
TR Bambusa chungii FDS: ZY210
TR Bambusa cerosissima FDS: ZY205
TR Bambusa pachinensis FDS: ZY204
TR Bambusa emeiensis FDS: BAM11
TR Bambusa boniopsis FDS: ZY207
TR Dendrocalamus latiflorus FDS: BAM21
TR Dendrocalamopsis oldhamii FDS: BAM22
TR Melocalamus compactiflorus FDS: HM3869
TR Thyrsostachys siamensis FDS: ZY206
353 Kinabaluchloa nebulosa KLU: WONG, K.M. 2982
353 Holttumochloa hainanensis KUN: ZMY062
353 Phuphanochloa speciosa KUN: LIU J−X. 18008
353 Bonia saxatilis KUN: JING−XIA LIU 17002
GEN Bonia amplexicaulis GENOBANK
TR Neomicrocalamus prainii FDS: Z18006
353 Temochloa liliana K: WONG WKM2869
353 Valiha diffusa K: VORONTSOVA, M.S. 1904
353 Cathariostachys madagascariensis K: DRANSFIELD 1532
353 Sokinochloa bosseri K: DRANSFIELD 1541
353 Decaryochloa diadelpha K: DRANSFIELD 1531
353 Sirochloa parvifolia K: DRANSFIELD 1542
353 Hickelia perrieri TAN: RAKOTONASOLO RRA63
353 Nastus aristatus K: VORONTSOVA, M.S. 1464
353 Nastus borbonicus K: HUBERT S.N.
SG Hitchcockella baronii P: D. RAVELONARIVO & T. AUGUSTIN 3430
SG Hickelia madagascariensis K: S. DRANSFIELD 1349
353 Neololeba atra KUN: JING−XIA LIU 19153
353 Pinga marginata KRB: BOGOR BOTANICAL GARDEN 28
353 Parabambusa kaini K: WIDJAJA, E.A. EAW6642
353 Dinochloa malayana KUN: DZL1503
353 Cyrtochloa toppingii K: DRANSFIELD, S. 1326
353 Sphaerobambos hirsuta KLU: MENG, W.K. 2984
353 Mullerochloa moreheadiana K: WESTON, P.H. 981
SG Neololeba atra ISC: L. CLARK & J. TRIPLETT 1663
353 Greslania circinata MO: MCPHERSON, G. 19217
SG Greslania sp. MO: G. MCPHERSON 19217
353 Racemobambos gibbsiae K: GIBBS, L.S. 4091
353 Ruhooglandia hooglandii K: WEBSTER, G.L.; HILDRETH, R. 15230
353 Widjajachloa producta K: WIDJAJA, E.A. ET AL. EAW6627
353 Chloothamnus elatus KUN: DZL1505
SG Racemobambos hepburnii VOUCHER N/A
353 Fimbribambusa horsfieldii DEK: WIDJAJA, E.A. 9018
353 Temburongia simplex KUN: JING−XIA LIU 19082
353 Schizostachyum blumei KUN: JING−XIA LIU 19087
353 Neohouzeaua fimbriata KUN: 17CS15186
353 Ochlandra stridula K: GOULD, F.W. 13424
353 Cephalostachyum capitatum K: HOOKER, J.D.; THOMSON, T. 1813
353 Davidsea attenuata US: GOULD, F.W. 13898
353 Pseudostachyum polymorphum KUN: LIU J−X. 17010
TR Melocanna arundina FDS: BAM15
TR Schizostachyum pergracile FDS: ZY215
SG Schizostachyum dumetorum 1BH−L002
353 Alvimia lancifolia K: CALDERON, C.E. 2467
353 Atractantha falcata K: DOS SANTOS, T.S. 3903
353 Filgueirasia cannavieira K: HERINGER, E.P. 4409
353 Aulonemia aristulata K: MCCLURE, F.A. 21293
353 Colanthelia rhizantha K: HATSCHBACH, G. 48104
353 Didymogonyx geminatum K: STERGIOS, B. & CARACAS, R. 19701
353 Arthrostylidium sp. TCD: HODKINSON, T.R. 562
353 Elytrostachys clavigera TCD: HODKINSON, T.R. 513
353 Actinocladum verticillatum K: CLARK, L. 767
353 Athroostachys capitata K: SODERSTROM, T.R. 1867
TR Rhipidocladum racemiflorum FDS: BAM12
SG Rhipidocladum pittieri ISC: L. CLARK & W. ZHANG 1349
TR Guadua chacoensis FDS: HM3874
GEN Guadua angustifolia GENOBANK
SG Guadua weberbaueri TULV: X. LONDON.O & M. KOBAYASHI 582
353 Eremocaulon aureofimbriatum UEC: SANTOS−GONCALVES 590
353 Apoclada simplex K: CLARK, L.; DE OLIVEIRA, W. 898
TR Otatea glauca FDS: BAM14
SG Otatea acuminata ISC: L. CLARK & W. ZHANG 1348
SG Olmeca reflexa FRANCISCO BOTANICAL GARDEN 312 (GCR)
TR Chusquea circinata FDS: BAM13
TR Chusquea coronalis PSUML: HM2603
TR Chusquea liebmannii PSUML: HM2601
GEN Phyllostachys edulis HTTP://SERVER.NCGR.AC.CN/BAMBOO/DOWN.PHP
GEN Phyllostachys edulis HTTP://DX.DOI.ORG/10.5524/100498
TR Phyllostachys nidularia FDS: ZY213
TR Phyllostachys aureosulcata FDS: HM2630
SG Shibataea kumasasa ISC: L. CLARK 1290
SG Phyllostachys aurea ISC: L. ATTIGALA 172
SG Arundinaria tecta ISC: J. TRIPLETT 173
SG Sasa veitchii ISC: L. CLARK 1325
353 Semiarundinaria fastuosa K: TOWNSEND, R.F.; BRIDGER, M.A. 110
353 Sinobambusa tootsik KUN: GY14300
353 Sasaella masamuneana KUN: GZH087
TR Pleioblastus distichus FDS: ZY084
TR Pleioblastus argenteostriatus FDS: HM3862
TR Acidosasa purpurea FDS: BAM10
TR Chimonobambusa marmorea FDS: HM3576
TR Indocalamus latifolius FDS: BAM08
TR Ferrocalamus rimosivaginus FDS: BAM09
TR Chimonocalamus pallens FDS: BAM03
SG Fargesia nitida ISC: SAARELA 597531
SG Fargesia murielae VOUCHER N/A
SG Oldeania humbertii K: M.S. VORONTSOVA 1223
SG Oldeania alpina ISC: L. ATTIGALA 170
SG Thamnocalamus spathiflorus ISC: L. CLARK 1319
353 Bergbambos tessellata K: LINDER 5099
TR Gaoligongshania megalothyrsa FDS: BAM05
TR Hsuehochloa calcarea FDS: BAM01
353 Himalayacalamus planatus K: STAPLETON, C. 918
SG Drepanostachyum falcatum ISC: L. CLARK & MORE 1756
TR Ampelocalamus actinotrichus FDS: BAM02
TR Ampelocalamus naibunensis FDS: BAM06
SG Chimonocalamus sp. ISC: CLARK & REINERS S.N.
353 Kuruna wightiana K: SODERSTROM, T.R. 2541
TR Raddia brasiliensis PSUML: HM2602
GEN Raddia distichophylla
GEN Raddia guianensis GENOBANK
TR Lithachne pauciflora PSUML: HM2599
353 Cryptochloa strictiflora TCD: HODKINSON, T.R. 554
GEN Olyra latifolia GENOBANK
353 Rehia nervata K: MAGUIRE, B. 54173
353 Reitzia smithii K: REITZ, P.R. 5939
SG Diandrolyra sp. ISC: L. CLARK 1301
353 Parodiolyra ramosissima K: CARVALHO, A.M. 4363
TR Pariana radicifiora PSUML: ZYP012
TR Eremitis sp. PSUML: ZYP010
353 Parianella lanceolata K: DOS SANTOS, T.S. 3892
353 Mniochloa pulchella US: AXELROD, F.S. 10331
353 Ekmanochloa aristata K: CLEMENT, B.; CHRYSOGONE 2563
353 Piresiella strephioides HUEFS: LONDONO, X. 958
SG Buergersiochloa bambusoides K: S. DRANSFIELD 1365
GEN Brachypodium distachyon BD3−1 V1
Bambuseae
Arundinarieae
Olyreae

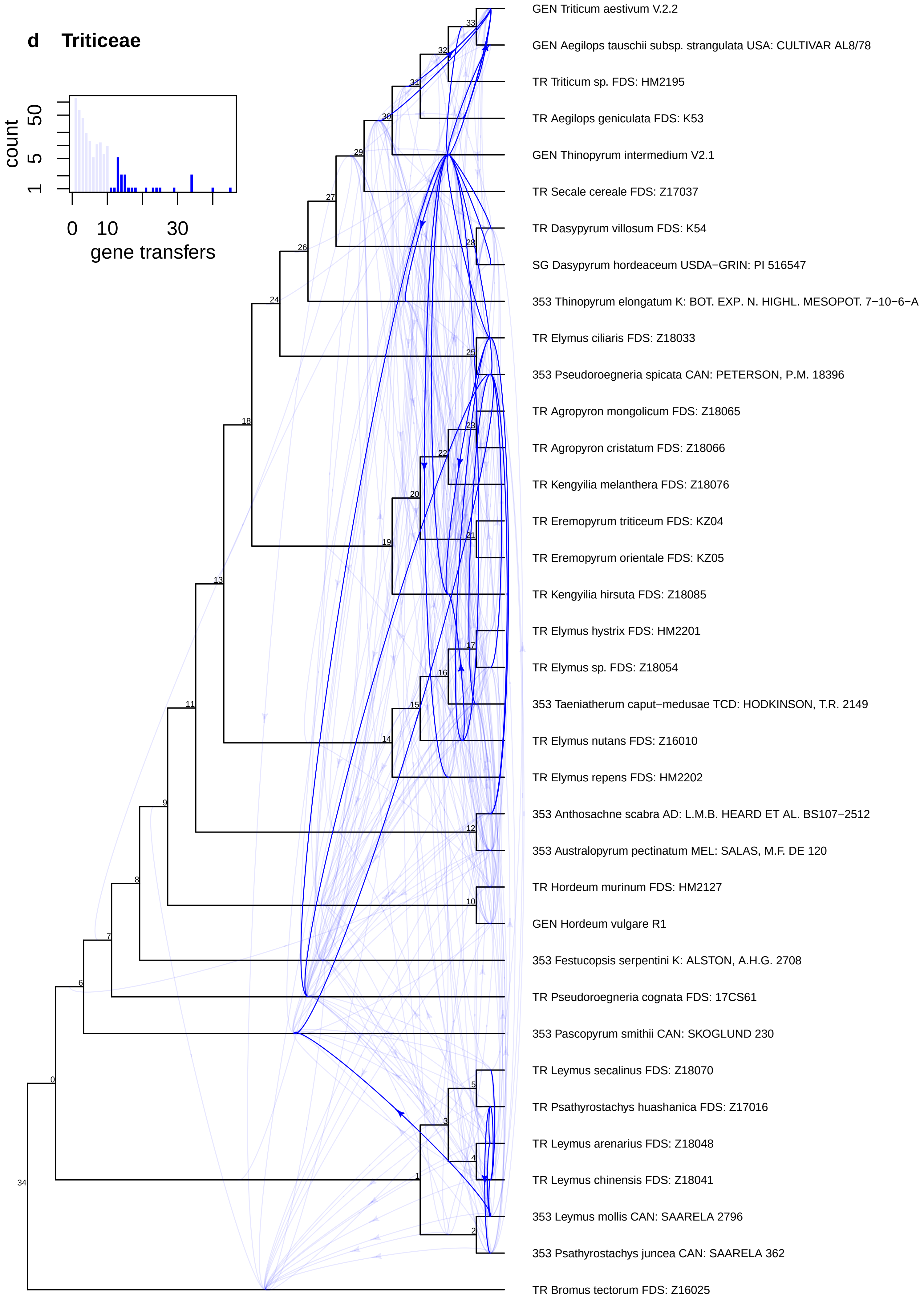
d Triticeae
count
1
5
50
0
10
30
gene transfers
GEN Triticum aestivum V.2.2
GEN Aegilops tauschii subsp. strangulata USA: CULTIVAR AL8/78
TR Triticum sp. FDS: HM2195
TR Aegilops geniculata FDS: K53
GEN Thinopyrum intermedium V2.1
TR Secale cereale FDS: Z17037
TR Dasypyrum villosum FDS: K54
SG Dasypyrum hordeaceum USDA−GRIN: PI 516547
353 Thinopyrum elongatum K: BOT. EXP. N. HIGHL. MESOPOT. 7−10−6−A
TR Elymus ciliaris FDS: Z18033
353 Pseudoroegneria spicata CAN: PETERSON, P.M. 18396
TR Agropyron mongolicum FDS: Z18065
TR Agropyron cristatum FDS: Z18066
TR Kengyilia melanthera FDS: Z18076
TR Eremopyrum triticeum FDS: KZ04
TR Eremopyrum orientale FDS: KZ05
TR Kengyilia hirsuta FDS: Z18085
TR Elymus hystrix FDS: HM2201
TR Elymus sp. FDS: Z18054
353 Taeniatherum caput−medusae TCD: HODKINSON, T.R. 2149
TR Elymus nutans FDS: Z16010
TR Elymus repens FDS: HM2202
353 Anthosachne scabra AD: L.M.B. HEARD ET AL. BS107−2512
353 Australopyrum pectinatum MEL: SALAS, M.F. DE 120
TR Hordeum murinum FDS: HM2127
GEN Hordeum vulgare R1
353 Festucopsis serpentini K: ALSTON, A.H.G. 2708
TR Pseudoroegneria cognata FDS: 17CS61
353 Pascopyrum smithii CAN: SKOGLUND 230
TR Leymus secalinus FDS: Z18070
TR Psathyrostachys huashanica FDS: Z17016
TR Leymus arenarius FDS: Z18048
TR Leymus chinensis FDS: Z18041
353 Leymus mollis CAN: SAARELA 2796
353 Psathyrostachys juncea CAN: SAARELA 362
TR Bromus tectorum FDS: Z16025

**Figure S8 (following pages).** Detailed version of the plastome tree inferred from a concatenated alignment of 70 coding regions and *trnL*-*trnF* (tree on the right in Fig. 3 in the main text) for 910 accessions, broken down into subclades. Text labels at nodes give the branch support as transfer bootstrap expectation. Tip labels show data type, species and voucher, isolate or germplasm information, where available, for each accession. Taxa from subtribe to subfamily level are labelled with coloured polygons. Taxonomic outliers falling outside the clade corresponding to their nominal taxon are labelled in brackets after the accession information.

### Figure S8 − Plastome tree (RAxML GTR+CAT, CDS + trnL−trnF)

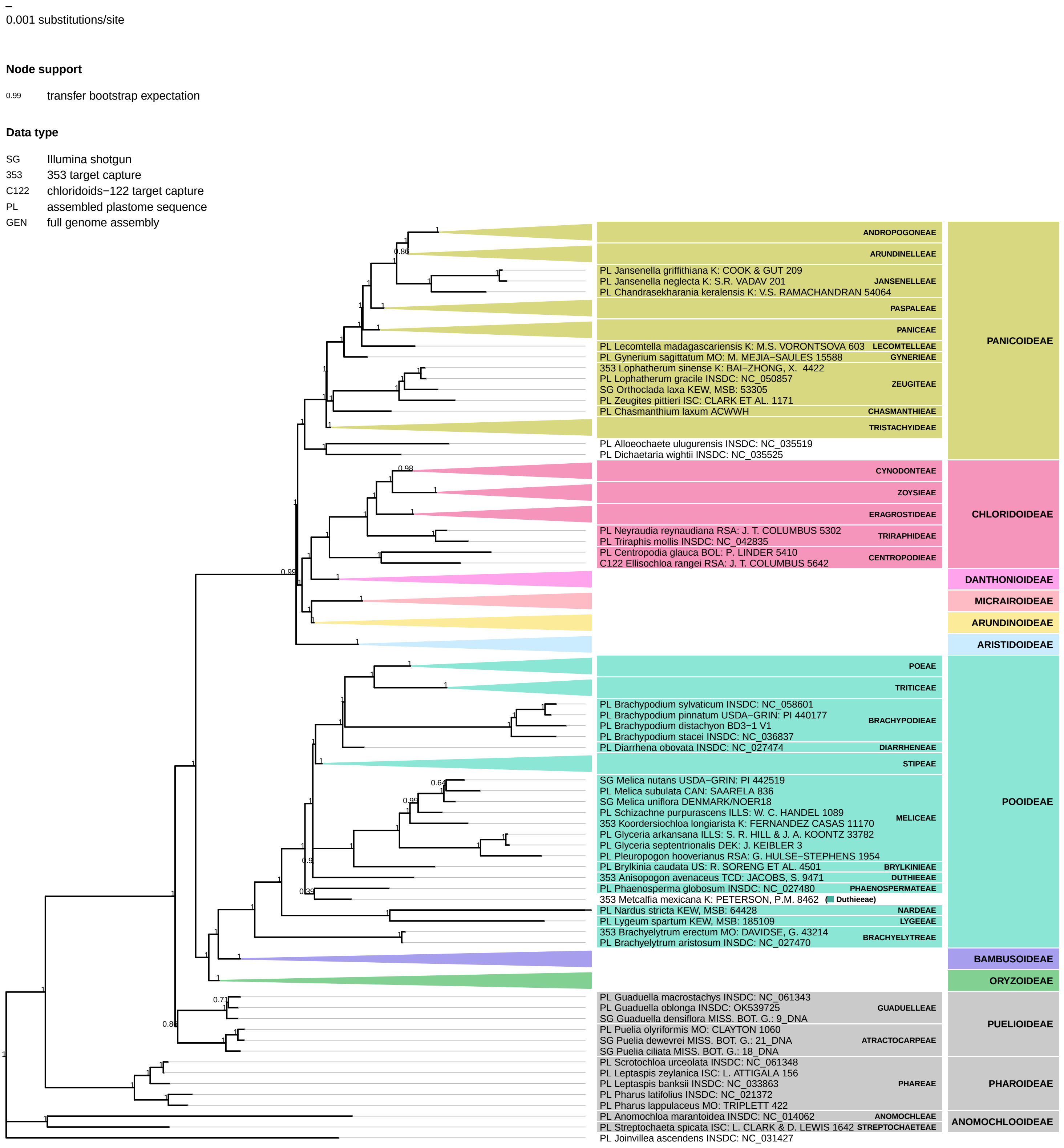

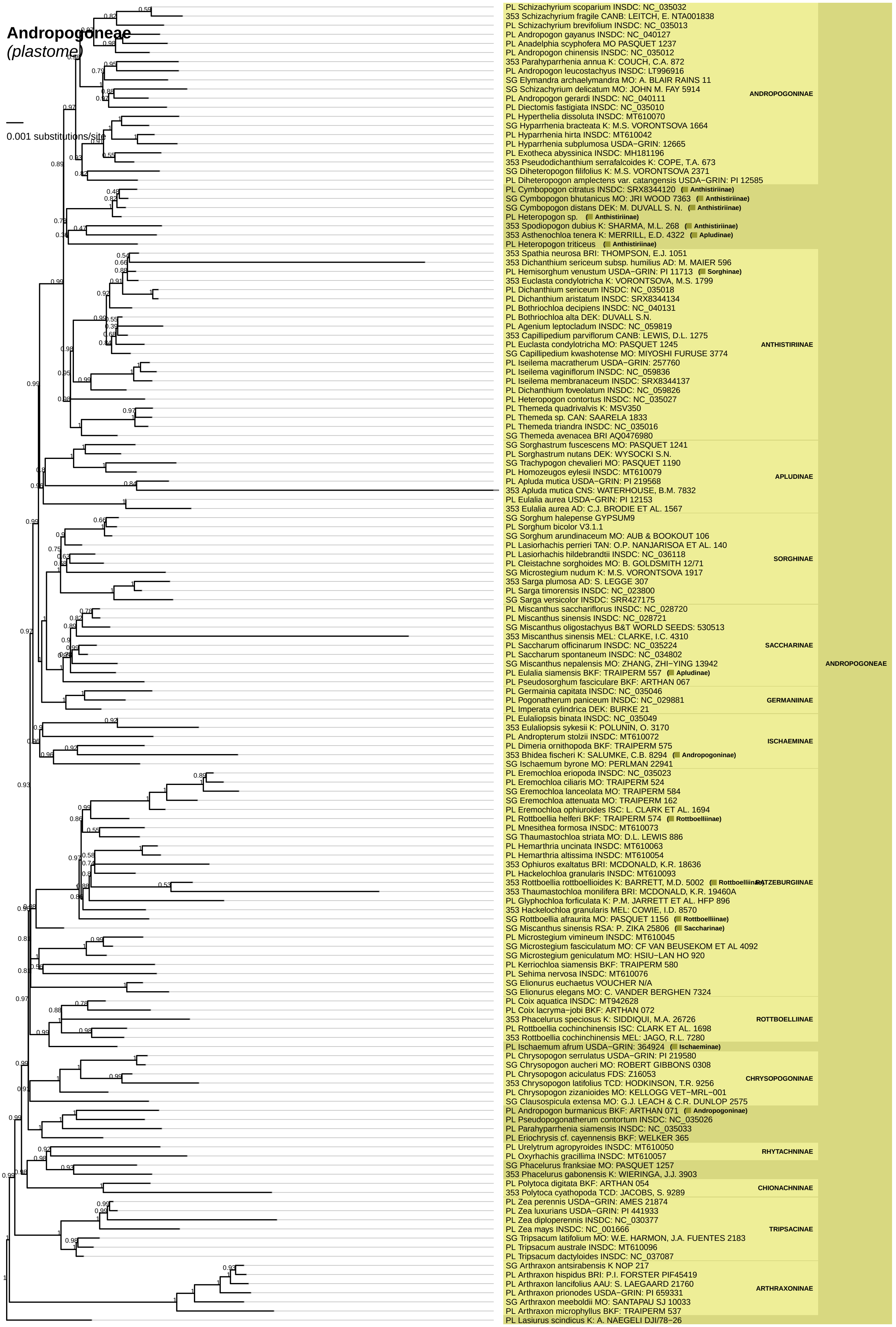
Andropogoneae
(plastome)
0.001 substitutions/site
PL Schizachyrium scoparium INSDC: NC_035032
353 Schizachyrium fragile CANB: LEITCH, E. NTA001838
PL Schizachyrium brevifolium INSDC: NC_035013
PL Andropogon gayanus INSDC: NC_040127
PL Anadelphia scyphofera MO PASQUET 1237
PL Andropogon chinensis INSDC: NC_035012
353 Parahyparrhenia annua K: COUCH, C.A. 872
PL Andropogon leucostachyus INSDC: LT996916
SG Elymandra archaelymandra MO: A. BLAIR RAINS 11
SG Schizachyrium delicatum MO: JOHN M. FAY 5914
PL Andropogon gerardi INSDC: NC_040111
PL Diectomis fastigiata INSDC: NC_035010
PL Hyperthelia dissoluta INSDC: MT610070
SG Hyparrhenia bracteata K: M.S. VORONTSOVA 1664
PL Hyparrhenia hirta INSDC: MT610042
PL Hyparrhenia subplumosa USDA−GRIN: 12665
PL Exotheca abyssinica INSDC: MH181196
353 Pseudodichanthium serrafalcoides K: COPE, T.A. 673
SG Diheteropogon filifolius K: M.S. VORONTSOVA 2371
PL Diheteropogon amplectens var. catangensis USDA−GRIN: PI 12585
ANDROPOGONINAE
PL Cymbopogon citratus INSDC: SRX8344120 (Anthistiriinae)
SG Cymbopogon bhutanicus MO: JRI WOOD 7363 (Anthistiriinae)
SG Cymbopogon distans DEK: M. DUVALL S. N. (Anthistiriinae)
PL Heteropogon sp. (Anthistiriinae)
353 Spodiopogon dubius K: SHARMA, M.L. 268 (Anthistiriinae)
353 Asthenochloa tenera K: MERRILL, E.D. 4322 (Apludinae)
PL Heteropogon triticeus (Anthistiriinae)
353 Spathia neurosa BRI: THOMPSON, E.J. 1051
353 Dichanthium sericeum subsp. humilius AD: M. MAIER 596
PL Hemisorghum venustum USDA−GRIN: PI 11713 (Sorghinae)
353 Euclasta condylotricha K: VORONTSOVA, M.S. 1799
PL Dichanthium sericeum INSDC: NC_035018
PL Dichanthium aristatum INSDC: SRX8344134
PL Bothriochloa decipiens INSDC: NC_040131
PL Bothriochloa alta DEK: DUVALL S.N.
PL Agenium leptocladum INSDC: NC_059819
353 Capillipedium parviflorum CANB: LEWIS, D.L. 1275
PL Euclasta condylotricha MO: PASQUET 1245
SG Capillipedium kwashotense MO: MIYOSHI FURUSE 3774
PL Iseilema macratherum USDA−GRIN: 257760
PL Iseilema vaginiflorum INSDC: NC_059836
PL Iseilema membranaceum INSDC: SRX8344137
PL Dichanthium foveolatum INSDC: NC_059826
PL Heteropogon contortus INSDC: NC_035027
PL Themeda quadrivalvis K: MSV350
PL Themeda sp. CAN: SAARELA 1833
PL Themeda triandra INSDC: NC_035016
SG Themeda avenacea BRI AQ0476980
ANTHISTIRIINAE
SG Sorghastrum fuscescens MO: PASQUET 1241
PL Sorghastrum nutans DEK: WYSOCKI S.N.
SG Trachypogon chevalieri MO: PASQUET 1190
PL Homozeugos eylesii INSDC: MT610079
PL Apluda mutica USDA−GRIN: PI 219568
353 Apluda mutica CNS: WATERHOUSE, B.M. 7832
PL Eulalia aurea USDA−GRIN: PI 12153
353 Eulalia aurea AD: C.J. BRODIE ET AL. 1567
APLUDINAE
SG Sorghum halepense GYPSUM9
PL Sorghum bicolor V3.1.1
SG Sorghum arundinaceum MO: AUB & BOOKOUT 106
PL Lasiorhachis perrieri TAN: O.P. NANJARISOA ET AL. 140
PL Lasiorhachis hildebrandtii INSDC: NC_036118
PL Cleistachne sorghoides MO: B. GOLDSMITH 12/71
SG Microstegium nudum K: M.S. VORONTSOVA 1917
353 Sarga plumosa AD: S. LEGGE 307
PL Sarga timorensis INSDC: NC_023800
SG Sarga versicolor INSDC: SRR427175
SORGHINAE
PL Miscanthus sacchariflorus INSDC: NC_028720
PL Miscanthus sinensis INSDC: NC_028721
SG Miscanthus oligostachyus B&T WORLD SEEDS: 530513
353 Miscanthus sinensis MEL: CLARKE, I.C. 4310
PL Saccharum officinarum INSDC: NC_035224
PL Saccharum spontaneum INSDC: NC_034802
SG Miscanthus nepalensis MO: ZHANG, ZHI−YING 13942
PL Eulalia siamensis BKF: TRAIPERM 557 (Apludinae)
PL Pseudosorghum fasciculare BKF: ARTHAN 067
SACCHARINAE
PL Germainia capitata INSDC: NC_035046
PL Pogonatherum paniceum INSDC: NC_029881
PL Imperata cylindrica DEK: BURKE 21
GERMANIINAE
PL Eulaliopsis binata INSDC: NC_035049
353 Eulaliopsis sykesii K: POLUNIN, O. 3170
PL Andropterum stolzii INSDC: MT610072
PL Dimeria ornithopoda BKF: TRAIPERM 575
353 Bhidea fischeri K: SALUMKE, C.B. 8294 (Andropogoninae)
SG Ischaemum byrone MO: PERLMAN 22941
ISCHAEMINAE
PL Eremochloa eriopoda INSDC: NC_035023
PL Eremochloa ciliaris MO: TRAIPERM 524
SG Eremochloa lanceolata MO: TRAIPERM 584
SG Eremochloa attenuata MO: TRAIPERM 162
PL Eremochloa ophiuroides ISC: L. CLARK ET AL. 1694
PL Rottboellia helferi BKF: TRAIPERM 574 (Rottboelliinae)
PL Mnesithea formosa INSDC: MT610073
SG Thaumastochloa striata MO: D.L. LEWIS 886
PL Hemarthria uncinata INSDC: MT610063
PL Hemarthria altissima INSDC: MT610054
353 Ophiuros exaltatus BRI: MCDONALD, K.R. 18636
PL Hackelochloa granularis INSDC: MT610093
353 Rottboellia rottboellioides K: BARRETT, M.D. 5002 (Rottboelliinae)
353 Thaumastochloa monilifera BRI: MCDONALD, K.R. 19460A
PL Glyphochloa forficulata K: P.M. JARRETT ET AL. HFP 896
353 Hackelochloa granularis MEL: COWIE, I.D. 8570
SG Rottboellia afraurita MO: PASQUET 1156 (Rottboelliinae)
SG Miscanthus sinensis RSA: P. ZIKA 25806 (Saccharinae)
RATZEBURGIINAE
PL Microstegium vimineum INSDC: MT610045
SG Microstegium fasciculatum MO: CF VAN BEUSEKOM ET AL 4092
SG Microstegium geniculatum MO: HSIU−LAN HO 920
PL Kerriochloa siamensis BKF: TRAIPERM 580
PL Sehima nervosa INSDC: MT610076
SG Elionurus euchaetus VOUCHER N/A
SG Elionurus elegans MO: C. VANDER BERGHEN 7324
PL Coix aquatica INSDC: MT942628
PL Coix lacryma−jobi BKF: ARTHAN 072
353 Phacelurus speciosus K: SIDDIQUI, M.A. 26726
PL Rottboellia cochinchinensis ISC: CLARK ET AL. 1698
353 Rottboellia cochinchinensis MEL: JAGO, R.L. 7280
ROTTBOELLIINAE
PL Ischaemum afrum USDA−GRIN: 364924 (Ischaeminae)
PL Chrysopogon serrulatus USDA−GRIN: PI 219580
SG Chrysopogon aucheri MO: ROBERT GIBBONS 0308
PL Chrysopogon aciculatus FDS: Z16053
353 Chrysopogon latifolius TCD: HODKINSON, T.R. 9256
PL Chrysopogon zizanioides MO: KELLOGG VET−MRL−001
SG Clausospicula extensa MO: G.J. LEACH & C.R. DUNLOP 2575
CHRYSOPOGONINAE
PL Andropogon burmanicus BKF: ARTHAN 071 (Andropogoninae)
PL Pseudopogonatherum contortum INSDC: NC_035026
PL Parahyparrhenia siamensis INSDC: NC_035033
PL Eriochrysis cf. cayennensis BKF: WELKER 365
PL Urelytrum agropyroides INSDC: MT610050
PL Oxyrhachis gracillima INSDC: MT610057
RHYTACHNINAE
SG Phacelurus franksiae MO: PASQUET 1257
353 Phacelurus gabonensis K: WIERINGA, J.J. 3903
PL Polytoca digitata BKF: ARTHAN 054
353 Polytoca cyathopoda TCD: JACOBS, S. 9289
CHIONACHNINAE
PL Zea perennis USDA−GRIN: AMES 21874
PL Zea luxurians USDA−GRIN: PI 441933
PL Zea diploperennis INSDC: NC_030377
PL Zea mays INSDC: NC_001666
SG Tripsacum latifolium MO: W.E. HARMON, J.A. FUENTES 2183
PL Tripsacum australe INSDC: MT610096
PL Tripsacum dactyloides INSDC: NC_037087
TRIPSACINAE
SG Arthraxon antsirabensis K NOP 217
PL Arthraxon hispidus BRI: P.I. FORSTER PIF45419
PL Arthraxon lancifolius AAU: S. LAEGAARD 21760
PL Arthraxon prionodes USDA−GRIN: PI 659331
SG Arthraxon meeboldii MO: SANTAPAU SJ 10033
PL Arthraxon microphyllus BKF: TRAIPERM 537
ARTHRAXONINAE
PL Lasiurus scindicus K: A. NAEGELI DJI/78−26
ANDROPOGONEAE

**Arundinelleae**
*(plastome)*
0.001 substitutions/site

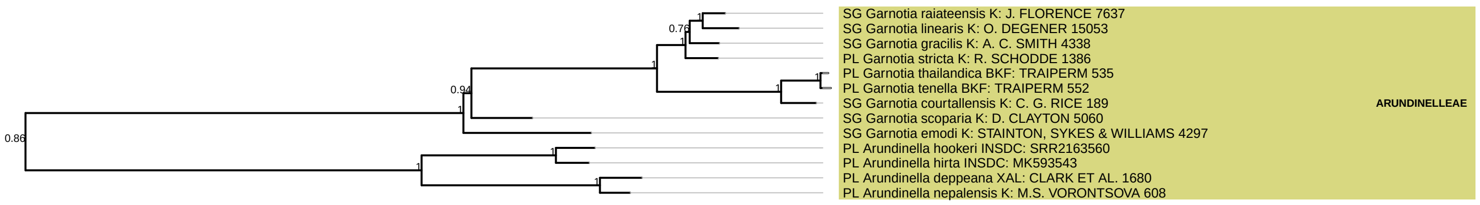

**Paspaleae**
*(plastome)*

0.001 substitutions/site

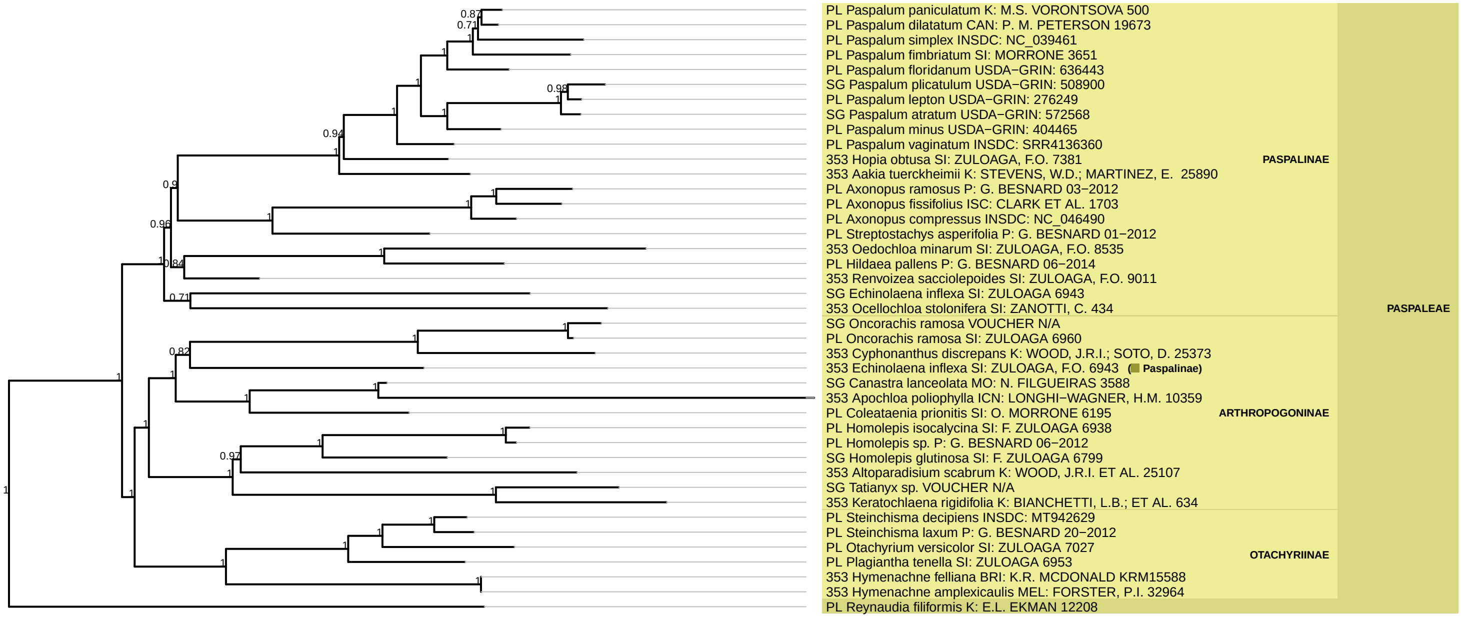

**Paniceae**
*(plastome)*

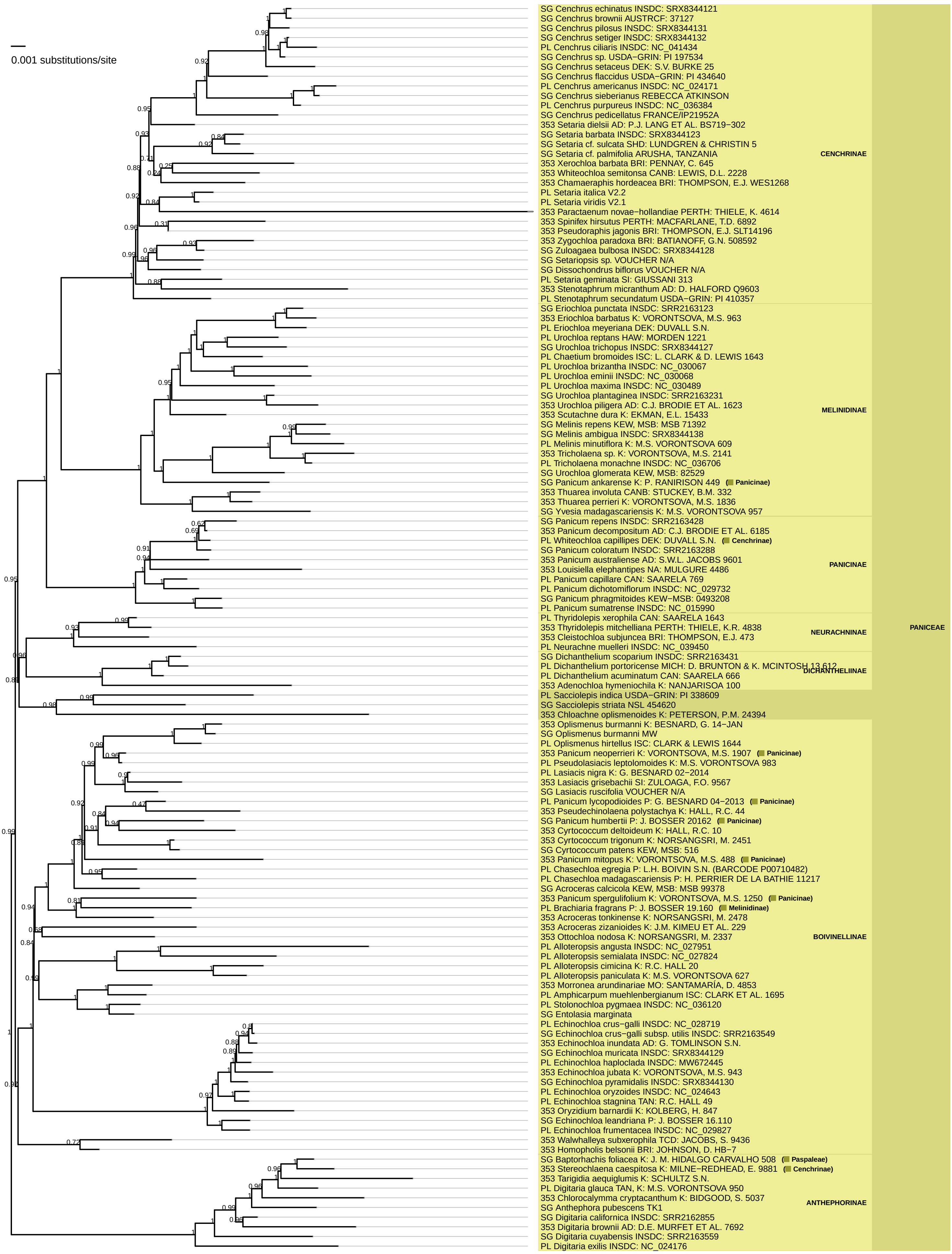

**Tristachyideae**
*(plastome)*

0.001 substitutions/site

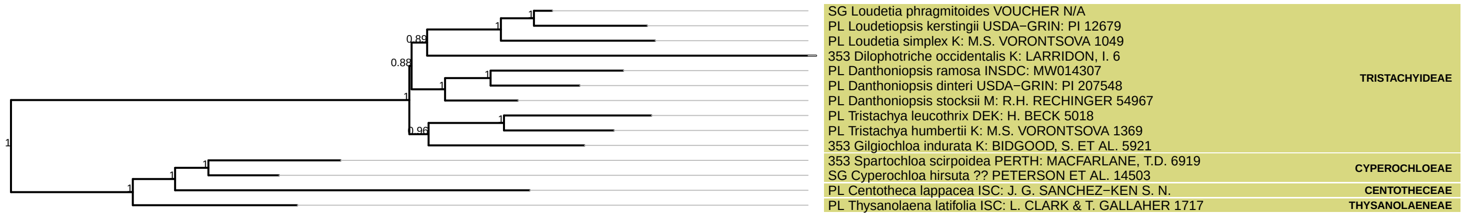

**Cynodonteae**
*(plastome)*

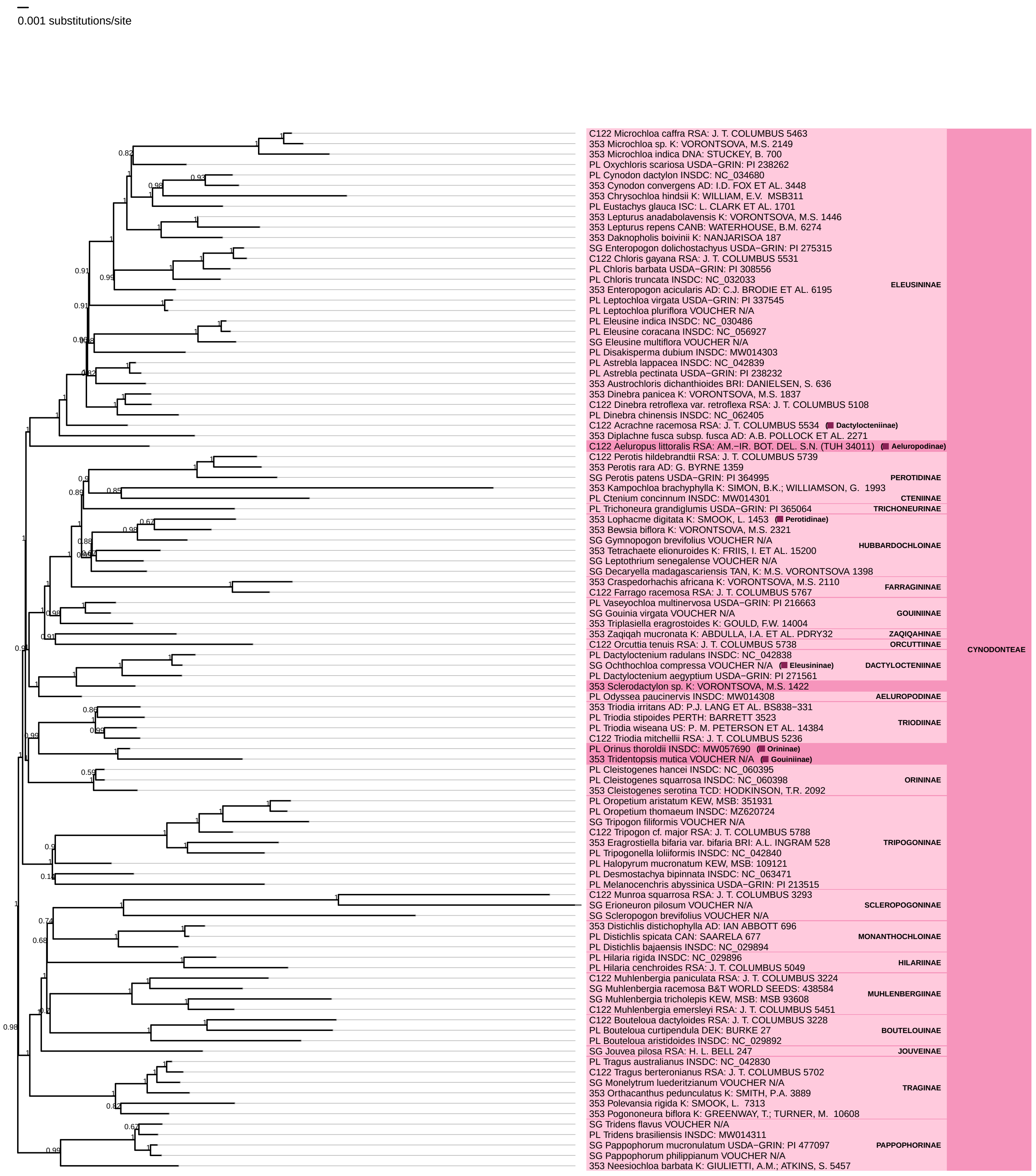

**Zoysieae**

*(plastome)*

0.001 substitutions/site

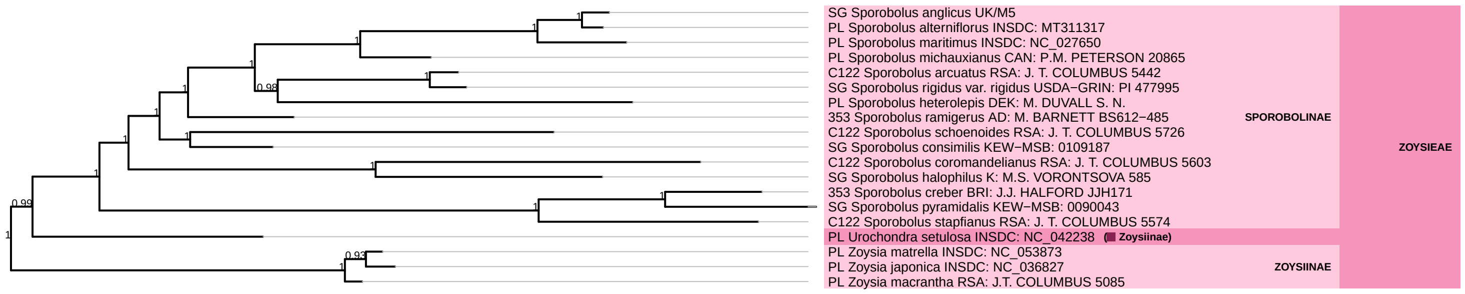

### **Eragrostideae**

*(plastome)*

0.001 substitutions/site

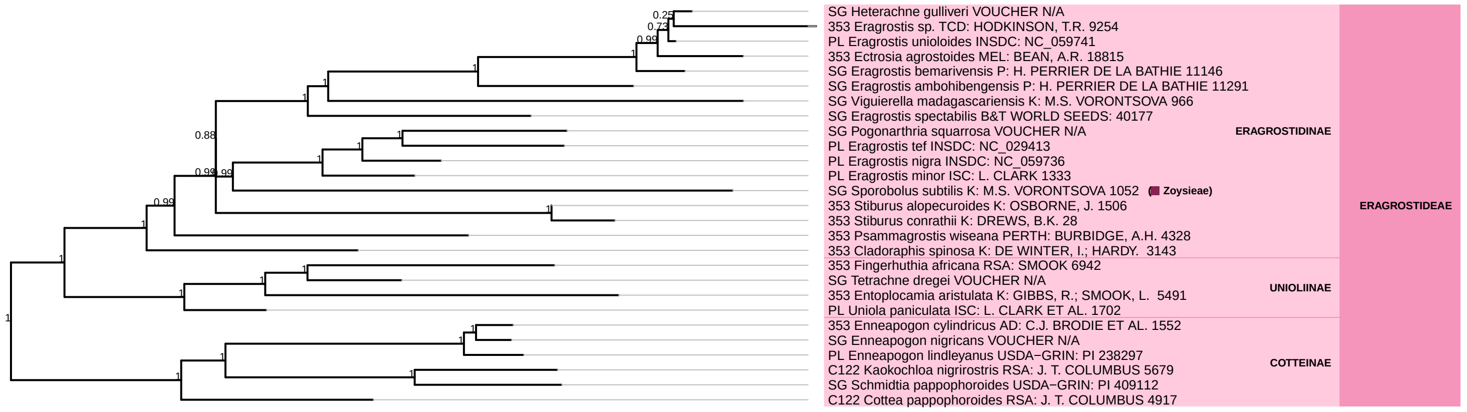

**Danthonioideae**
*(plastome)*

0.001 substitutions/site

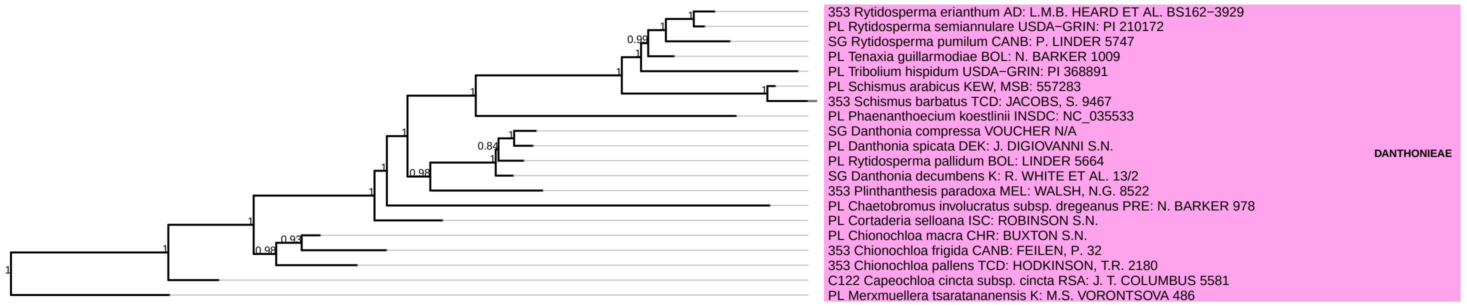

**Arundinoideae**
*(plastome)*
0.001 substitutions/site

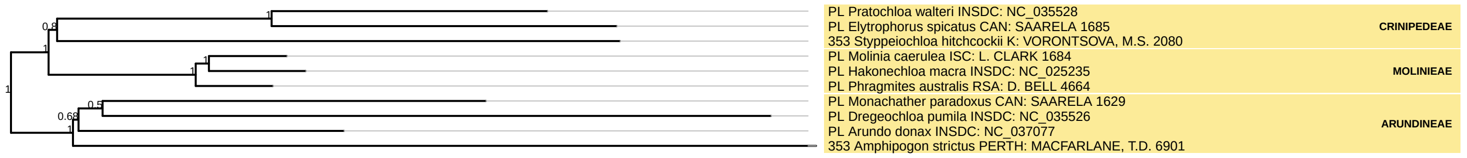

### Micrairoideae

*(plastome)*

0.001 substitutions/site

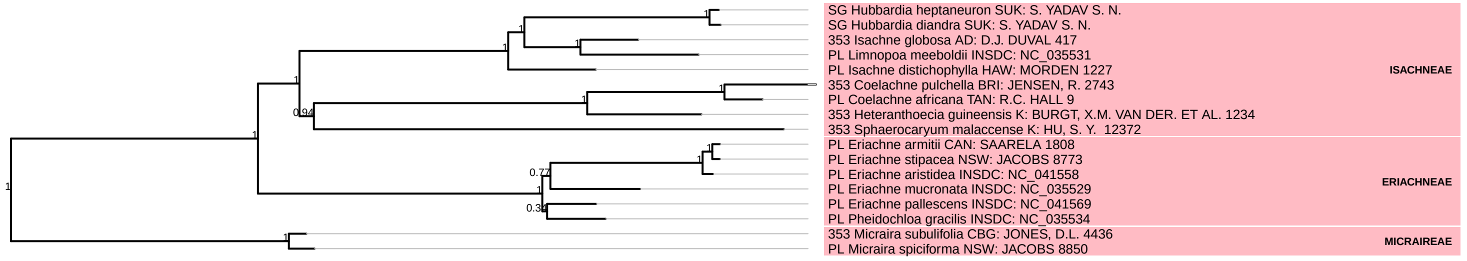

### Aristidoideae
*(plastome)*

0.001 substitutions/site

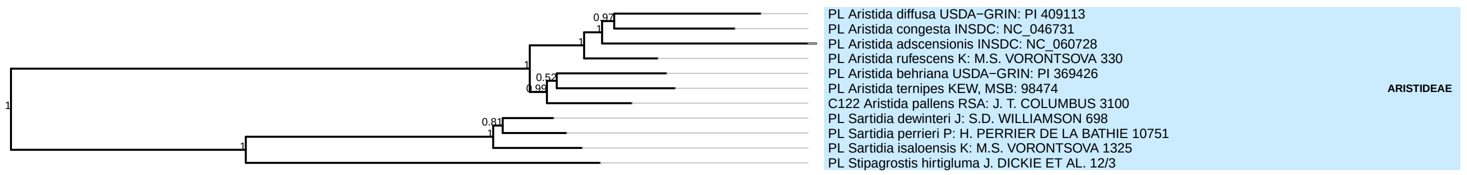

**Poeae**
*(plastome)*

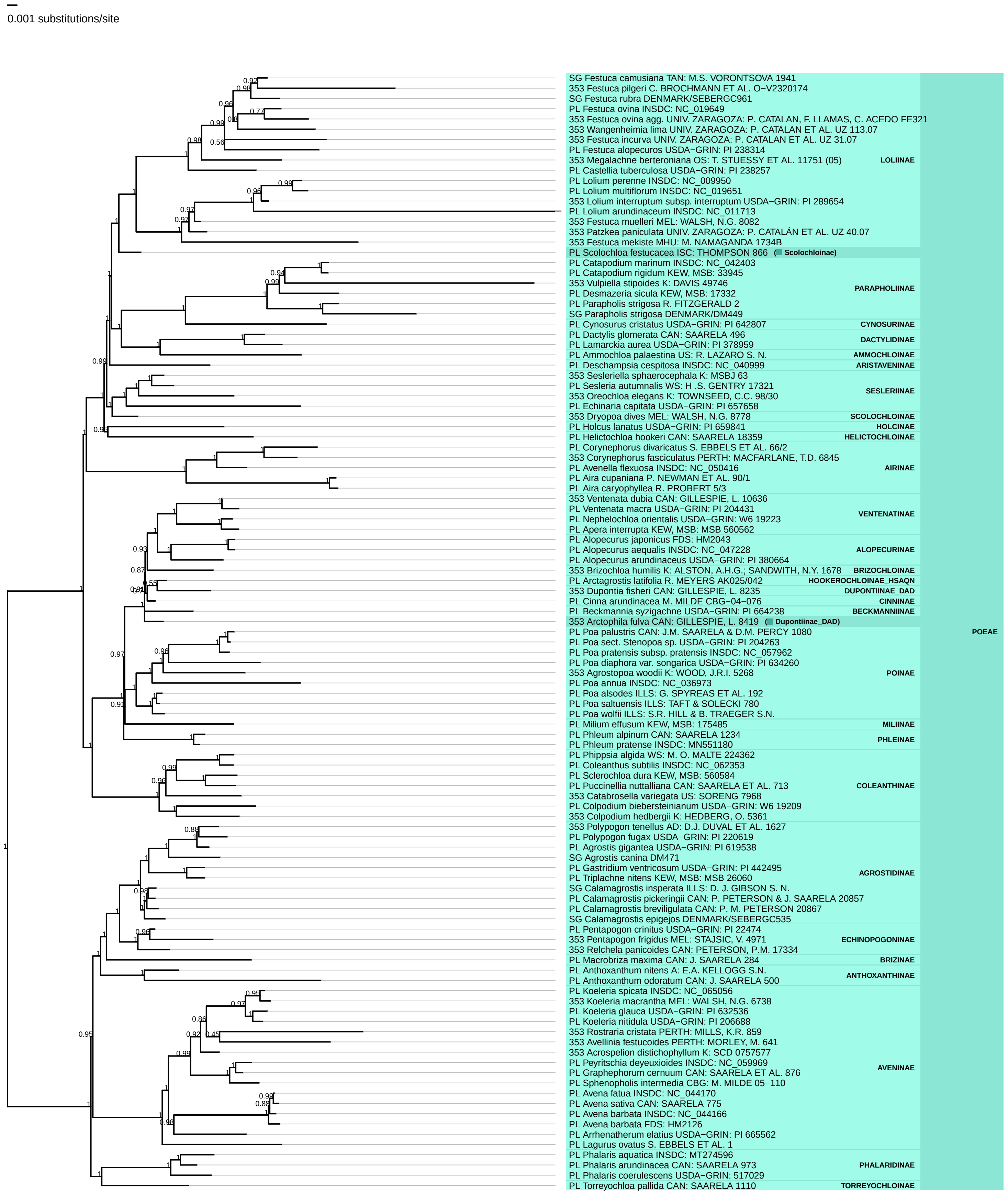

**Triticeae**
*(plastome)*

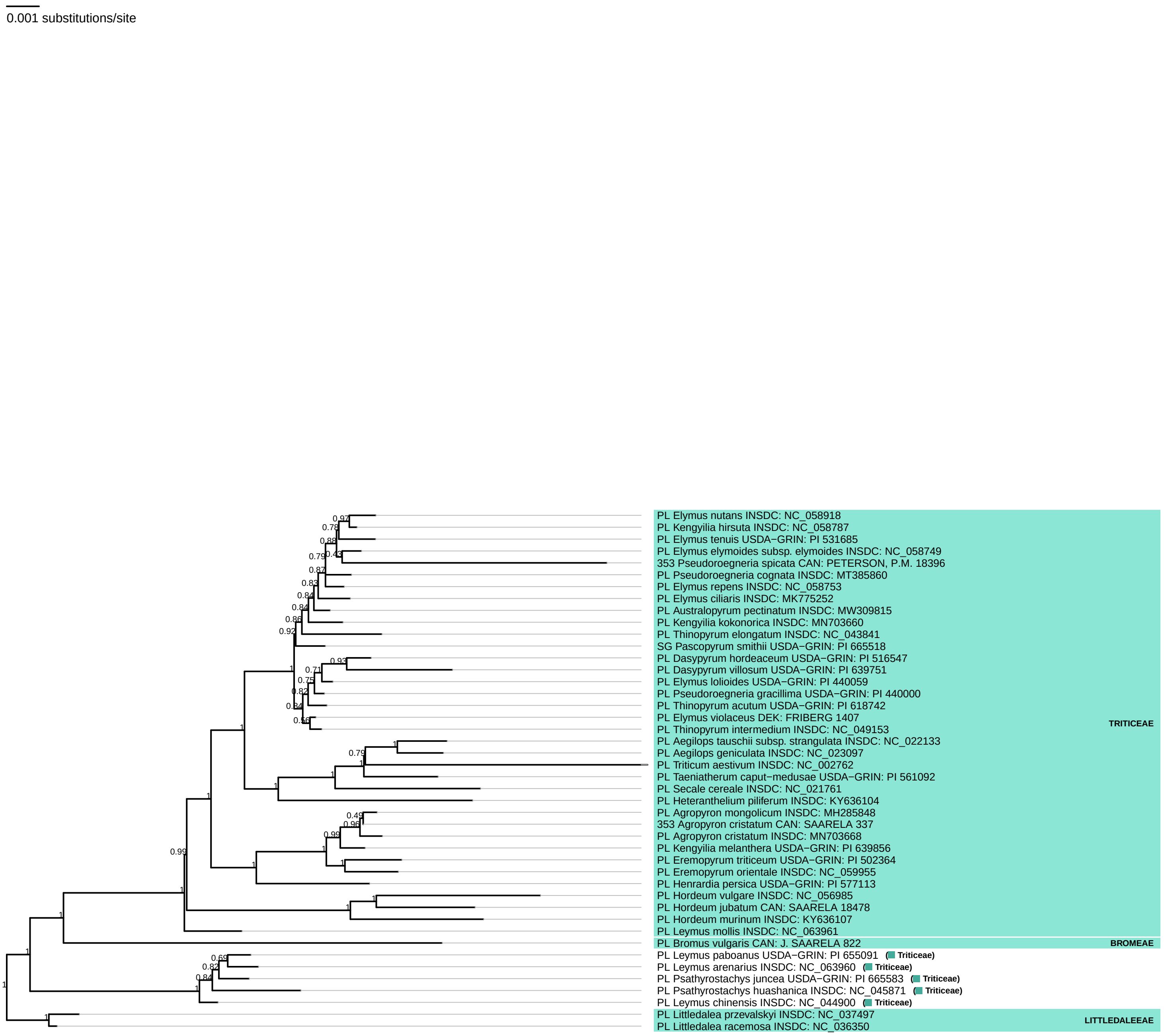

**Stipeae**
*(plastome)*

0.001 substitutions/site

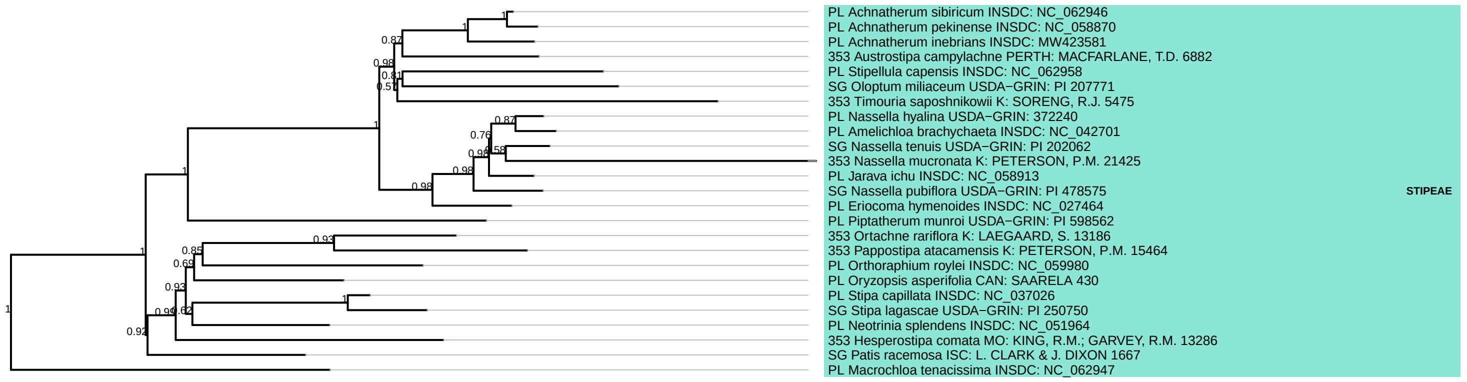

**Bambusoideae**
*(plastome)*

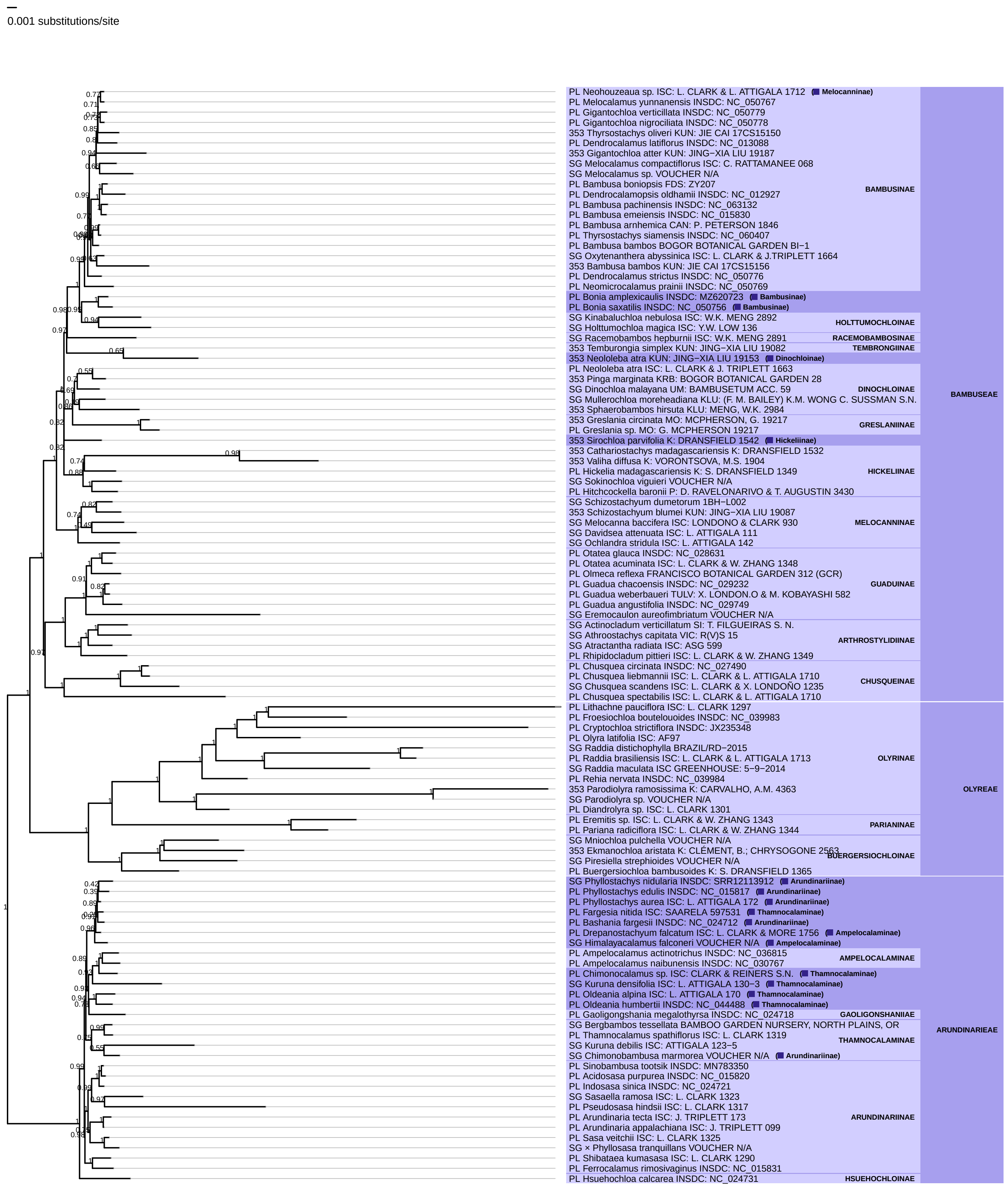

**Oryzoideae**
*(plastome)*

0.001 substitutions/site

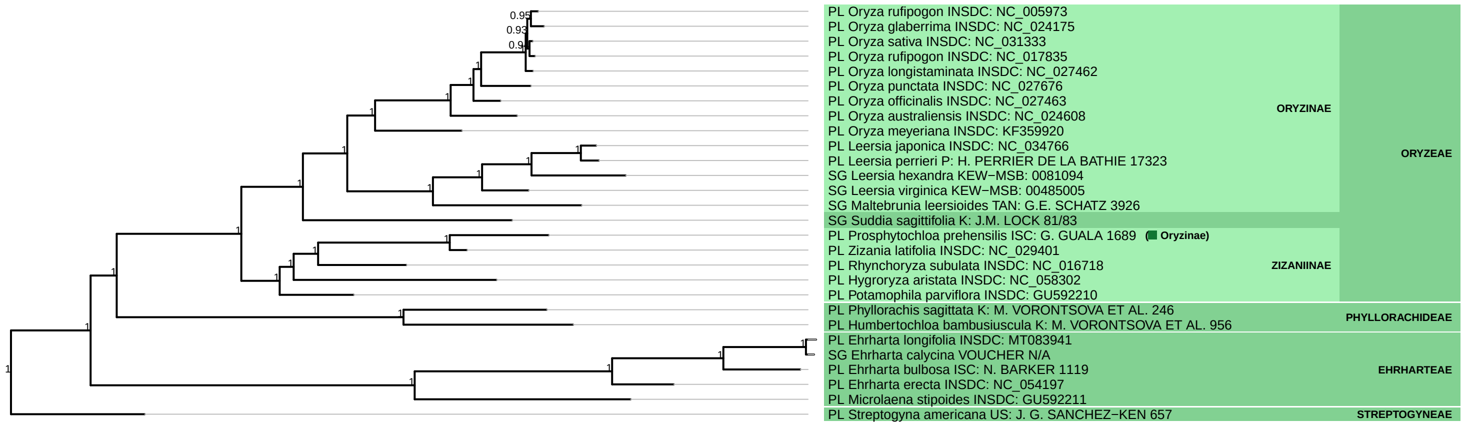
